# Discordant transcriptional signatures of mitochondrial genes in Parkinson’s disease human myeloid cells

**DOI:** 10.1101/2020.07.20.212407

**Authors:** Elisa Navarro, Evan Udine, Katia de Paiva Lopes, Madison Parks, Giulietta Riboldi, Brian M. Schilder, Jack Humphrey, Gijsje J. L. Snijders, Ricardo A. Vialle, Maojuan Zhuang, Tamjeed Sikder, Charalambos Argyrou, Amanda Allan, Michael Chao, Kurt Farrell, Brooklyn Henderson, Sarah Simon, Deborah Raymond, Sonya Elango, Roberto A. Ortega, Vicki Shanker, Matthew Swan, Carolyn W. Zhu, Ritesh Ramdhani, Ruth H. Walker, Winona Tse, Mary Sano, Ana C. Pereira, Tim Ahfeldt, Alison M. Goate, Susan Bressman, John F. Crary, Lotje de Witte, Steven Frucht, Rachel Saunders-Pullman, Towfique Raj

## Abstract

An increasing number of identified Parkinson’s disease (PD) risk loci contain genes highly expressed in innate immune cells, yet their potential role in pathological mechanisms is not obvious. We have generated transcriptomic profiles of CD14^+^ monocytes from 230 individuals with sporadic PD and age-matched healthy subjects. We identified dysregulation of genes involved in mitochondrial and proteasomal function. We also generated transcriptomic profiles of primary microglia from autopsied brains of 55 PD and control subjects and observed discordant transcriptomic signatures of mitochondrial genes in PD monocytes and microglia. We further identified PD susceptibility genes, whose expression, relative to each risk allele, is altered in monocytes. These findings reveal that transcriptomic mitochondrial alterations are detectable in PD monocytes and are distinct from brain microglia, and facilitates efforts to understand the roles of myeloid cells in PD.

## Introduction

Parkinson’s disease (PD) is a progressive neurodegenerative disorder of aging that affects motor, cognitive and other functions (*1*). Several lines of evidence suggest that the immune system plays an important role in PD, however the mechanisms underlying immune dysfunction are largely unknown. Recent genetic studies have identified over 78 PD risk loci (*2*), and many of these loci contain genes involved in immune function. Genomic analysis has demonstrated that PD-associated susceptibility alleles alter the expression of nearby genes in peripheral monocytes (*3–5*) and significant enrichment of PD-heritability in gene sets highly expressed in microglia (*6*). Healthy microglia are essential for clearing of debris, such as α-synuclein (*7, 8*) and for maintaining brain homeostasis. α-synuclein can activate microglia releasing neurotoxic factors that may lead to the death of dopaminergic neurons (*9–12*). Peripheral monocytes from PD patients have been shown to be hyperactive in response to α**-**synuclein stimulation (*13*). Monocytes have also been found to be capable of entering and interacting with the central nervous system (CNS) via the meninges (*14–16*) and may be involved in the phagocytosis of protein aggregates of debris from degenerating neurons (*17, 18*). The Braak hypothesis, which proposes that α-synuclein pathology starts in the periphery (*19*), and the gut-origin hypothesis of PD (*20–22*) also postulate that peripheral immune cells might be exposed to PD-pathology early during the disease. Collectively, these studies support the importance of non-neuronal cell types including peripheral immune cells and brain resident glial cells in PD pathophysiology. However, there are critical gaps in our understanding of how these cells contribute to PD, in part due to the challenge of accessing patient-derived samples, and while some studies have characterized monocytes or microglia in PD (*23, 24*), they were limited in sample size.

The Myeloid cells in Neurodegenerative Diseases (MyND) initiative is a collaborative effort with the goal of creating a multi-omic atlas of myeloid cells from the periphery and from autopsied brains of subjects with PD, Alzheimer’s disease (AD), and age-matched controls. This study reports the first phase of this initiative, which profiles the transcriptome of CD14^+^ monocytes and microglia from PD subjects and age-matched controls. As a source of patient tissue in which to study early disease processes, blood samples are easily accessible and cost**-**efficient and can be obtained with minimal risk to the patients. Peripheral blood cells such as monocytes perform many of the fundamental cellular processes that are perturbed in PD (*25*), and we hypothesize that they may recapitulate some of the cellular pathology observed in the PD brain. Here, we performed large-scale, unbiased, systematic analysis incorporating genomic, and bulk and single-cell transcriptomic data to identify genes and co-expression networks that are dysregulated in PD myeloid cells. We further performed expression quantitative trait loci (eQTL) analysis of monocytes to identify colocalization between alleles driving variation in mRNA abundance and PD susceptibility. Overall, our data demonstrate mitochondrial and proteasomal transcriptome alterations in PD monocytes, with mitochondrial genes discordantly expressed in PD monocytes and microglia.

## Results

### Participant recruitment and sample collection

Participants have been recruited from five clinical sites in New York City (see Methods). For each participant we have isolated peripheral blood mononuclear cells (PBMCs); of these isolated PBMCs, 5 million were sorted to CD14^+^ monocytes while the rest were cryopreserved for future studies. We also have banked whole blood for DNA isolation and plasma for biomarker discovery (**Fig. 1**). For this study, we used data from 230 participants, including 135 with a diagnosis of idiopathic PD (“cases”) and 95 age-matched participants (“controls”) with no reported neurological or auto-immune diseases. Participants have a mean age of 67 years old. The sex is balanced overall but within the PD group, 36% are women as it is more common in men (**Fig. S1B, Table S1**). The average age of onset (considered as age of diagnosis) in the PD group is 57.3 years old, with a disease duration of 8.3 years and Hoehn & Yahr (H&R) (*26*) scale of 1.8 (see additional clinical information in **Fig. S1C**).

**Figure 1.**
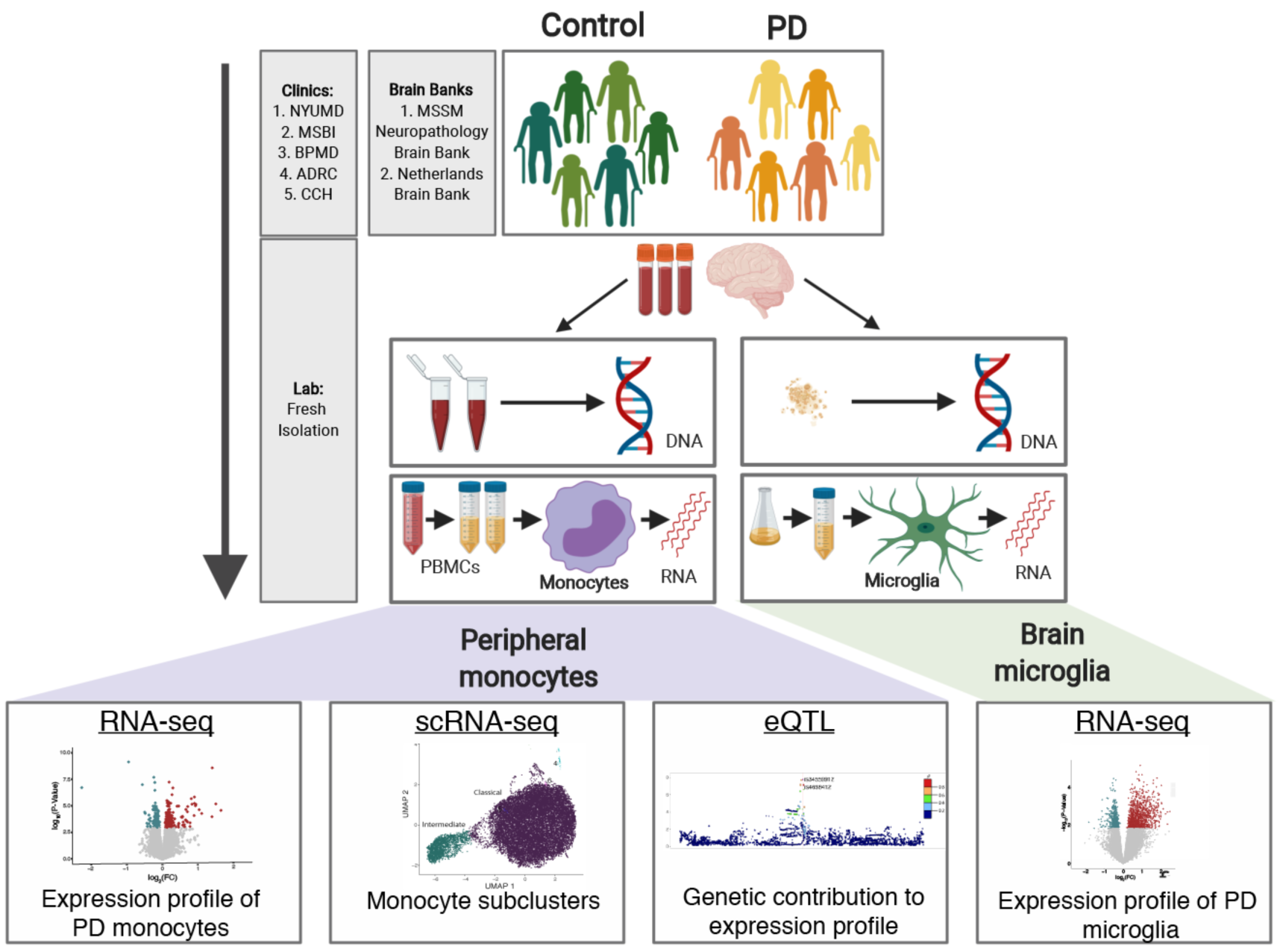
Overview of the study design. Parkinson’s disease and age-matched control subjects were recruited from five clinical sites: Movement Disorder Center at Mount Sinai Beth Israel (MSBI), Bendheim Parkinson and Movement Disorders Center at Mount Sinai (BPMD), Fresco Institute for Parkinson’s and Movement Disorders at New York University (NYUMD), and the Alzheimer’s Research Center (ADRC) and Center for Cognitive Health (CCH) at Mount Sinai Hospital. Fresh blood samples from PD and age-matched healthy subjects were collected following a rigorous, standardized set of procedures and used to isolate peripheral blood mononuclear cells (PBMCs). From the PBMCs, CD14^+^ monocytes were isolated using magnetic beads. Primary microglia were isolated from autopsied brains from two brain banks: Netherlands Brain Bank (NBB) and the Neuropathology Brain Bank and Research CoRE at Mount Sinai Hospital. Primary human microglia were isolated using CD11b^+^ beads. mRNAs from these cells were profiled using RNA-seq and single-cell RNA-Seq. Genome-wide genotyping was performed using DNA isolated from these samples. The data generated enabled to (from left to right) (i) description of the transcriptomic profiling of PD-monocytes at the gene, transcript and splicing levels, (ii) understanding of the contribution of the different monocyte subpopulation to the disease, (iii) integration of genomic and expression data to identify monocyte eQTLs and (iv) comparison of the transcriptome signatures of PD peripheral monocytes to CNS microglia.

Primary microglia have been isolated from postmortem brain tissues from the Netherlands Brain Bank and the Neuropathology Brain Bank & Research Core at the Icahn School of Medicine at Mount Sinai Hospital, New York. For this study, microglia from up to six different brain regions from 13 PD donors (22 samples) and 42 age-matched control donors (106 samples) have been used for RNA sequencing and downstream analysis (**Table S2**). The average age of death is 80.22 years old and 78.5 years old for control and PD cases, respectively. Disease duration in the PD group is 13.5 years, and sex is balanced. Further details on the donors and samples used in this study are in the Methods.

### Peripheral monocytes of PD patients show mitochondrial and proteasomal alterations

We isolated human fresh monocytes from patient-derived blood using CD14^+^ beads. After rigorous quality control, we retained RNA-sequencing (RNA-seq) data from monocytes of 230 subjects for all downstream analyses. RNA-seq data were normalized and corrected to account for the effect of known biological and technical covariates (see methods; **Fig. S2-S5**). RNA-seq based quantifications enabled assessment of coding and non-coding differential gene expression, differential isoform expression, and differential splicing analyses (**Fig. S2**). A total of 300 differentially expressed genes (DEGs) were identified when comparing PD-derived monocytes to controls (False Discovery Rate [FDR] < 0.05). Of these, 162 identified DEGs were upregulated while 138 DEGs were downregulated (**Fig. 2A, Table S3**). The effect sizes for most of the DEGs were small (|log_2_ fold change (FC)| < 0.5). The DEGs were not driven by *LRRK2* or *GBA* mutation carriers, or by individuals of Ashkenazi Jewish (AJ) ancestry (**Fig. S4C, D**). Additionally, sex contributed small proportion of total variation in gene expression (mean variance 0.25% and standard deviation of 2.1%) (**Fig. S4A**). As the majority of PD cases were taking dopaminergic medication, Levodopa (L-dopa), we tested if gene expression was correlated with L-dopa equivalent daily dose (LEDD) on a subset of individuals (n = 110). We found no significant correlation for any genes at FDR < 0.05, and of the DEGs, only four genes were significant at a nominal *P-value* < 0.05 threshold (**Fig. S4E**).

**Figure 2.**
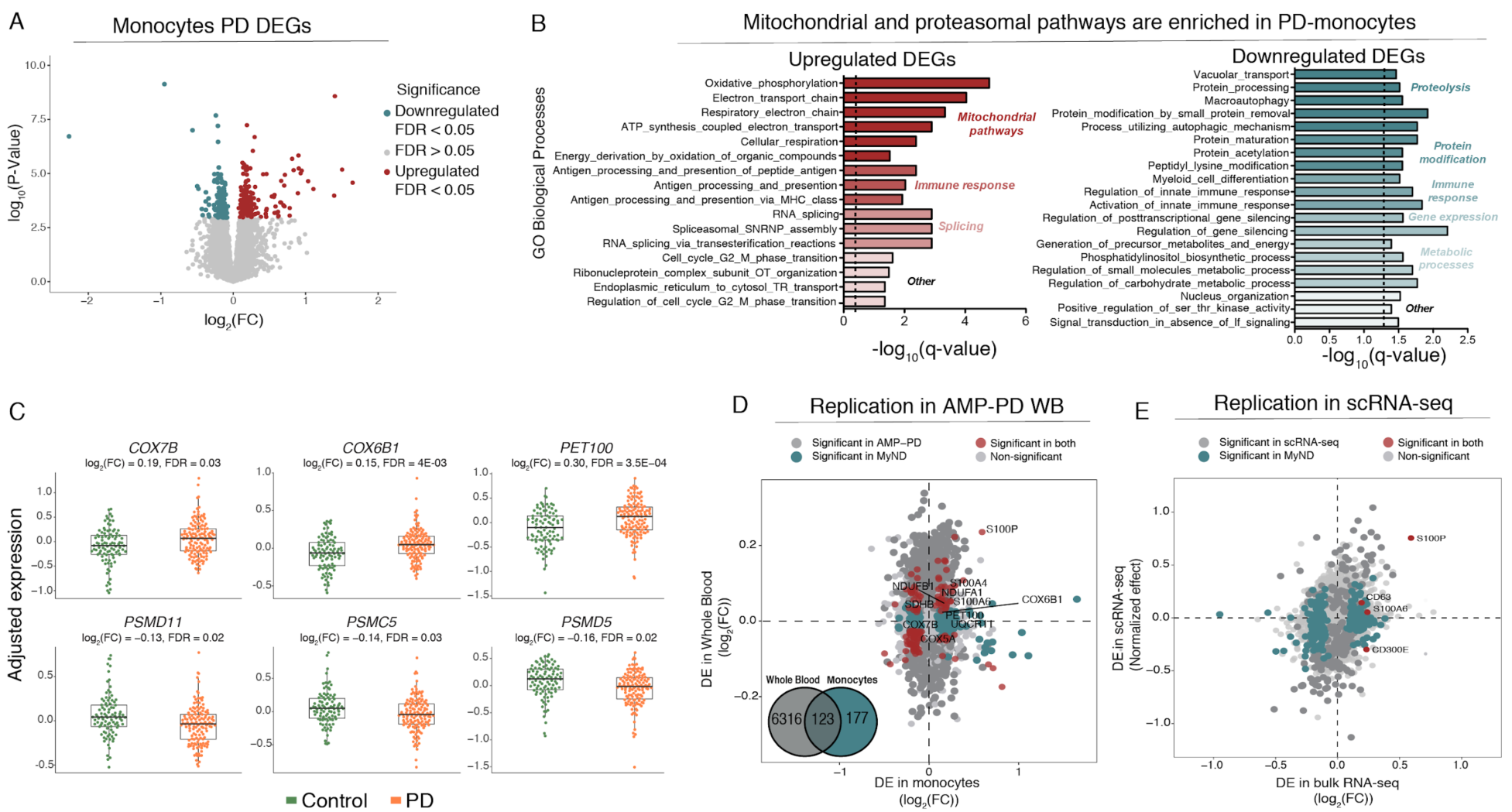
Transcriptomic analysis of PD-derived monocytes and age-matched controls. **(A)** Volcano plot showing the fold-change (FC) of genes (log_2_ scale) between PD-monocytes (n = 135) and controls (n = 95) (x-axis) and their significance (y-axis, -log_10_ scale). DEGs at FDR < 0.05 are highlighted in red (upregulated genes) and blue (downregulated genes). **(B)** Pathway analysis for the upregulated (left panel) and downregulated (right panel) DEGs. Significance is represented in the x-axis (-log_10_ scale of the q-value). Only the 20 most significant pathways (FDR q-value < 0.05) with a minimum of 5 genes overlap are shown. Pathways are grouped and colored by biologically-related processes. **(C)** Examples of selected mitochondrial (top panel) and proteasomal (bottom panel) DEGs. Adjusted expression of the voom normalized counts after regressing covariates is shown. **(D)** Fold-change (log_2_ scale) correlation of DEGs between MyND monocytes (x-axis) and AMP-PD whole blood (y-axis). Bottom left: Venn-diagram showing the overlap of significant genes in whole blood differential expression analysis from AMP-PD (FDR < 0.05) and monocytes from MyND (FDR < 0.05). Genes are colored by significance, considering significant DEGs at FDR < 0.05. **(E)** Fold-change (log_2_ scale) correlation of DEGs between bulk monocytes (x-axis) and single-cell across-clusters analysis (y-axis). Four outlier genes were removed for easier visualization. Genes are colored by significance, considering significant DEGs at q-value < 0.05.

We performed gene set enrichment analysis (GSEA) to evaluate which biological processes and molecular functions were enriched for DEGs. The upregulated DEGs were significantly enriched for a number of Gene Ontology (GO) biological processes (BP) including mitochondrial function, immune response, and RNA splicing (**Fig. 2B, Table S4**). The most significant BP were related to mitochondrial oxidative phosphorylation (OXPHOS), which includes essential components of respiratory chain complexes such as NADH dehydrogenase (*NDUFA1* and *B1*) and Cytochrome C Oxidase (*COX5A, 6B1, 7A2*, and *7B)* (FDR q-value < 0.05) (**Fig. 2B, C, Table S4**). Using a curated mitochondrial gene list (*27*), we found significant enrichment for OXPHOS genes (*P-value* = 0.00015, Fisher’s exact test), but not for other mitochondrial processes such as dynamics or mito-nuclear crosstalk (*P-value* = 0.80, Fisher’s exact test) and quality control (*P-value* = 0.66, Fisher’s exact test). The downregulated DEGs were overrepresented for functions including proteolysis, protein modification, immune response, and metabolic processes (FDR q-value < 0.05). Some of the genes in the proteolysis process are involved in proteasomal structure (*PSMC5, PSMD5*, and *PSMD11*) (**Fig. 2B, C**), ubiquitin (*USP10*), and autophagy-related function (*GSK3B, PIK3R4, STAM*). While most DEGs were part of a coherent biological function, we identified many that were not part of any processes including members encoding for the S100 proteins (*S100A4, S100A6, S100P*), which play an important role in inflammatory responses and function as damage-associated molecular pattern (DAMP) molecules (*28, 29*). The S100 proteins have been shown to be upregulated in the substantia nigra and cerebrospinal fluid (CSF) of patients with PD as well as in a mouse model following MPTP (1-methyl-4-phenyl-1,2,3,6-tetrahydropyridine), a toxin that causes parkinsonism in treated mice (*30*).

We next expanded these analyses to isoform transcript-level and local splicing (using intronic excision ratios) to identify transcriptomic dysregulation due to alternative splicing. We observed 1020 differentially expressed transcripts (DETs) and 161 differential splicing events (DS) at FDR < 0.05, corresponding to 939 and 158 unique genes, respectively (**Fig. S6, S7, Table S5, S6**). With the exception of mitochondrial function, the pathway analysis of DET and DS identified the same biological processes as the DEGs and expanded the list of genes involved in the protein degradation machinery including autophagy-related, proteasome, and lysosomal functions (**Fig. S6 and S7; Table S7, S8**). These include the transmembrane protein 175 (*TMEM175*), a lysosomal K^+^ channel, a gene in a PD GWAS locus that has been shown to play a critical role in lysosomal and mitochondrial function and PD pathogenesis (*31*). Also, two members (*MTOR* and *RICTOR*) of the rapamycin (mTOR) signaling pathway, a central regulator of the autophagy process (*32*), were identified in the DS analysis. Together, these results highlight key genes involved in protein degradation machinery have aberrant RNA splicing in PD monocytes.

With respect to the reproducibility of our results, we have performed two separate analyses to replicate our findings. First, we incorporated whole blood (WB) transcriptomic data from 780 PD cases and 504 controls from the Parkinson’s Progression Markers Initiative (PPMI) (one of the cohorts of the Accelerating Medicines Partnership: Parkinson’s Disease [AMP-PD]). Although not a direct replication since the AMP-PD transcriptome is from WB but given the large sample size, we expected to capture some of the monocyte-specific effects in blood. After QC (**Fig. S8**), we found 6439 DEGs at FDR < 0.05 (**Table S9**). Of 300 monocyte DEGs, 123 were also significant DEGs at FDR < 0.05. For the majority (6 out of 8 genes) of the mitochondrial DEGs in monocytes, we observed a concordant direction of effect in WB (**Fig. 2D, 6C**). However, the effect size in AMP-PD WB was weaker than in monocytes (mean FC = 0.09 in WB; mean FC = 0.27 in monocytes; for genes with FDR < 0.05; *P-value* < 2×10^−16^, independent-sample t-test) despite the large sample size (n = 1284) in AMP-PD. These results demonstrate the improved power of purified cell populations over mixtures of cell types such as whole blood, which may result in failure to properly capture the activity of cell-type-specific effects. Finally, we also validated our bulk RNA-seq findings in single-cell RNA-seq (scRNA-seq) of CD14^+^ monocytes by multiplexing 10 independent monocytes samples (seven PD, three controls; see below for further details). The effect size (normalized effect) (*33*) of DEGs from across-clusters gene expression from scRNA-seq were highly correlated with effect size from bulk RNA-seq DEGs (Spearman ρ =0.59, *P-value* = 5.04×10^−6^, Spearman rank correlation) (**Fig. 2E, Table S10**). For example, *S100P* and *S100A6* were significant in both datasets (adjusted *P-value* < 0.05), and the majority (9 of 11) of the other members of the *S100* gene family shared the same directionality in both datasets.

To place the transcriptome changes in a systems-level framework, we performed Weighted Gene Co-expression Network Analysis (WGCNA). Network analysis partitions the monocyte transcriptome into modules of co-expressed genes linked to specific biological processes and pathways. We identified 65 modules of strongly co-expressed groups of genes, ranging from *turquoise* (largest, 2541 genes) to *orangered3* (smallest, 30 genes) (**Fig. S9, Table S11**). Of these, six modules were enriched for DEGs including *sienna3* and *green*, which are highly enriched for mitochondrial function with upregulated DEGs, and the *greenyellow* module enriched for genes in proteasomal and lysosomal functions with downregulated DEGs (**Fig. 3A**). Another example is the *black* module, containing 41 proteasomal and 35 ubiquitin-related genes, which is also enriched for downregulated DEGs and genes in PD GWAS loci (such as *ATP6AP2, ITPKB*, and *KPNA1*). We used LD score regression (LDSC) (*34*) to partition PD GWAS heritability into bins of correlated SNPs located within genes from each module. We found that 16 out of 65 modules showed enrichment for PD heritability (FDR < 0.05) including *green, salmon*, and *red* modules involving mitochondrial, lysosomal, and immune function, respectively (**Fig. 3A**). Next, we correlated the module eigengene, the first principal component (PC) of the module gene expression level, with PD diagnosis. We observed three modules that were significantly correlated with PD diagnosis (FDR < 0.05, Wilcoxon signed-rank test), including *turquoise* (ubiquitin-related activity), *antiquewhite4* (proteasome), and *darkseagreen4* (**Fig. S10B**). Given that multiple modules are associated with mitochondrial or lysosomal function, we considered taking the eigengene of all genes in either mitochondrial (n = 1302) or lysosomal (n = 526) GO categories. We found that the mitochondrial and lysosomal eigengenes were significantly upregulated and downregulated in PD, respectively (**Fig. 3B, C**). Taken together, these results illustrate that several co-expression gene modules in monocytes are enriched for PD heritability and further suggest subtle disruption of gene expression in specific biological networks including those with mitochondrial and proteo-lysosomal function.

**Figure 3.**
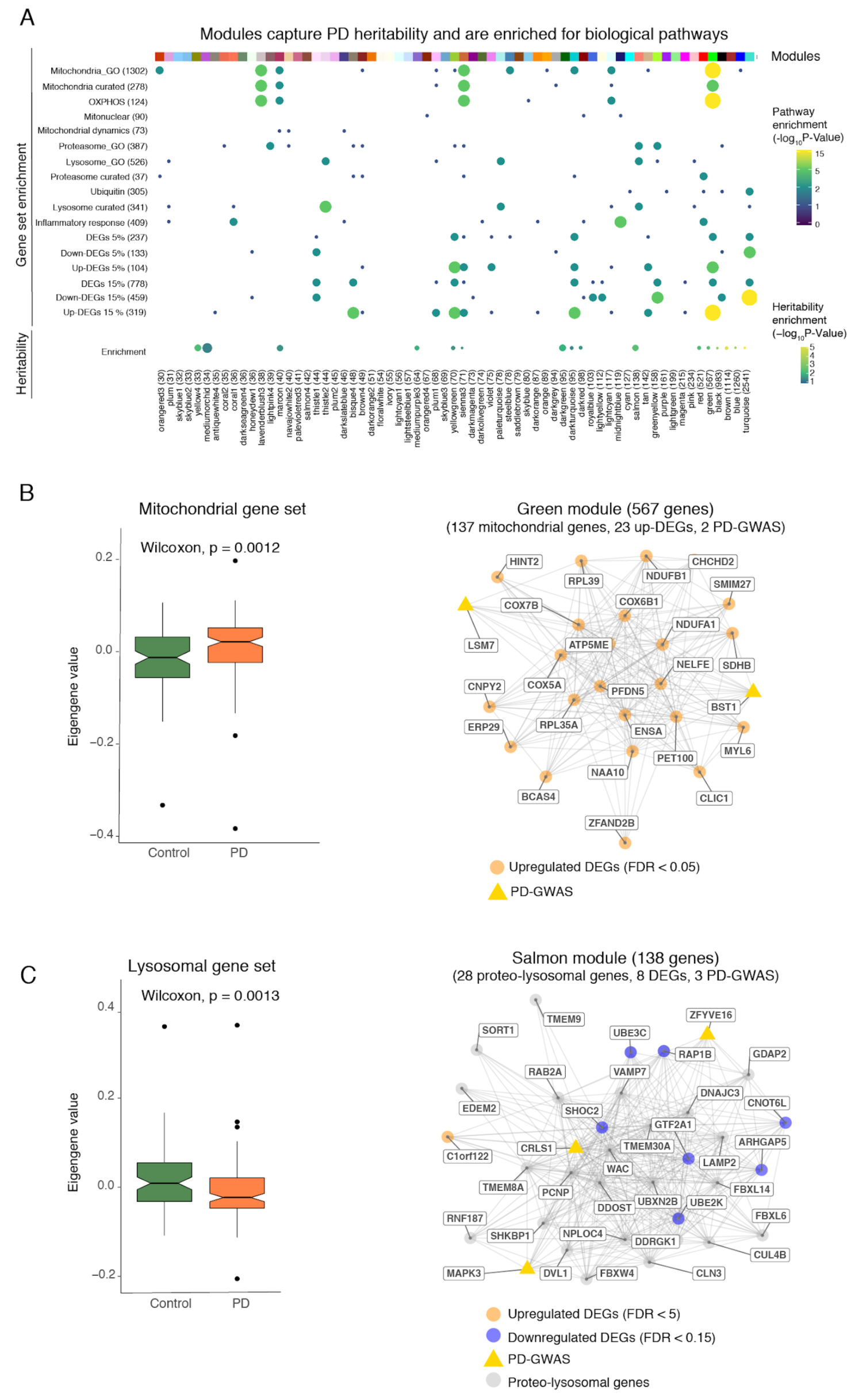
Co-expression networks in monocytes capture PD-specific processes. **(A)** Enrichment of modules (x-axis) containing co-expressed genes for specific biological pathways and curated gene sets (y-axis). Modules are represented by color names and are ordered by size. Enrichment for selected gene sets and GO biological processes (top panel). The size and color of the circles indicate the significance level (-log_10_ *P-value*). Enrichments for PD heritability, using stratified LD score regression (bottom panel). The size and color of circles indicate the enrichment value (from LD score) and significance level (-log_10_ *P-value*) of enrichment, respectively. Only modules that were significant at a nominal *P-value <* 0.05 are shown here. **(B)** Eigengene analysis of all genes in the “mitochondrial” GO category (n = 1302) between PD and controls (Left panel; Wilcoxon rank-signed test, *P-value* = 0.0012). Example of a module (*green*) enriched for PD heritability, mitochondrial genes, and upregulated DEGs (Right panel). Edges represent co-expression connectivity. Nodes in orange are upregulated DEGs at FDR < 0.05; yellow triangles are genes in PD GWAS loci. **(C)** Eigengene analysis of all genes in the “lysosome” GO category (n = 526) between PD and controls (Wilcoxon rank-sum test, *P-value* = 0.0013) (Left panel). Example of a module (*salmon*) enriched for PD heritability, proteo-lysosomal genes, and downregulated DEGs. Nodes in orange are upregulated DEGs at FDR < 0.05; grey are selected proteo-lysosomal genes; and yellow triangles are genes in PD GWAS loci (Right panel).

### Common variants in PD susceptibility loci contribute to altered gene expression in monocytes

The majority of PD risk-associated variants are located in non-coding regions of the genome. It is reasonable to hypothesize that a subset of these may cause disease by altering gene regulatory mechanisms as either expression (eQTL) or splicing (sQTL) quantitative trait loci. Here, we performed *cis*-eQTL analysis using monocytes from 180 subjects of European ancestry (**Fig. S11**) to systematically interrogate PD risk loci from the most recent GWAS (*2*) to uncover putative PD-dysregulated genes based on gene expression and splicing regulation. We identified 4,030 and 1,786 genes with *cis*-eQTLs and sQTLs at FDR < 0.05, respectively (**Table S12, S13**). Using a mediated expression score regression (MESC) (*35*), we estimate 26% (S.E.10%) of PD disease heritability is mediated by the *cis*-genetic component of monocyte gene expression levels (**Fig. 4A**). This estimate in monocytes is similar to what we observed in primary microglia (23%; S.E. 16%) (*36*) but lower than in prefrontal cortex (40%; S.E. 11%) (*37*), suggesting that a substantial proportion of PD heritability can be attributed to other CNS cells. Nevertheless, given that a large proportion of PD disease heritability is mediated by eQTLs in myeloid cells (*3*), we performed colocalization analysis (*38*) to determine whether a shared variant is responsible for both GWAS and QTL signal in a locus. We found that GWAS and eQTL signals colocalized in 15 out of 78 PD loci, suggesting that the disease-associated SNP (or one in very high LD to it) drove variation in expression in monocytes (**Fig. 4B, Table S14**). We observed suggestive levels of colocalization (PPH4 > 0.5) of GWAS and eQTL at three additional loci, including at the *NOD2* locus (**Fig. 4C**), where the PD risk allele rs34559912-A decreases expression of *NOD2*. At the *LRRK2* locus, we observe that the PD risk allele rs76904798-T increases *LRRK2* expression in monocytes, consistent with what has previously been reported (*3*). We validate a previously identified eQTL at the cathepsin B (*CTSB*) locus, where the PD risk allele rs2740595-C decreases expression of *CTSB* (**Fig. 4C**). We found that the PD risk rs34559912-T allele, located within an intron of *BST1*, was strongly associated with lower expression of *BST1* (**Fig. 4C**). The genetic analysis suggests that decreased expression of *BST1* in monocytes is associated with increased risk for PD. Using a functional fine-mapping approach (see Methods), we found that the lead eQTL SNP (rs34559912) is also the top fine-mapped SNP (the SNP with the highest posterior probability of being causal within the 95% credible set) at the *BST1* locus (**Fig. 4D**) and is within a monocyte-specific enhancer **(Table S14)**. Notably, 60% (9 of 15 colocalized loci) of the lead eQTL SNP (or the lead GWAS SNP) are within CD14^+^ monocytes histone acetylation marks (H3K27ac) associated with enhancer activity, with one (rs34559912-*BST1*) specific to monocytes. Additionally, 27% (4 of 15 colocalized loci) of these lead eQTL SNPs (or the lead GWAS SNPs) are within microglia histone marks (H3K27ac), with two (rs6658353-*B4GALT3* and rs1293298-*CTSB*) specific to microglia (*39*). Of these four, all are within chromatin accessibility (ATAC-seq) peaks in microglia (*39*), and of which three and one are within a PU.1 enhancer and promoter, respectively **(Table S14)**. In a companion study, we have additionally fine-mapped all the PD GWAS loci and show that variants within the 95% credible sets for *CTSB, LRRK2, RAB29*, and *GPNMB* loci are located within microglia-specific enhancers (Schilder et al. *in prep*). In addition to expression, we performed a sQTL analysis to identify local genetic effects that drive variation in RNA splicing in monocytes. We observed six PD risk alleles affecting the splicing of nearby genes (**Fig. 4B**). An example is PD risk allele rs2306528-T associated with an exon skipping event in *FAM49B* (**Fig. 4E**), a novel regulator of mitochondrial function (*40*). These results suggest that PD risk alleles modulate disease susceptibility by regulating the expression or splicing of genes in peripheral monocytes.

**Figure 4.**
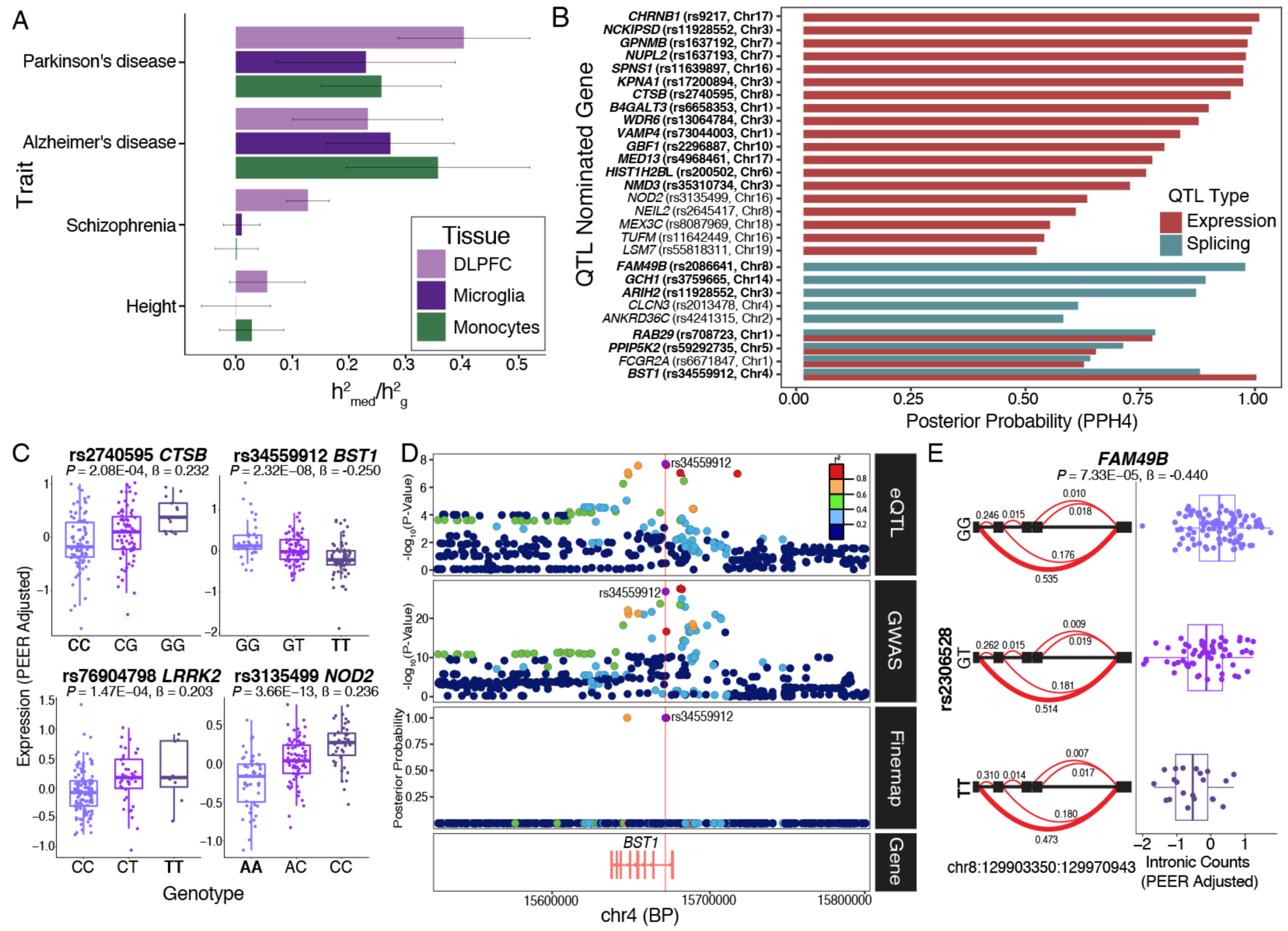
Parkinson’s disease susceptibility alleles alter gene expression in monocytes. **(A)** Estimated proportion of heritability mediated by *cis*-genetic component of expression (**□**^2^_med_/**□**^2^_g_) in monocytes, DLPFC (*37*), and microglia (*36*) for AD (*87*), PD (*2*), Schizophrenia (*88*) and Height (*89*) GWAS. **(B)** Colocalization of PD GWAS loci and monocyte *cis* expression or splicing QTLs. Shown in the bar plots are Posterior Probability (PPH4) from *coloc* (*38*) that supports the hypothesis (PPH4) that both eQTL (or sQTL) and PD GWAS share the same single variant. PD loci with suggestive colocalization (PPH4 > 0.5) are shown along with the eGene and the lead eQTL SNP (in LD with the lead GWAS SNP; r^2^ > 0.8). Genes in bold indicate reliable evidence in favor of a colocalized signal (defined as PPH3 + PPH4 > 0.8, PPH4/PPH3 > 2). **(C)** Boxplot of selected eQTLs with gene expression (PEER adjusted) per individual stratified by genotype. The eQTL *P-value* and effect size are listed on top. The PD GWAS effect allele is in bold. **(D)** Fine-mapping of the *BST1* locus. Colocalization of monocyte eQTL (top panel) and PD GWAS association (middle panel). Fine-mapping of *BST1* using PolyFun (*110*) prioritizes two variants within the 95% credible set (bottom panel), one of which is a lead eQTL SNP (rs34559912). **(E)** Example of an sQTL within *FAM49B* showing intronic ratios stratified by genotypes (left panel). The PD effect allele and most significant intronic excision (chr8:129903350:129970943) within *FAM49B* are in bold. The red (bold) line represents the most significant junction. sQTL boxplot of chr8:129903350:129970943 intronic excision ratio (PEER adjusted) per individual stratified by genotype (right panel).

### scRNA-seq profiling of PD CD14^+^ monocytes

Human monocytes are subdivided into at least three different subpopulations (classical, intermediate, and non-classical) according to their surface expression of the receptor CD14 and the Fc receptor CD16 (*41*). The three monocyte subsets are phenotypically and functionally different (*42, 43*). To investigate whether the composition and gene expression profiles of monocyte subpopulations are altered in PD, we used flow cytometry (FACS) and scRNA-seq analysis to characterize different monocyte subpopulations from PD patients and age-matched healthy controls. First, we performed FACS analysis to assess differences in proportion of monocyte subsets between PD and controls. Using FACS, we did not observe any differences in the proportions of monocyte subpopulations in a subset of PD (n = 11) and control (n = 11) samples (*P-value* > 0.05, unpaired t-test) (**Fig. S13A**), contrary to previous reports (*44, 45*). Secondly, we performed scRNA-seq of CD14^+^ monocytes by multiplexing 10 individuals (seven PD, three controls, **Table S15**) on the 10x Chromium system with an expected yield of 20,000 single-cells. We identified six clusters including two main subpopulations that were detected corresponding to classical (CD14^++^/CD16^-^) and a CD16^+^ population that corresponds to intermediate (CD14^++^/CD16^+^) monocytes (**Fig. 5A**). The non-classical (CD14^-^/CD16^++^) monocyte subpopulation was not captured in the scRNA-seq due to the use of CD14^+^ selection method. Similar to our findings with FACS, we found no differences in proportions of monocyte subpopulations in PD vs controls (*P-value* > 0.05; Wilcoxon signed-rank test) (**Fig. S13B**). After QC (**Fig. S13C-F**), we performed differential gene expression between the subpopulations (classical and intermediate) and observed that 927 total genes were differentially expressed at FDR < 0.05 (**Table S16**). As expected many of the DEGs between clusters were marker genes for classical monocytes (*CD14*) or for intermediate populations (*FCGR3A*). We found that genes implicated in mitochondrial and proteasomal function, pathways enriched for DEGs in bulk monocytes, were highly expressed in the intermediate population relative to the classical population, suggesting this subpopulation may be key to the disease. Specifically, genes that are members of the mitochondrial cytochrome c oxidase and NADH dehydrogenase families and proteasomal genes were highly expressed in the intermediate monocytes (**Fig. 5B**). We did observe some disease-related genes to be highly expressed in the classical subpopulation as well (e.g., *S100A8*). Finally, we performed differential expression analysis within each subpopulation and identified several DEGs that were only detected within the intermediate subpopulation but not in the bulk analysis. These included genes from several members of the complement component (*C1QA*), interferons (*IFITM2*), and chemokine (*CXCL16*) in the intermediate monocytes (**Fig. 5C, Table S17**). In summary, our scRNA-seq data enables the evaluation of molecular aspects of monocyte heterogeneity. Overall, these results suggest that intermediate monocytes, which comprise about ∼8% of circulating monocytes and are involved in the production of reactive oxygen species (ROS) and inflammatory responses, are affected at the transcriptional level in PD.

**Figure 5.**
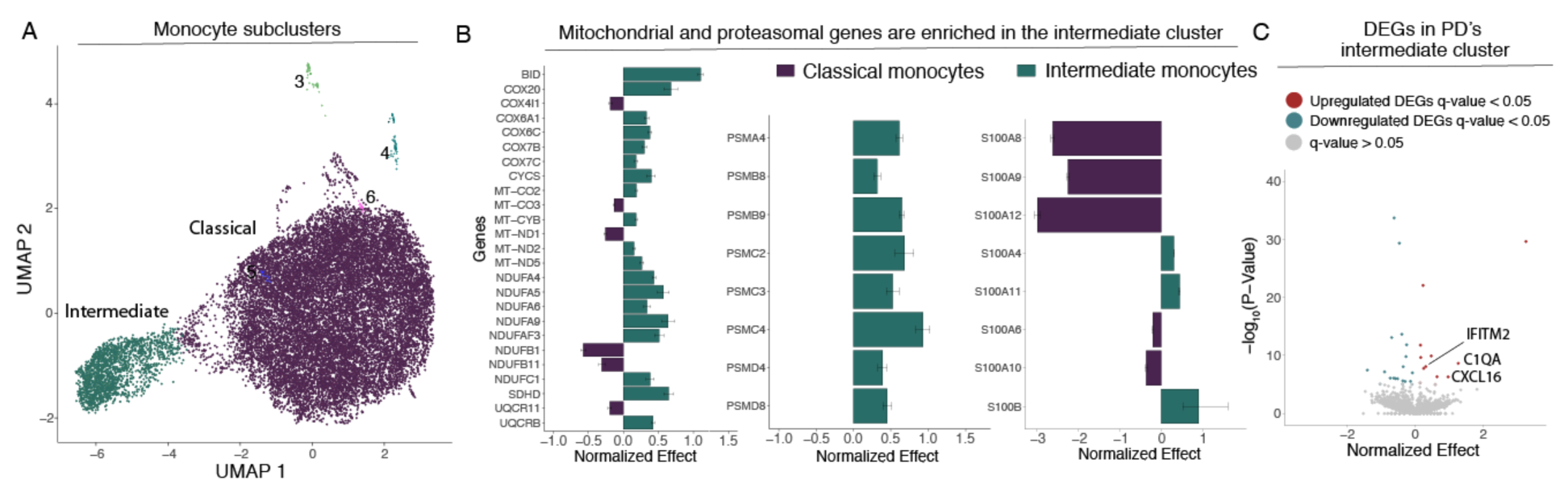
Single-cell profiling of CD14^+^ monocytes from PD and control subjects. **(A)** Generation of scRNA-seq from seven PD and three controls yielded 19,144 cells. Uniform Manifold Approximation and Projection (UMAP) visualization representing the six clusters including CD14^++^/CD16^-^ classical monocytes (purple) CD14^++^/CD16^+^ intermediate cluster (green). **(B)** Comparison of the relative levels of expression of mitochondrial and proteasomal genes in the classical vs. intermediate monocytes using normalized effect (*33, 111*). **(C)** Volcano plot showing the normalized effect within CD14^++^/CD16^+^ intermediate cluster of PD-monocytes and controls (x-axis) and their significance (y-axis, -log_10_ *P-value*). DEGs at q-value < 0.05 are highlighted in red (upregulated genes) and in blue (downregulated genes).

### Monocyte transcriptomic signature in PD is distinct from microglia

We next sought to determine whether PD monocyte signatures are recapitulated in primary microglia and postmortem brain tissues of autopsied PD subjects. To address this question, we isolated CD11b^+^ primary microglia from fresh post mortem autopsied brains of 13 donors with PD and 42 controls (with non-neurological diagnoses and including various causes of death such as euthanasia, cardio-respiratory disease or cancer, **Fig. S14A, B, Table S2**). For comparison, we also isolated microglia from patients with other brain disorders (including depression, Schizophrenia [SCZ], and Bipolar disorder). The microglia samples were isolated from multiple brain regions based on the availability of high-quality tissues from the brain banks (**Fig. S14A, B**). The resulting microglia samples were then subjected to RNA-seq using a low input library preparation. After rigorous QC and controlling for biological and technical covariates (**Fig. S14**), we performed differential gene expression between 22 PD and 106 control samples using a statistical method that accounts for repeated measures (in this case multiple brain regions from the same donor) while properly controlling the false discovery rate (*46*). Given the small sample size, we did not find any DEGs at FDR 0.05 in microglia but identified 222 DEGs at a suggestive threshold (FDR < 0.10) (**Table S18**). Interestingly, we found genes involved in mitochondrial function (*NDUFB5, NDUFA11* and *NDUFAB1*) (**Fig. 6B**), proteasomal function (*PSMB5, PSMG2, PSMB2*), complement component (*C1QA, C1QB*, and *C1QC*), and S100 calcium-binding proteins (*S100A4* and *S100A6*) (nominal *P-value* < 0.05). While the mitochondrial, proteasomal, and many of the S100 family genes were also differentially expressed in monocytes, the complement component genes were differentially expressed only in intermediate monocytes and in microglia. The directionality of fold change of DEGs in monocytes compared to microglia was concordant for some genes (e.g., *S100A4*) but we also observed discordant effects in these two cell types. In particular, we found discordant signatures for genes involved in mitochondrial OXPHOS. Specifically, the cytochrome c oxidase and NADH dehydrogenase family of genes were significantly upregulated in PD monocytes but were downregulated in PD microglia compared to controls (**Fig. 6C**). The discordant gene expression signature between monocytes and microglia is consistent across most nuclear-encoded mitochondrial genes (**Fig. 6C**). Due to the absence of an independent and sufficiently large microglia dataset from PD subjects, we were unable to directly replicate our findings. Nevertheless, we accessed an independent dataset obtained from meta-analysis of 8 studies with gene expression profiles from bulk brain substantia nigra (SN) of 83 PD cases and 70 controls (*47*). 20 out of 151 OXPHOS genes were significant at FDR < 0.05 in the meta-analysis DEG of SN and all 20 genes were downregulated in PD. Although this is not a direct replication as we only had access to independent tissue-level data (from SN) and not primary microglia, it highlights the downregulation of OXPHOS genes in post-mortem PD brains (**Fig. 6C**). We further found that the downregulation of mitochondrial gene signatures in microglia is specific to PD as we did not observe similar patterns in microglia DEGs of subjects with depression (n = 74), or a combination of SCZ and Bipolar disorder (n = 37) (**Fig. 6D**), although it remains to be seen if this signature is present in other neurodegenerative diseases. Taken together, our results show a reproducible discordant pattern of gene expression for mitochondrial OXPHOS genes in the periphery and the CNS of PD subjects.

**Figure 6.**
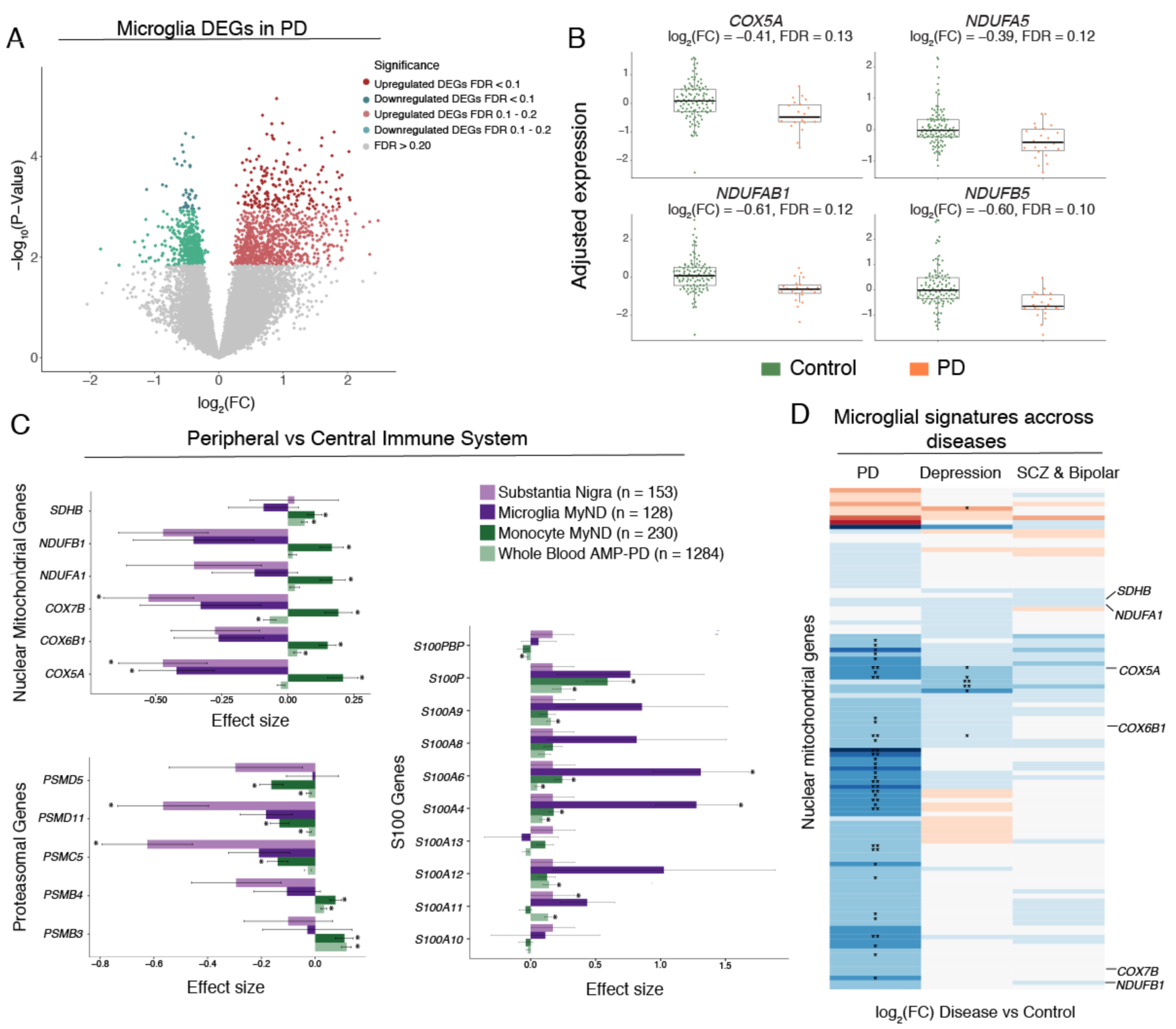
Comparing the transcriptome profiles of PD monocytes and primary microglia. **(A)** Volcano plot showing the fold-change of genes (log_2_ scale) between PD-microglia (22 samples from 13 donors) and controls (106 samples from 42 donors) (x-axis) and their significance (y-axis, -log_10_ scale). **(B)** Expression of selected mitochondria-specific genes in microglia. Adjusted gene expression levels after normalization are shown. **(C)** Effect size (log_2_[FC]) barplots of PD vs control differential expression in different datasets: substantia nigra (SN; light purple) (*47*), human microglia from MyND (dark purple), monocytes from MyND (dark green) and whole blood from AMP-PD (light green). Left panel: nuclear mitochondrial genes and proteasomal genes which are DEGs at FDR < 0.05 in monocytes from MyND. Right panel: All S100 genes tested across datasets. Corrected *P-value*: *FDR < 0.05 in all datasets; *FDR < 0.15 for microglia MyND. **(D)** Heatmap showing the fold-change (log_2_ scale) of disease vs controls of OXPHOS genes (y-axis) across different diseases (PD, Depression or Psychiatric disorders [Bipolar and Schizophrenia]). Blue represents log_2_(FC) < 0 (downregulated genes) and red represents log_2_(FC) > 0 (upregulated genes) when comparing disease vs. controls. Selected mitochondrial genes are shown. Nominal *P-value*: * *P-value* < 0.05; ** *P-value* < 0.01 for disease vs control differential expression.

## Discussion

Multiple lines of evidence implicate alterations in the immune system in PD (*25*), but the contribution of specific immune cells and their mechanisms in PD remains unclear. Here, we present a population-scale transcriptomic study of peripheral monocytes and primary microglia from subjects with PD. Our findings suggest widespread gene expression alterations in the PD myeloid cells, some of which are shared between the periphery and the CNS, while others have discordant effects. A key finding of our work is that genes in the mitochondrial respiratory chain are upregulated in peripheral monocytes but are downregulated in CNS microglia. Our single-cell-resolution analysis further suggests that these transcriptional alterations are specific to the intermediate monocyte subpopulation. By intersecting transcriptomics and genetics, we also demonstrate a large proportion (∼22%) of PD risk alleles alter the expression or splicing of genes in monocytes.

Although dysregulation of mitochondrial homeostasis in PD has been previously reported, these studies were mostly restricted to studying dopaminergic neurons, fibroblasts or blood from individuals with PD (*45, 48–54*). Some functional work of PD monocytes with limited sample size has been published before, reporting altered phagocytosis, metabolic alterations, and increased activation status upon different stimuli, (*13, 23, 44, 45, 55, 56*) but none of these studies performed unbiased transcriptome-wide gene expression analysis in a large PD cohort. Our work provides a unique view of monocyte transcriptome alterations associated with PD pathophysiology. We find robust evidence that OXPHOS genes are upregulated in PD peripheral monocytes, and more specifically, we show that OXPHOS genes are highly expressed in the pro-inflammatory intermediate CD14^++^/CD16^+^ subpopulation. We also generate a first (to our knowledge) unbiased transcriptomic dataset of freshly isolated microglia from PD and controls, where we observed the opposite effect, as the OXPHOS genes are downregulated in microglia. The data from CNS resident microglia and SN data presented in this study are consistent and confirm previous observations showing reduction of OXPHOS gene expression in post-mortem brains of individuals with PD (*48, 49*). However, contrary to most published results, we report an unexpected finding that OXPHOS genes are upregulated in peripheral monocytes from individuals with sporadic PD, a finding that has only been shown for two genes (*SDHB* and *ATP5A*) in PD lymphoblasts (*57*). A plausible explanation could be that PD monocytes reflect a hyperactive state with increased OXPHOS activity (*57*), and maybe responsible for the elevated oxidative stress in PD. Another possibility is that the increased OXPHOS activity in PD monocytes is a compensatory effect of dysfunctional mitochondria due to the rapid turnover of monocytes. In terms of the discordant expression of OXPHOS genes between periphery and CNS, unlike peripheral monocytes, which are short-lived and have a rapid turnover, CNS resident microglia have a longer lifespan (*58*) and may accumulate ROS-mediated mitochondrial damage over time (*59*). This increase in the production of ROS can cause a gradual accumulation of damage to mitochondrial activity in PD CNS cells but the rapid turnover of peripheral cells allows them to avoid the long-term adverse consequences of mitochondrial damage.

This work also improves our understanding of the PD-associated genetic risk factors influencing innate-immune mechanisms. Although a large proportion of PD heritability is mediated by the *cis*-genetic component of gene expression in neuronal tissues, our findings provide evidence that about ∼25% of PD heritability is estimated to be mediated by myeloid cell-specific *cis*-eQTLs. This estimate is consistent with our observation that in at least 17 loci, the PD risk variants are likely to modify disease susceptibility, at least in part, by modulating gene expression or splicing in peripheral monocytes. However, given that many of the monocytes lead eQTL SNPs (or fine-mapped credible set of SNPs) are also within microglia enhancers, it is plausible that the observed genetic effect on monocytes gene expression may be a proxy for infiltrating macrophages and/or resident microglia found at the sites of neuropathology. Given the current data, it is difficult to discern the exact cellular context in which these variants may act. It is also plausible that many of these eQTL-PD GWAS colocalizations may be identified in other CNS cell types (e.g., neurons, astrocytes or oligodendrocytes), where some of these genes are also expressed. Future studies incorporating eQTL datasets from primary human microglia, or from scRNA-seq will be an important resource in pinpointing the cellular contexts in which PD-causing genetic variants affect gene expression.

Our work has major implications for discovering novel blood-based biomarkers for differentiating individuals with PD from control individuals, as well as predicting the rate of PD progression. To date, biomarker studies in PD have largely focused on candidate approaches, with an emphasis on protein measures obtained in the CSF or brain imaging (*1, 60, 61*) which is considerably more difficult to obtain than blood. The use of monocyte gene expression to discover novel biomarkers of the disease state and/or its progression has several advantages. Firstly, monocytes isolated from peripheral blood are highly accessible human tissue, unlike the brain. While peripheral blood may be more easily obtained than primary monocytes, our results suggest that the magnitude of effects for DEGs was two-fold higher in monocytes compared to whole blood despite the smaller sample size (n = 230 in monocytes compared to 1,284 for whole blood from AMP-PD cohort). These results emphasize the power of purified cell populations that are not mixtures of cell types such as whole blood, which may result in the failure to properly capture the activity of cell-type-specific effects. Secondly, our study clearly shows that peripheral monocytes from PD cases differ from those of control subjects. To this end, we identified several genes whose expression is altered not only in PD monocytes but also exhibit altered expression levels in microglia and SN of individuals with PD. For example, the S100 family of genes whose upregulation is reproducible in all four datasets that we have compared (monocytes, whole blood, microglia, and SN) and which have also been shown to be upregulated in CSF of patients with PD (*30*) are excellent candidates for potential blood-based biomarkers. Further longitudinal studies are necessary to assess whether transcriptional changes in monocytes are predictive of disease progression.

In summary, by defining the transcriptional signatures of peripheral monocytes from sporadic PD patients, we have uncovered PD-associated alterations of mitochondrial and proteo-lysosomal genes in peripheral tissues. We demonstrate that although the same mitochondrial processes are altered in PD monocytes and microglia, the direction of effect of altered genes are distinct. Building on our data, future research should assess the functional bioenergetic properties of the CNS and peripheral tissues in sporadic PD to unravel potential mechanisms leading to the dysregulation described here. Overall, these results provide support for the utility of monocyte gene expression profiles as potent tools for understanding molecular mechanisms, for the identification of novel therapeutic targets, and for the development of blood-based biomarkers.

## Author contributions

T.R conceived the study. T.R, E.N, and E.U led the project, designed and performed the experiments and analysis, and wrote the manuscript. K.P.L. performed a co-expression network and microglial transcriptomic analysis. M.P. and E.N. optimized experimental approach. M.P., M.Z, and A.A. performed experimental isolations. B.M.S. and J.H. performed single-cell and splicing analyses, respectively. T.S., C.A., B.H., and S.S. contributed with clinical coordination. R.A.V. contributed to data analyses. G.R. and S.F. were responsible for the implementation and recruitment of movement disorder patients in NYUMD and BPMD. M.C. contributed to genetic and population characterization. D.R., S.E., R.A.O., V.S., M.S., S.B., and R.S-P. contributed with recruitment and clinical characterization of patients from MSBI; C.W.Z. and M.S. contributed to recruitment donors from ADRC; A.C.P. implemented recruitment of donors from CCH; R.R., R.H.W., and W.T. helped in recruitment patients from BPMD. T.A. and A.M.G. helped with intellectual discussion and interpretation of the results. K.F. R.H.W. and J.F.C. contributed to brain sample collection. G.J.L.S. and L.dW. contributed to microglial isolation and characterization. All authors read and approved the final manuscript.

## Material and methods

### Demographics and Clinical Overview of Study cohorts

This study is the first part of the MyND initiative. As part of this initiative, to date we have recruited 330 PD cases, 114 AD (Mild Cognitive Impairment [MCI] and AD; Clinical Dementia Rating [CDR] > 0.5), and 321 aged-match controls. In this study, 230 samples (including controls and PD patients) have been included. Samples were recruited from the following clinical cohorts: Movement Disorder Center at Mount Sinai Beth Israel (MSBI), Bendheim Parkinson and Movement Disorders Center at Mount Sinai (BPMD), Fresco Institute for Parkinson’s and Movement Disorders at New York University (NYUMD), the Alzheimer’s Research Center (ADRC) and Center for Cognitive Health (CCH) at Mount Sinai Hospital. This study was approved by the Institutional Review Board of each institution. Sample collection occurred during routine visits of the patients to the clinic, minimizing inconvenience to the patients and their families. All patients provided written informed consent for the collection of samples and subsequent analysis.

#### Mount Sinai Movement Disorder Centers (BPMD and MSBI)

PD subjects were recruited from two Mount Sinai Movement Disorder Centers: The Robert and John M. Bendheim Parkinson and Movement Disorders Center (BPMD) and the Mount Sinai Beth Israel (MSBI). Study participants from BPMD and MSBI included PD cases and family and friend controls who were part of two genetic studies of PD, one focused on gene identification in PD, and a second on ascertainment of biologic markers of glucocerebrosidase mutations. Affected individuals met the UK Parkinson’s disease Society Brain Bank (UKBB) criteria for probable PD (*62*), except that family history of PD was not an inclusion criteria. Controls did not have a history of a neurodegenerative disorder. All gave informed consent. Family history and pedigree were ascertained. A movement disorder trained neurologist assessed clinical features as well as performed the Unified Parkinson Disease Rating Scale (UPDRS)(*63*). Medical history and medications were captured.

#### New York Movement Disorder (NYUMD)

Patients affected with PD and controls were enrolled at the Marlene and Paolo Fresco Institute for Parkinson’s disease and Movement Disorders by Movement Disorder specialists between March 2018 and December 2019. Inclusion criteria were a diagnosis of PD according to the United Kingdom Parkinson’s Disease Society Brain Bank Clinical Diagnostic Criteria and age between 18 and 100 years (*64*). A population of aged and sex-matched non-affected subjects was enrolled among subjects who did not have a known diagnosis of PD at the time of evaluation and no history of other relevant neurological conditions. For each enrolled subject, both PD patients and controls, the following assessments were performed by qualified personnel and the following data were collected: demographic information (age, sex, self-reported ancestry), family history of PD, hand dominance, Montreal Cognitive Assessment (MoCA), UPDRS, H&Y rating scale, self-reported presence of the following motor and non-motor symptoms associated with PD (constipation, urinary symptoms, symptomatic orthostatic hypotension, subjective loss of sense of smell, REM sleep behavior disorders, hallucinations, anxiety, depression, motor fluctuations, dyskinesia, dopamine-related impulse control disorders), medication history, history of concomitant clinical conditions with particular attention of inflammatory diseases. For PD patients only the following information was also collected: age of onset of the disease (since onset of motor symptoms), symptoms at onset, and PD motor subtype (tremor dominant (TD) versus postural instability and gait difficulty (PIGD)). Data were collected in a password-protected database. Only subjects with the cognitive capacity to understand the study procedures, and the risks and benefits of the study, as assessed by licensed clinicians and established based on MoCA score greater or equal to 22, were enrolled.

#### Mount Sinai Alzheimer’s Disease Research Center

The Alzheimer’s Disease Research Center (ADRC) at the Icahn School of Medicine at Mount Sinai is a comprehensive research facility and clinical program dedicated to the study and treatment of normal aging and Alzheimer’s disease. With research into the causes of dementia, diagnostic services, and caregiver programs, the ADRC seeks to improve diagnosis, delay disease progression, and enhance the well-being of those affected by AD. The ADRC recruits participants who are cognitively normal, MCI, AD and other dementia into the National Alzheimer’s Coordinating Center Uniform Data Set (NACC UDS). Participants are followed annually and provide permission to contact them as additional studies become available including studies for the contribution of biosamples for new or ongoing projects. DNA is banked both locally and through the National Cell Repository for Alzheimer’s Disease (NCRAD). All subjects included have a Clinical Dementia Rating (CDR) of 0, thus only subjects cognitively normal were included in this study as controls.

#### Mount Sinai Center for Cognitive Health (CCH)

Through collaborations with Mount Sinai ADRC, the CCH evaluates individuals with concerns about cognition, such as memory, language and thinking difficulties. At the initial visit, a comprehensive neurological examination is performed which includes a thorough review of medical history, social history and detailed descriptions of cognitive complaints, and changes in behavior. The neurologists may do cognitive testing to assess memory, language, visual processing and other symptoms related to thinking. All subjects recruited from the CCH have a Mini-Mental State Exam (MMSE) test of cognitive function. Subjects that are cognitively normal based on cognitive testing were included in this study.

### PBMC and monocyte isolation

A maximum of 30 ml of blood was collected in Vacutainer blood collection tubes with acid citrate dextrose (ACD) (BD Biosciences). Fresh blood was shipped to the Raj laboratory and processed within 2-3 hours. First, blood was centrifuged at 1,500 g for 15 mins, and aliquots of whole blood and plasma were stored at -80 °C. Subsequently, blood was diluted in 2-fold PBS (Gibco) and PBMCs were isolated using SepMate tubes (StemCell Technologies) filled with 15 ml of Ficoll-Plaque PLUS (GE Healthcare) through a 15 mins centrifugation at 1,200 g. After washing with PBS, 5 million PBMCs were sorted to monocytes using CD14^+^ magnetic beads (Miltenyi) in the AutoMacs sorter and following manufacturer’s instructions. PBMCs and monocyte viability was assessed using Countess II Automated Cell counter (Thermo Fisher). After sorting, monocytes were stored in RLT buffer (Qiagen) + 1% 2-Mercaptoethanol (Sigma Aldrich) at -80 °C. Purity of the monocyte sorting was assessed via FACS and expression markers (RNA-seq). Remaining PBMCs were cryopreserved in 90% FBS (Germini) + 10% DMSO (Sigma Aldrich) at a concentration of 10 million cells/ml in Nalgene cryogenic vials (ThermoScientific). Vials were placed in NalGene CryoFreezing containers at -80 °C during 24-72 hours, and subsequently placed at liquid nitrogen for storage long-term.

### DNA isolation and genotyping

#### DNA isolation and genotyping

When isolating DNA from blood, an aliquot of 1 ml was used. We used the QiAamp DNA Blood Midi kit (Qiagen) and followed the manufacturer’s instructions. DNA quality and concentration was assessed using a Nanodrop. Samples were genotyped using the Illumina Infinium Global Screening Array (GSA), which contains a genome-wide backbone of 642,824 common variants plus custom disease SNP content (∼ 60,000 SNPs). Additionally, we performed targeted genotyping for specific regions associated with neurodegenerative diseases (LRRK2, GBA and APOE). LRRK2 and GBA genotyping was outsourced to the Dr. William Nichols’ laboratory at the Cincinnati Children’s Hospital. SNP genotyping was performed for the 2019S variant in *LRRK2* and the 11 most common variants in *GBA* (84GG, IVS2+1, E326K, T369M, N370S, V394L, D409G, L444P, A456P, RecNcil, R496H). The three major APOE isoforms (APOE 2, APOE 3, APOE 4) were assessed in the laboratory using Taqman assays for both rs429358 (C___3084793_20) and rs7412 (C___904973_10) from ThermoScientific following manufacturer’s instructions. 10 ng of DNA were added to the SNP reaction mix in a 96-well plate. Fluorescence reading of the Taqman assays was performed using QuantStudio 7 Flex (Applied Biosystems).

#### Genotype Quality Control and Imputation

We applied genotype quality control (QC) metrics such as SNP call rate > 95%, minor allele frequency (MAF) > 5%, Hardy-Weinberg equilibrium (HWE) *P-value* > 1 × 10^−6^, sample call rate > 95% to prepare high quality genotype data. Duplicated/related samples were determined based on pairwise IBD (identity-by-descent) estimation using PLINK (*65*). Duplicated samples with PLINK PI_HAT values between 0.99 to 1 were identified and files were converted to Variant Call Format (VCF) using VCFTools. Genetic ancestry of samples was confirmed by principal components analysis using PLINK; MDS (multidimensional scaling) values of study subjects were compared to those of 1000 Genome Project samples (Phase 3). An AJ only analysis was completed using a custom reference panel following that same protocol, and 82 samples (35.6%) were of AJ ancestry, 58 PD cases and 24 controls (**Fig. S1B, S11**).

Genotype Imputation was done using the Michigan Imputation Server v1.0.4 (Minimac 3) (*66*) using the 1000 genomes phase 3 v5 mixed panel and eagle v2.3 phasing in quality control and imputation mode. The following filtering was applied post imputation: SNPs that had imputation R^2^ > 0.3, removing multi-allelic SNPs, filtering MAF > 5% and HWE *P-value* > 1 × 10^−6^, and removing indels. We performed liftover of the imputed VCF files to hg38 using hg19toHg38 liftover chain file from UCSC Genome browser and liftoverVCF from Picard to match imputed genotypes to the hg38 reference used for RNA-seq. After imputation and liftover, a total of 5,951,770 variants were included in downstream analyses.

### Transcriptomic analysis

#### RNA isolation, library preparation and sequencing

RNA was isolated from monocyte samples stored in RLT buffer. After thawing on ice, RNA was isolated using RNeasy Mini kit (Qiagen) following the manufacturer’s instructions including the DNase I optional step. Once RNA was isolated, samples were stored at -80 °C upon library preparation. Prior to library preparation, RNA concentration was assessed using Qubit and the RNA integrity number (RIN) by TapeStation using Agilent RNA ScreenTape System (Agilent Technologies). The median for the RIN values across the cohort is 9.7. Only 8 samples showed RIN < 5 and removing these samples did not alter results. Library preparations were done either *in-house* or at Genewiz Inc. using in both cases ribo-depletion strategy to remove rRNA. For in-house library preparation, we used the TruSeq Stranded Total RNA Sample Preparation kit (Illumina), with the Low Sample (LS) protocol and followed the manufacturer’s instructions. For samples prepared at Genewiz, we shipped the RNA and samples were processed using the Standard RNA-seq protocol. Samples were sequenced in 3 independent batches at Genewiz Inc. with a depth of 60 million 150-bp paired-end reads using Illumina HiSeq 4000 platform.

#### RNA-seq data processing, quality control, and normalization

To process FASTQ files, we utilized RAPiD-nf, an efficient RNA-seq processing pipeline implemented in the NextFlow framework (*67*). RAPiD allows us to automate quality control, alignment and quantification of each RNA-seq sample. Following adapter trimming with trimmomatic (v0.36) (*68*), all samples were aligned to the hg38 build (GRCh38.primary_assembly) of the human reference genome using STAR (2.7.2a) (*69*) with indexes created from GENCODE (v30) (*70*). Gene expression was quantified using RSEM (1.3.1) (*71*); splice junction reads were extracted and quantified using Regtools (0.5.1) (*72*). Sequencing quality and technical metrics were assessed both before alignment with FASTQC (0.11.8) (*73*) and after alignment with Picard (2.20) (*74*) and Samtools (v1.9) (*75*).

As part of the RAPiD 3.0 pipeline, FASTQC was run for all samples and MultiQC was used to visualize and interpret the results. No samples were removed based on FASTQC metrics. Post alignment quality control of RNA-seq data was performed using Picard. Initial inclusion criteria consisted of at least 20 million passed reads, at least 20% of reads mapping to coding regions, and ribosomal rate < 30%. Additional QC was completed analyzing estimated counts, Transcripts Per Million (TPM), Counts Per Million (CPM), and TMM-voom (trimmed means of M-values) normalizations. Samples were removed if they were determined to be sex mismatches based on the expression of genes *UTY* and *XIST* compared to reported sex (**Fig. S3D**). Four samples were removed based on sex mismatches. Based on immune cell marker gene expression no samples were removed for having cell type contamination. Outliers were also removed after adjusting for covariates using dimensionality reduction through principal component analysis (PCA) and MDS that were selected to be used in differential analyses. Seven samples were removed after PCA and MDS analysis (**Fig. S3E)**.

Individual gene and transcript level counts and TPM used for downstream analyses were generated using RSEM and assembled to a matrix via the tximport R package. Then, CPM were calculated using cpm() function from the edgeR packing in R. Lowly expressed genes were filtered out, which were defined as having less than one count per million in at least 30% of the samples leading to a total of 13,667 genes.

#### Understanding sources of expression variation and covariate selection

To understand major sources of variation in the gene expression data, we used the R package variancePartition (*76*), which uses a linear mixed model to attribute a percentage of variation in expression based on selected covariates on a per gene basis. As highly correlated covariates cannot be included in the model, we selected covariates that were not very strongly correlated to run the variancePartition analysis (**Fig. S4A, B**). Gene counts were normalized using TMM values calculated from edgeR and *voom* transformed, which is a method that estimates the mean-variance relationship of the log-counts (*77*) as input to variancePartition. We ran several differential expression (DE) models and tested the suitability of each of them. In all the models we included sex, age, RIN value and the four MDS from the genotypes (which are enough to separate different populations, see *Genotyping and QC* section) as covariates. For each model, we added different covariates ranging from the simplest (no additional covariates) to the most complex (all covariates from variancePartition), with all the intermediate designs. None of the covariates were correlated with diagnosis except an expected correlation with sex. In all cases, there was a high fold change correlation between differentially expressed genes (DEGs) generated from the different designs (>0.90) and high percentage of shared DEGs and similar pathways. We decided to use a design which includes those covariates that explained the most variance in gene expression (on average across genes) according to variancePartition results, which is as follows: *expression* ∼ *rna_batch* + *age* + *sex* + *RIN* + *PCT_USABLE_BASES* + *PCT_RIBOSOMAL_RNA* + *MDS1* + *MDS2* + *MDS3* + *MDS4*.

#### Differential Expression Analysis

Differential expression analysis was performed between PD cases and controls through a linear model using the R package limma version 3.38.3 (*78*). For this analysis, inputs included the count matrix and the covariate file. These data were normalized using TMM values calculated from edgeR and *voom* transformed (**Fig. S5A**). Limma fits a linear model, and then runs a Bayesian moderated t-test which provides a *P-value. P-values* were then adjusted for multiple testing correction using the Benjamini-Hochberg FDR correction, which is implemented in the limma package. Description and selection of the covariates used can be found in the *understanding sources of variation section*. Differential isoform expression was performed following the same protocol.

#### Pathway and Gene Set Enrichment analysis

(i) *Pathway analysis:* we performed pathway analysis independently using the following input gene sets: upregulated DEGs (162), downregulated DEGs (138), differential splicing events (161) and DE transcripts (939) at FDR < 0.05. We used GSEA (*79*), focusing on Biological processes from Gene Ontology and limiting to gene sets between 10-500 genes. We show the 20 more significant enriched pathways with at least five genes that overlap. (ii) *Gene set enrichment analysis*: to test specific pathways we used curated gene sets and tested statistical enrichment using Fisher exact test. The pathway lists were arranged as follows: (1) All gene ontology gene sets were downloaded from the amigo.geneontology.org resource searching for the specific pathways: mitochondria (315 genes), proteasome (450 genes), lysosome (682), inflammatory response (694). (2) Mitochondrial curated list: 315 genes (*27*). From this gene list, we separated the different specific mitochondrial pathways (OXPHOS, Mitonuclear cross-talk and mitochondrial dynamics) following the paper specifications. (3) Proteasomal curated list: 39 genes (*80*). (4) Ubiquitin-related curated list: 428 genes combined from ubiquitin-like modifier activating enzymes (HGNC dataset), ubiquitin conjugating enzymes E2 (HGNC dataset) and ubiquitin ligase E3. (5) Lysosomal curated list: 435 genes from The Human Lysosome Gene Dataset.

### Splicing analysis

Differential splicing (DS) was assessed using Leafcutter (*81*). Leafcutter pools splice junction-spanning reads from each sample together and clusters junctions that overlap at either end. Differential splicing is then defined in a shift of junction usage within a cluster between two groups. Firstly, splice junction reads were extracted from each BAM file using regtools (*72*) and any junction reads aligned to scaffold chromosomes were removed. All junction files were clustered using leafcutter_cluster_regtools.py, specifying for each junction in a cluster a maximum length of 100kb. This led to 194,127 junctions within 45,631 clusters. We used a custom script to restrict our analysis set to junctions present in at least 25% of samples contributing at least 5% of the total reads to their cluster (**Fig. S7A**). Any cluster with only a single junction remaining after filtering or more than 10 junctions were removed. This led to a final set of 22,888 junctions within 8,882 clusters. Differential splicing between PD cases and controls was performed, testing each cluster if it was present in at least 50 samples per group with a minimum coverage of 20 reads in total. The same covariates were used in model fitting as for the differential expression analysis. Results were visualized using the LeafViz browser. Junction ratios were corrected for covariates using the “quantify_PSI.R” script found within the “psi_2019” branch of Leafcutter.

### Parkinson’s Disease Progressive Marker Initiative (PPMI) RNA-seq data analysis

RNA-seq counts and TPMs generated from whole blood from the PPMI (part of the AMP-PD cohorts) were downloaded from AMP-PD Knowledge Platform (*82*). AMP-PD has RNA-seq FASTQ files and workflow products from Salmon, STAR, and featureCounts for the PPMI cohort. All RNA sequencing was performed by Hudson Alpha and is supplied along with corresponding clinical data. RNA-seq samples from the baseline visit were extracted for subjects with idiopathic PD and controls. We followed a similar QC pipeline to our monocyte transcriptome analysis (**Fig. S8**). Differential expression analysis was performed with the read counts generated from rsubread featureCounts using the R package limma, adjusting for the following covariates: *% usable bases* + *% intergenic bases* + *sex* + *age* + *race* + *ethnicity* + *plate* (**Fig. S8**).

### Co-expression Network Analysis

Prior to the network analysis, expressed genes were filtered by protein-coding according to GENCODE annotation version 30 (n = 11,475 protein-coding genes), and expression data was transformed using *voom*. To minimize the effect of confounders in our dataset we used the “num.sv” function in the Bioconductor package *sva* embedded with the permutation-based approach algorithm “be” (*83, 84*) to get the number of surrogate variables (SVs) and correct the expression data. This approach estimated 12 SVs to be regressed from the whole matrix, including PD cases and controls (n = 230 samples). Then, using the *sva_network* package, we computed the SV loadings of the standardized expression matrix with singular value decomposition (SVD), and computed the residuals after regressing the top 12 SVs. Linear regression between the SVs and the covariates showed correlation mostly with technical covariates, including lane, batch, percentage of ribosomal bases and other sequencing metrics such as % of mRNA and intergenic bases (**Fig. S9A**).

The co-expression network analysis was performed using the R package of Weighted Gene Correlation Network Analysis (WGCNA) (*85*) following the standard pipeline to fit a scale-free topology (R^2^ > 0.8) and applying a Soft Threshold power of 5 into a signed network model (**Fig. S9B**). The adjacency matrices were constructed using the average linkage hierarchical clustering of the topological overlap dissimilarity matrix (1-TOM). Coexpression modules were defined using a dynamic tree cut method with minimum module size of 20 genes and deep split parameter of 4. Modules highly correlated with each other, corresponding to a module eigengene (ME) correlation of 0.75 were merged, resulting in a total of 65 modules (**Fig S9D**). The genes were prioritized based on their module membership value, also known as eigengene-based connectivity (kME). Highly connected intramodular hub genes tend to have high module membership in their respective module. The top hub genes for each module are shown in **Table S11**. Since we were interested in modules associated with PD, we calculated the Pearson correlation between the MEs and disease diagnosis, and prioritized those modules with FDR adjusted *P-value* from a Wilcoxon rank-signed test. Network visualization was done using the “exportNetworkToCytoscape” function from *WGCNA* R package to export the lists of nodes and edges, and the *ggraph* R package (*86*) to create the figures.

### Heritability analysis

Stratified LD score regression (S-LDSC) (*34*) was used to partition SNP-based disease heritability within each co-expression module. Using GWAS summary statistics from PD and LD modeled from 1000 genomes reference panel of European ancestry, we calculated the proportion of genome-wide SNP-based heritability that be attributed to SNPs (by mapping SNPs within each gene plus 10 kb +/- from transcript start and stop sites) within each module. To improve model accuracy, the LD-scores from each co-expression module were added to the ‘full baseline model’ which included 53 functional categories capturing a broad set of functional and regulatory elements. Enrichment is defined as the proportion of SNP-heritability accounted for by each module divided by the proportion of total SNPs within the module. Modules with FDR-corrected enrichment *P-values* of less than 0.05 were considered significant heritability contributors. Mediated expression score regression (MESC) method (*36*) was used to estimate disease heritability mediated by the *cis* genetic component of gene expression levels. The expression scores were estimated using eQTL summary statistics from monocytes (this study), microglia and DLPFC. The MESC python script also estimates the expression *cis*-heritability of each gene (using LDSC for eQTL summary statistics). The expression scores and GWAS summary statistics (PD, AD, Schizophrenia, and Height) (*2, 87–89*) were then used to estimate expression-mediated heritability (h2med).

### Quantitative Trait Loci Analysis

To perform eQTL mapping, following the GTEx pipeline (*90*) we completed a separate normalization and filtering method to previous analyses. Gene expression matrices were converted to BED format, TMM normalized, filtered for lowly expressed genes, removing any gene with less than one TPM in 20% of samples and at least 6 counts in 20% of samples, and each gene was inverse normal transformed across samples. After filtering, we tested a total of 18,430 genes. Then, PEER (*91*) factors were calculated to estimate hidden confounders within our expression data. We created a combined covariate matrix that included the PEER factors and the first four genotyping ancestry MDS values as input to the analysis. We used 15 PEER factors as covariates in our QTL model (**Fig. S12A**). To confirm that our DNA and RNA samples were from the same donor, we used mbv from QTLtools (*92*). Based on this, we removed 7 samples from QTL analysis. To test for cis-eQTLs, linear regression was performed using the QTLtools nominal pass for each SNP-gene pair using a one megabase window within the transcription start site (TSS) of a gene. To test for association between gene expression and the top variant in *cis* we used the QTLtools permutation pass which performs gene-based permutation with 1000 permutations. To identify eGenes, we performed FDR correction (using a threshold of ≤ 0.05) on the *P-value* of association adjusted for the number of variants tested in *cis* given by the fitted beta distribution. We estimated replication of MyND monocyte cis-eQTLs (discovery) using CD14^+^ eQTL data set from Fairfax et al. (replication) (*93*) using the q-value R package to estimate π1 (**Fig. S12C**).

To perform splicing quantitative trait loci analysis (sQTLs), we used junction counts generated from regtools. All junction files were clustered using the Leafcutter script, specifying for each junction in a cluster a maximum length of 100kb. Following the GTEx pipeline, introns without read counts in at least 50% of samples or with fewer than 10 read counts in at least 10% of samples were removed. Introns with insufficient variability across samples based on the thresholds again provided by GTEx consortium (*94*) leaving us with a final set of 107,838 junctions within 35,056 clusters. Filtered counts were then normalized using prepare_phenotype_table.py from leafcutter, merged, and converted to BED format, with the start/end positions from the gene to which an intron was mapped. We created a combined covariate matrix that includes the PEER factors and the first 4 genotyping ancestry MDS values as input to the analysis. We used 15 PEER factors as covariates in our QTL model (**Fig. S12B**). QTL mapping was performed as before with QTLtools, testing all variants within 1 megabase of the transcription start site, with 1000 permutations, grouping SNPs by gene. Genes with splicing QTLs were identified by FDR correction (<0.05) of the permutation *P-values*.

### Colocalization analysis

#### GWAS Data

we used the latest PD GWAS full summary statistics (*2*). Liftover of the full summary statistics from hg19 to hg38 was performed using GWAS harmonization (*95*). Because of differences in LD between populations, we completed a European only QTL analysis and used these results as input for colocalization and fine-mapping analyses. To perform the colocalization analysis between GWAS and eQTL data, we used the coloc.abf function from the coloc package (*38*) with default parameters. Our criteria for considering a signal to be colocalized was PPH3 + PPH4 > 0.8, PPH4/PPH3 > 2. All SNPs tested within 1 Mb either side of each GWAS locus were considered for colocalization analysis. To annotate SNPs, we incorporated CD14^+^ monocyte H3K27ac marks from HaploReg v4.1 (*96*) and microglia H3K27ac marks, ATAC-seq peaks, and PU.1 annotations from Nott et al. (*39*) data on the UCSC genome browser.

#### Fine-mapping

To functionally fine-map PD GWAS loci, we used PolyFun+SuSiE (*97, 98*) which computes SNP-wise heritability-derived prior probabilities using a L2-regularized extension of stratified LD SCore (S-LDSC) regression (*34, 99, 100*). A UK Biobank baseline model composed of 187 binarized epigenomic and genomic annotations was used as the annotation input (*101*). We applied PolyFun+SuSiE to PD GWAS summary statistics and LD reference generated from 337K UK Biobank individuals of white British ancestry.

### Single-cell RNA-seq data generation and processing

Using cryopreserved PBMCs, monocytes were isolated for scRNA-seq following the same protocol as previously described. scRNA-seq with multiplexed cell hashing (*102*) was performed at the New York Genome Center on purified monocytes from 10 donors, including three controls and seven PD patients (two with GBA mutations and one with a LRRK2 mutation). We first used the R package *Seurat* (v3.1.0) (*103, 104*) to remove non-protein-coding genes identified through *biomaRt* (*105*), keeping 14,827 protein-coding genes out of a total 24,914 genes. We also filtered-out low-quality cells that expressed less than 200 genes or over 2,500 genes, and cells that expressed greater than 5% mitochondrial genes using the “FilterCells*”* function in *Seurat*, reducing the total number of cells from 22,113 to 19,144. Lastly, expression counts were normalized using the “preprocess_cds*”* function in *monocle3* with unique molecular identifier (UMI) count and % mitochondrial genes as covariates. Dimensionality reduction was then performed using Uniform Manifold Approximation and Projection (UMAP) (*106, 107*) via the “reduce_dimension*”* function in *monocle3* (*33*). To identify cell subpopulations, we applied Louvain clustering via the “cluster_cells*”* function in *monocle3* using only the top 2000 most variable genes (identified with the “FindVariableGenes*”* function in *Seurat*) (**Fig. S13C**). This yielded 6 discrete clusters, of which the largest two were identified as Classical (Cluster 1; CD14^++^/CD16^-^) and Intermediate (Cluster 2; CD14^++^/CD16^+^) monocytes based on the expression of cell-type markers. We performed differential expression between PD and controls without considering the independent subclusters (“across-clusters”) as replication of the bulk RNA-seq. We also identified differentially expressed genes between Classical and Intermediate monocyte clusters with the “fit_models*”* function in *monocle3*, which by default fits a generalized linear model for each gene with a quasi-Poisson expression response function, calculates coefficients under the Wald test, and corrects for multiple hypothesis testing using false discovery rate (*108*).

### Human microglia isolation and transcriptome data generation

#### Fresh isolation of human microglia

Post-mortem brain samples were obtained from the Netherlands Brain Bank (NBB) and the Neuropathology Brain Bank and Research CoRE at Mount Sinai Hospital. Informed consent for autopsy and necessary clinical data was previously obtained. We included clinical diagnosis without neuropathological confirmation. Brain tissue was stored in Hibernate media (Gibco) at 4 °C upon processing, which happened within 24 hours after autopsy (**Fig. S14**). Microglia were isolated from the following regions: corpus callosum (CC; 13 samples), medial frontal gyrus (MFG; 40 samples), superior temporal gyrus (STG; 30 samples), thalamus (THA; 23 samples), sub-ventricular zone (SVZ; 18 samples) and substantia nigra (SN; 1 sample). Microglia were isolated as previously described before (*109*) with minor modifications. In brief, tissue was first mechanically homogenized with the help of cell strainer and pipetting following enzymatic digestion with 0.33 mg/ml of DNase I (Sigma Aldrich) and 0.2% of Trypsin (Invitrogen) in a shaking incubator (140 rpm, 37 °C) for 30 minutes. After washing the tissue in GKN/BSA buffer (PBS + 2 g/L d**-**(1)**-**glucose + 0.3% bovine serum albumin (BSA), pH 7.4), cells were resuspended in 20 ml of GKN/BSA and 10 ml of Percoll (GE Healthcare) was added to the top drop-wise. The Percoll gradient was generated with 40 minutes of centrifugation at 4000 rpm 4 °C with no brake. The top myelin phase was discarded and the second layer, mainly containing astrocytes and microglia, was transferred to a new tube. Microglia were purified using human CD11b^+^ magnetic beads (Miltenyi) following the manufacturer’s instructions and the manual magnetic sorter. Microglia samples were stored in RLT buffer + 1% 2-Mercaptoethanol. RNA was isolated using RNeasy Mini kit adding the DNase I optional step. Library preparation was performed at Genewiz using the Ultra-low input system which uses Poly-A selection to remove the rRNA. Purity of microglia was confirmed by qPCR comparing the homogenate, positive and negative fraction. Briefly, RNA was reversed transcribed to cDNA and qPCR was performed using Taqman assays (ThermoScientific) for targeted genes. Fluorescence reading of the Taqman assays was performed using QuantStudio 7 Flex (Applied Biosystems). Results were analyzed using the comparative threshold cycle (Ct) and expressed as fold-change vs homogenate.

#### Transcriptomic analysis

RNA-seq data were processed using the RAPiD pipeline, with the same configuration as MyND analysis. RNA-seq QC was performed by applying three filters to remove samples (considering the whole cohort): (i) less than 10 million reads aligned to the reference genome (GRCh38) using the STAR aligner; (ii) samples with more than 20% of the reads aligned to ribosomal regions; (iii) samples with less than 10% of the reads mapping to coding bases. Gene counts were generated by RSEM and tximport. Genes with more than 1 cpm in 30% of the samples were kept for downstream analysis. Differential expression was performed using the DREAM method (*46*) from variancePartition R package (*76*) to account for repeated measures (**Fig. S14F**). Since each donor can contribute multiple samples from different brain regions (**Fig. S14B**), we modeled the donor as a random effect and added selected covariates to adjust for possible technical and biological confounders. In order to determine the covariates to add to the model we ran variancePartition (**Fig. S14E**). The final model used was *expression ∼ donor_id* + *tissue* + *sex* + *age* + *fastqc_percent_gc* + *featurecounts_assigned* + *picard_pct_mrna_bases* + *picard_pct_pf_reads_aligned* + *picard_pct_ribosomal_bases* + *lane.*

## Acknowledgements

We thank the study participants for providing blood samples and for their generous gifts of brain donation to the MyND study. We thank the Netherlands Brain Bank and the Neuropathology Brain Bank & Research Core at Mount Sinai for assistance in collecting human brain samples. We thank the Flow Cytometry Core and the Human Immune Monitoring Center at Icahn School of Medicine at Mount Sinai (ISMMS) for optimization of cell isolations; Genewiz Inc. for RNA-sequencing; Christos Proukakis and Robert O. Watson for feedback on the manuscript; Seunghee Kim-Schulze for help optimizing PBMC and monocyte isolation; Ying-Chih Wang for help with processing RNA-seq data; Javier Fernandez-Lopez for help with data analysis; members of the Ronald Loeb Center for Alzheimer’s disease for helpful discussion; research participants and employees of 23andMe who contributed to the PD GWAS. Data used in the preparation of this article were obtained from the AMP-PD Knowledge Platform. AMP-PD – a public-private partnership – is managed by the NIH and funded by Celgene, GSK, the Michael J. Fox Foundation for Parkinson’s Research, the National Institute of Neurological Disorders and Stroke, Pfizer, Sanofi, and Verily. PPMI – a public-private partnership – is funded by the Michael J. Fox Foundation for Parkinson’s Research and funding partners (names of all of the PPMI funding partners found at www.ppmi-info.org/fundingpartners). The PPMI Investigators have not participated in reviewing the data analysis or content of the manuscript. For up-to-date information on the study, visit www.ppmi-info.org.

## Data and materials availability

Raw RNA-seq and scRNA-seq data generated in this work are available through Sequence Research Archive (SRA) (in progress). Imputed genotype data are available from the Database of Genotypes and Phenotypes (dbGAP) (in progress). Processed read counts and full eQTL summary statistics are available from Zenodo online data sharing portal (in progress).

## References

1. W. Poewe, K. Seppi, C. M. Tanner, G. M. Halliday, P. Brundin, J. Volkmann, A.-E. Schrag, A. E. Lang, Parkinson disease. Nat Rev Dis Primers. 3, 17013 (2017).

2. M. A. Nalls, C. Blauwendraat, C. L. Vallerga, K. Heilbron, S. Bandres-Ciga, D. Chang, M. Tan, D. A. Kia, A. J. Noyce, A. Xue, J. Bras, E. Young, R. von Coelln, J. Simón-Sánchez, C. Schulte, M. Sharma, L. Krohn, L. Pihlstrøm, A. Siitonen, H. Iwaki, H. Leonard, F. Faghri, J. R. Gibbs, D. G. Hernandez, S. W. Scholz, J. A. Botia, M. Martinez, J.-C. Corvol, S. Lesage, J. Jankovic, L. M. Shulman, M. Sutherland, P. Tienari, K. Majamaa, M. Toft, O. A. Andreassen, T. Bangale, A. Brice, J. Yang, Z. Gan-Or, T. Gasser, P. Heutink, J. M. Shulman, N. W. Wood, D. A. Hinds, J. A. Hardy, H. R. Morris, J. Gratten, P. M. Visscher, R. R. Graham, A. B. Singleton, 23andMe Research Team, System Genomics of Parkinson’s Disease Consortium, International Parkinson’s Disease Genomics Consortium, Identification of novel risk loci, causal insights, and heritable risk for Parkinson’s disease: a meta-analysis of genome-wide association studies. Lancet Neurol. 18, 1091–1102 (2019).

3. Y. I. Li, G. Wong, J. Humphrey, T. Raj, Prioritizing Parkinson’s disease genes using population-scale transcriptomic data. Nat. Commun. 10, 994 (2019).

4. S. A. Gagliano, J. G. Pouget, J. Hardy, J. Knight, M. R. Barnes, M. Ryten, M. E. Weale, Genomics implicates adaptive and innate immunity in Alzheimer’s and Parkinson’s diseases. Annals of Clinical and Translational Neurology. 3 (2016), pp. 924–933.

5. T. Raj, K. Rothamel, S. Mostafavi, C. Ye, M. N. Lee, J. M. Replogle, T. Feng, M. Lee, N. Asinovski, I. Frohlich, S. Imboywa, A. Von Korff, Y. Okada, N. A. Patsopoulos, S. Davis, C. McCabe, H.-I. Paik, G. P. Srivastava, S. Raychaudhuri, D. A. Hafler, D. Koller, A. Regev, N. Hacohen, D. Mathis, C. Benoist, B. E. Stranger, P. L. De Jager, Polarization of the effects of autoimmune and neurodegenerative risk alleles in leukocytes. Science. 344, 519–523 (2014).

6. R. H. Reynolds, International Parkinson’s Disease Genomics Consortium (IPDGC), J. Botía, M. A. Nalls, J. Hardy,S. A. Gagliano Taliun, M. Ryten, System Genomics of Parkinson’s Disease (SGPD), Moving beyond neurons: the role of cell type-specific gene regulation in Parkinson’s disease heritability. npj Parkinson’s Disease. 5 (2019),, doi: 10.1038/s41531-019-0076-6.

7. E.-J. Bae, H.-J. Lee, E. Rockenstein, D.-H. Ho, E.-B. Park, N.-Y. Yang, P. Desplats, E. Masliah, S.-J. Lee, Antibody-Aided Clearance of Extracellular -Synuclein Prevents Cell-to-Cell Aggregate Transmission. Journal of Neuroscience. 32 (2012), pp. 13454–13469.

8. I. Choi, Y. Zhang, S. P. Seegobin, M. Pruvost, Q. Wang, K. Purtell, B. Zhang, Z. Yue, Microglia clear neuron-released α-synuclein via selective autophagy and prevent neurodegeneration. Nat. Commun. 11, 1386 (2020).

9. M. F. Duffy, T. J. Collier, J. R. Patterson, C. J. Kemp, K. C. Luk, M. G. Tansey, K. L. Paumier, N. M. Kanaan, D. Luke Fischer, N. K. Polinski, O. L. Barth, J. W. Howe, N. N. Vaikath, N. K. Majbour, O. M. A. El-Agnaf, C. E. Sortwell, Correction to: Lewy body-like alpha-synuclein inclusions trigger reactive microgliosis prior to nigral degeneration. Journal of Neuroinflammation. 15 (2018),, doi: 10.1186/s12974-018-1202-9.

10. L. Fellner, R. Irschick, K. Schanda, M. Reindl, L. Klimaschewski, W. Poewe, G. K. Wenning, N. Stefanova, Toll-like receptor 4 is required for α-synuclein dependent activation of microglia and astroglia. Glia. 61 (2013), pp. 349–360.

11. K. J. Doorn, T. Moors, B. Drukarch, W. van de Berg, P. J. Lucassen, A.-M. van Dam, Microglial phenotypes and toll-like receptor 2 in the substantia nigra and hippocampus of incidental Lewy body disease cases and Parkinson¿s disease patients. Acta Neuropathologica Communications. 2 (2014), p. 90.

12. C. W. Olanow, C. Warren Olanow, M. Savolainen, Y. Chu, G. M. Halliday, J. H. Kordower, Temporal evolution of microglia and α-synuclein accumulation following foetal grafting in Parkinson’s disease. Brain. 142 (2019), pp. 1690–1700.

13. V. Grozdanov, L. Bousset, M. Hoffmeister, C. Bliederhaeuser, C. Meier, K. Madiona, L. Pieri, M. Kiechle, P. J. McLean, J. Kassubek, C. Behrends, A. C. Ludolph, J. H. Weishaupt, R. Melki, K. M. Danzer, Increased Immune Activation by Pathologic α-Synuclein in Parkinson’s Disease. Ann. Neurol. 86, 593–606 (2019).

14. J. Herz, A. J. Filiano, A. Smith, N. Yogev, J. Kipnis, Myeloid Cells in the Central Nervous System. Immunity. 46, 943–956 (2017).

15. A. S. Harms, V. Delic, A. D. Thome, N. Bryant, Z. Liu, S. Chandra, A. Jurkuvenaite, A. B. West, α-Synuclein fibrils recruit peripheral immune cells in the rat brain prior to neurodegeneration. Acta Neuropathol Commun. 5, 85 (2017).

16. A. S. Harms, A. D. Thome, Z. Yan, A. M. Schonhoff, G. P. Williams, X. Li, Y. Liu, H. Qin, E. N. Benveniste, D. G. Standaert, Peripheral monocyte entry is required for alpha-Synuclein induced inflammation and Neurodegeneration in a model of Parkinson disease. Exp. Neurol. 300, 179–187 (2018).

17. C. Bliederhaeuser, V. Grozdanov, A. Speidel, L. Zondler, W. P. Ruf, H. Bayer, M. Kiechle, M. S. Feiler, A. Freischmidt, D. Brenner, A. Witting, B. Hengerer, M. Fändrich, A. C. Ludolph, J. H. Weishaupt, F. Gillardon, K. M. Danzer, Age-dependent defects of alpha-synuclein oligomer uptake in microglia and monocytes. Acta Neuropathol. 131, 379–391 (2016).

18. R. S. Wijeyekoon, D. Kronenberg-Versteeg, K. M. Scott, S. Hayat, J. L. Jones, M. R. Clatworthy, R. A. Floto, R. A. Barker, C. H. Williams-Gray, Monocyte Function in Parkinson’s Disease and the Impact of Autologous Serum on Phagocytosis. Front. Neurol. 9, 870 (2018).

19. H. Braak, K. Del Tredici, U. Rüb, R. A. I. de Vos, E. N. H. Jansen Steur, E. Braak, Staging of brain pathology related to sporadic Parkinson’s disease. Neurobiol. Aging. 24, 197–211 (2003).

20. F. Scheperjans, P. Derkinderen, P. Borghammer, The Gut and Parkinson’s Disease: Hype or Hope? Journal of Parkinson’s Disease. 8 (2018), pp. S31–S39.

21. G. Chapelet, L. Leclair-Visonneau, T. Clairembault, M. Neunlist, P. Derkinderen, Can the gut be the missing piece in uncovering PD pathogenesis? Parkinsonism Relat. Disord. 59, 26–31 (2019).

22. L. Klingelhoefer, H. Reichmann, Pathogenesis of Parkinson disease—the gut–brain axis and environmental factors. Nat. Rev. Neurol. 11, 625–636 (2015).

23. J. C. M. Schlachetzki, I. Prots, J. Tao, H. B. Chun, K. Saijo, D. Gosselin, B. Winner, C. K. Glass, J. Winkler, A monocyte gene expression signature in the early clinical course of Parkinson’s disease. Sci. Rep. 8, 10757 (2018).

24. V. Donega, S. M. Burm, M. E. van Strien, E. J. van Bodegraven, I. Paliukhovich, H. Geut, W. D. J. van de Berg, K. W. Li, A. B. Smit, O. Basak, E. M. Hol, Transcriptome and proteome profiling of neural stem cells from the human subventricular zone in Parkinson’s disease. Acta Neuropathol Commun. 7, 84 (2019).

25. E.-K. Tan, Y.-X. Chao, A. West, L.-L. Chan, W. Poewe, J. Jankovic, Parkinson disease and the immune system — associations, mechanisms and therapeutics. Nature Reviews Neurology. 16 (2020), pp. 303–318.

26. M. M. Hoehn, M. D. Yahr, Parkinsonism: onset, progression and mortality. Neurology. 17, 427–442 (1967).

27. A. B. Cuperfain, Z. L. Zhang, J. L. Kennedy, V. F. Gonçalves, The Complex Interaction of Mitochondrial Genetics and Mitochondrial Pathways in Psychiatric Disease. Mol Neuropsychiatry. 4, 52–69 (2018).

28. J. Roth, T. Vogl, C. Sorg, C. Sunderkötter, Phagocyte-specific S100 proteins: a novel group of proinflammatory molecules. Trends Immunol. 24, 155–158 (2003).

29. C. Xia, Z. Braunstein, A. C. Toomey, J. Zhong, X. Rao, S100 Proteins As an Important Regulator of Macrophage Inflammation. Front. Immunol. 8, 1908 (2017).

30. K. Sathe, W. Maetzler, J. D. Lang, R. B. Mounsey, C. Fleckenstein, H. L. Martin, C. Schulte, S. Mustafa, M. Synofzik, Z. Vukovic, S. Itohara, D. Berg, P. Teismann, S100B is increased in Parkinson’s disease and ablation protects against MPTP-induced toxicity through the RAGE and TNF-α pathway. Brain. 135, 3336–3347 (2012).

31. S. Jinn, R. E. Drolet, P. E. Cramer, A. H.-K. Wong, D. M. Toolan, C. A. Gretzula, B. Voleti, G. Vassileva, J. Disa, M. Tadin-Strapps, D. J. Stone, TMEM175 deficiency impairs lysosomal and mitochondrial function and increases α-synuclein aggregation. Proc. Natl. Acad. Sci. U. S. A. 114, 2389–2394 (2017).

32. K. Maiese, Z. Z. Chong, Y. C. Shang, S. Wang, mTOR: on target for novel therapeutic strategies in the nervous system. Trends Mol. Med. 19, 51–60 (2013).

33. J. Cao, M. Spielmann, X. Qiu, X. Huang, D. M. Ibrahim, A. J. Hill, F. Zhang, S. Mundlos, L. Christiansen, F. J. Steemers, C. Trapnell, J. Shendure, The single-cell transcriptional landscape of mammalian organogenesis. Nature. 566, 496–502 (2019).

34. H. K. Finucane, B. Bulik-Sullivan, A. Gusev, G. Trynka, Y. Reshef, P.-R. Loh, V. Anttila, H. Xu, C. Zang, K. Farh, S. Ripke, F. R. Day, ReproGen Consortium, Schizophrenia Working Group of the Psychiatric Genomics Consortium, RACI Consortium, S. Purcell, E. Stahl, S. Lindstrom, J. R. B. Perry, Y. Okada, S. Raychaudhuri, M. J. Daly, N. Patterson, B. M. Neale, A. L. Price, Partitioning heritability by functional annotation using genome-wide association summary statistics. Nat. Genet. 47, 1228–1235 (2015).

35. D. W. Yao, L. J. O’Connor, A. L. Price, A. Gusev, Quantifying genetic effects on disease mediated by assayed gene expression levels. Nat. Genet. (2020), doi: 10.1038/s41588-020-0625-2.

36. A. Young, N. Kumasaka, F. Calvert, T. R. Hammond, A map of transcriptional heterogeneity and regulatory variation in human microglia. bioRxiv (2019) (available at https://www.biorxiv.org/content/10.1101/2019.12.20.874099v1.abstract).

37. B. Ng, C. C. White, H.-U. Klein, S. K. Sieberts, C. McCabe, E. Patrick, J. Xu, L. Yu, C. Gaiteri, D. A. Bennett, S. Mostafavi, P. L. De Jager, An xQTL map integrates the genetic architecture of the human brain’s transcriptome and epigenome. Nat. Neurosci. 20, 1418–1426 (2017).

38. C. Giambartolomei, D. Vukcevic, E. E. Schadt, L. Franke, A. D. Hingorani, C. Wallace, V. Plagnol, Bayesian test for colocalisation between pairs of genetic association studies using summary statistics. PLoS Genet. 10, e1004383 (2014).

39. A. Nott, I. R. Holtman, N. G. Coufal, J. C. M. Schlachetzki, M. Yu, R. Hu, C. Z. Han, M. Pena, J. Xiao, Y. Wu, Z. Keulen, M. P. Pasillas, C. O’Connor, C. K. Nickl, S. T. Schafer, Z. Shen, R. A. Rissman, J. B. Brewer, D. Gosselin, D. D. Gonda, M. L. Levy, M. G. Rosenfeld, G. McVicker, F. H. Gage, B. Ren, C. K. Glass, Brain cell type-specific enhancer-promoter interactome maps and disease-risk association. Science. 366, 1134–1139 (2019).

40. M. S. Chattaragada, C. Riganti, M. Sassoe, M. Principe, M. M. Santamorena, C. Roux, C. Curcio, A. Evangelista, P. Allavena, R. Salvia, B. Rusev, A. Scarpa, P. Cappello, F. Novelli, FAM49B, a novel regulator of mitochondrial function and integrity that suppresses tumor metastasis. Oncogene. 37, 697–709 (2018).

41. S. Gordon, P. R. Taylor, Monocyte and macrophage heterogeneity. Nat. Rev. Immunol. 5, 953–964 (2005).

42. A. M. Zawada, K. S. Rogacev, B. Rotter, P. Winter, R.-R. Marell, D. Fliser, G. H. Heine, SuperSAGE evidence for CD14++CD16+ monocytes as a third monocyte subset. Blood. 118, e50–61 (2011).

43. R. Mukherjee, P. Kanti Barman, P. Kumar Thatoi, R. Tripathy, B. Kumar Das, B. Ravindran, Non-Classical monocytes display inflammatory features: Validation in Sepsis and Systemic Lupus Erythematous. Sci. Rep. 5, 13886 (2015).

44. V. Grozdanov, C. Bliederhaeuser, W. P. Ruf, V. Roth, K. Fundel-Clemens, L. Zondler, D. Brenner, A. Martin-Villalba, B. Hengerer, J. Kassubek, A. C. Ludolph, J. H. Weishaupt, K. M. Danzer, Inflammatory dysregulation of blood monocytes in Parkinson’s disease patients. Acta Neuropathol. 128, 651–663 (2014).

45. A. M. Smith, C. Depp, B. J. Ryan, G. I. Johnston, J. Alegre-Abarrategui, S. Evetts, M. Rolinski, F. Baig, C. Ruffmann, A. K. Simon, M. T. M. Hu, R. Wade-Martins, Mitochondrial dysfunction and increased glycolysis in prodromal and early Parkinson’s blood cells. Mov. Disord. 33, 1580–1590 (2018).

46. G.E. Hoffman, P. Rousos, Dream: Powerful differential expression analysis for repeated measures designs. bioRxiv (2020) (available at https://www.biorxiv.org/content/10.1101/432567v2.full).

47. Q. Wang, Y. Zhang, M. Wang, W.-M. Song, Q. Shen, A. McKenzie, I. Choi, X. Zhou, P.-Y. Pan, Z. Yue, B. Zhang, The landscape of multiscale transcriptomic networks and key regulators in Parkinson’s disease. Nat. Commun. 10, 5234 (2019).

48. A. H. Schapira, J. M. Cooper, D. Dexter, J. B. Clark, P. Jenner, C. D. Marsden, Mitochondrial complex I deficiency in Parkinson’s disease. J. Neurochem. 54, 823–827 (1990).

49. Y. Mizuno, S. Ohta, M. Tanaka, S. Takamiya, K. Suzuki, T. Sato, H. Oya, T. Ozawa, Y. Kagawa, Deficiencies in Complex I subunits of the respiratory chain in Parkinson’s disease. Biochemical and Biophysical Research Communications. 163 (1989), pp. 1450–1455.

50. J. M. Y. Teves, V. Bhargava, K. R. Kirwan, M. J. Corenblum, R. Justiniano, G. T. Wondrak, A. Anandhan, A. J. Flores, D. A. Schipper, Z. Khalpey, J. E. Sligh, C. Curiel-Lewandrowski, S. J. Sherman, L. Madhavan, Parkinson’s Disease Skin Fibroblasts Display Signature Alterations in Growth, Redox Homeostasis, Mitochondrial Function, and Autophagy. Front. Neurosci. 11, 737 (2017).

51. B. J. Ryan, S. Hoek, E. A. Fon, R. Wade-Martins, Mitochondrial dysfunction and mitophagy in Parkinson’s: from familial to sporadic disease. Trends Biochem. Sci. 40, 200–210 (2015).

52. P. Seibler, J. Graziotto, H. Jeong, F. Simunovic, C. Klein, D. Krainc, Mitochondrial Parkin recruitment is impaired in neurons derived from mutant PINK1 induced pluripotent stem cells. J. Neurosci. 31, 5970–5976 (2011).

53. H. Mortiboys, K. J. Thomas, W. J. H. Koopman, S. Klaffke, P. Abou-Sleiman, S. Olpin, N. W. Wood, P. H. Willems, J. A. M. Smeitink, M. R. Cookson, Others, Mitochondrial function and morphology are impaired in parkin-mutant fibroblasts. Annals of Neurology: Official Journal of the American Neurological Association and the Child Neurology Society. 64, 555–565 (2008).

54. H. Mortiboys, K. K. Johansen, J. O. Aasly, O. Bandmann, Mitochondrial impairment in patients with Parkinson disease with the G2019S mutation in LRRK2. Neurology. 75, 2017–2020 (2010).

55. S. K. Nissen, K. Shrivastava, C. Schulte, D. E. Otzen, D. Goldeck, D. Berg, H. J. Møller, W. Maetzler, M. Romero-Ramos, Alterations in Blood Monocyte Functions in Parkinson’s Disease. Mov. Disord. (2019), doi: 10.1002/mds.27815.

56. D. A. Cook, G. T. Kannarkat, A. F. Cintron, L. M. Butkovich, K. B. Fraser, J. Chang, N. Grigoryan, S. A. Factor, A. B. West, J. M. Boss, M. G. Tansey, LRRK2 levels in immune cells are increased in Parkinson’s disease. NPJ Parkinsons Dis. 3, 11 (2017).

57. S. J. Annesley, S. T. Lay, S. W. De Piazza, O. Sanislav, E. Hammersley, C. Y. Allan, L. M. Francione, M. Q. Bui, Z.-P. Chen, K. R. W. Ngoei, F. Tassone, B. E. Kemp, E. Storey, A. Evans, D. Z. Loesch, P. R. Fisher, Immortalized Parkinson’s disease lymphocytes have enhanced mitochondrial respiratory activity. Dis. Model. Mech. 9, 1295–1305 (2016).

58. P. Réu, A. Khosravi, S. Bernard, J. E. Mold, M. Salehpour, K. Alkass, S. Perl, J. Tisdale, G. Possnert, H. Druid, J. Frisén, The Lifespan and Turnover of Microglia in the Human Brain. Cell Rep. 20, 779–784 (2017).

59. J. Marschallinger, T. Iram, M. Zardeneta, S. E. Lee, B. Lehallier, M. S. Haney, J. V. Pluvinage, V. Mathur, O. Hahn, D. W. Morgens, J. Kim, J. Tevini, T. K. Felder, H. Wolinski, C. R. Bertozzi, M. C. Bassik, L. Aigner, T. Wyss-Coray, Lipid-droplet-accumulating microglia represent a dysfunctional and proinflammatory state in the aging brain. Nat. Neurosci. 23, 194–208 (2020).

60. L. Parnetti, L. Gaetani, P. Eusebi, S. Paciotti, O. Hansson, O. El-Agnaf, B. Mollenhauer, K. Blennow, P. Calabresi, CSF and blood biomarkers for Parkinson’s disease. Lancet Neurol. 18, 573–586 (2019).

61. L. Parnetti, A. Castrioto, D. Chiasserini, E. Persichetti, N. Tambasco, O. El-Agnaf, P. Calabresi, Cerebrospinal fluid biomarkers in Parkinson disease. Nature Reviews Neurology. 9 (2013), pp. 131–140.

62. S. E. Daniel, A. J. Lees, Parkinson’s Disease Society Brain Bank, London: overview and research. J. Neural Transm. Suppl. 39, 165–172 (1993).

63. Fahn, S, Members of the UPDRS Development Committee. Unified Parkinson’s Disease Rating Scale. Recent developments in Parkinson’s disease. 2, 293–304 (1987).

64. A. J. Hughes, Y. Ben-Shlomo, S. E. Daniel, A. J. Lees, UK Parkinson’s disease society brain bank clinical diagnostic criteria. J. Neurol. Neurosurg. Psychiatry. 55, e4 (1992).

65. S. Purcell, B. Neale, K. Todd-Brown, L. Thomas, M. A. R. Ferreira, D. Bender, J. Maller, P. Sklar, P. I. W. de Bakker, M. J. Daly, P. C. Sham, PLINK: a tool set for whole-genome association and population-based linkage analyses. Am. J. Hum. Genet. 81, 559–575 (2007).

66. S. Das, L. Forer, S. Schönherr, C. Sidore, A. E. Locke, A. Kwong, S. I. Vrieze, E. Y. Chew, S. Levy, M. McGue, D. Schlessinger, D. Stambolian, P.-R. Loh, W. G. Iacono, A. Swaroop, L. J. Scott, F. Cucca, F. Kronenberg, M. Boehnke, G. R. Abecasis, C. Fuchsberger, Next-generation genotype imputation service and methods. Nat. Genet. 48, 1284–1287 (2016).

67. Nextflow - A DSL for parallel and scalable computational pipelines, (available at https://www.nextflow.io/).

68. A. M. Bolger, M. Lohse, B. Usadel, Trimmomatic: a flexible trimmer for Illumina sequence data. Bioinformatics. 30, 2114–2120 (2014).

69. A. Dobin, C. A. Davis, F. Schlesinger, J. Drenkow, C. Zaleski, S. Jha, P. Batut, M. Chaisson, T. R. Gingeras, STAR: ultrafast universal RNA-seq aligner. Bioinformatics. 29, 15–21 (2013).

70. GENCODE - Human Release 30, (available at https://www.gencodegenes.org/human/release_30.html).

71. B. Li, C. N. Dewey, RSEM: accurate transcript quantification from RNA-Seq data with or without a reference genome. BMC Bioinformatics. 12, 323 (2011).

72. Y.-Y. Feng, A. Ramu, K. C. Cotto, Z. L. Skidmore, J. Kunisaki, D. F. Conrad, Y. Lin, W. C. Chapman, R. Uppaluri, R. Govindan, O. L. Griffith, M. Griffith, RegTools: Integrated analysis of genomic and transcriptomic data for discovery of splicing variants in cancer. bioRxiv (2018), p. 436634.

73. Babraham Bioinformatics - FastQC A Quality Control tool for High Throughput Sequence Data, (available at http://www.bioinformatics.babraham.ac.uk/projects/fastqc/).

74. Picard Tools - By Broad Institute, (available at https://broadinstitute.github.io/picard/).

75. H. Li, B. Handsaker, A. Wysoker, T. Fennell, J. Ruan, N. Homer, G. Marth, G. Abecasis, R. Durbin, 1000 Genome Project Data Processing Subgroup, The Sequence Alignment/Map format and SAMtools. Bioinformatics. 25, 2078–2079 (2009).

76. G.E. Hoffman, E.E. Schadt, variancePartition: Interpreting drivers of variation in complex gene expression studies. BMC Bioinformatics, 17, 483. (2016)

77. C. W. Law, Y. Chen, W. Shi, G. K. Smyth, voom: Precision weights unlock linear model analysis tools for RNA-seq read counts. Genome Biol. 15, R29 (2014).

78. M. E. Ritchie, B. Phipson, D. Wu, Y. Hu, C. W. Law, W. Shi, G. K. Smyth, limma powers differential expression analyses for RNA-sequencing and microarray studies. Nucleic Acids Res. 43, e47 (2015).

79. A. Subramanian, P. Tamayo, V. K. Mootha, S. Mukherjee, B. L. Ebert, M. A. Gillette, A. Paulovich, S. L. Pomeroy, T. R. Golub, E. S. Lander, J. P. Mesirov, Gene set enrichment analysis: a knowledge-based approach for interpreting genome-wide expression profiles. Proc. Natl. Acad. Sci. U. S. A. 102, 15545–15550 (2005).

80. A. V. Gomes, Genetics of proteasome diseases. Scientifica. 2013, 637629 (2013).

81. Y. I. Li, D. A. Knowles, J. Humphrey, A. N. Barbeira, S. P. Dickinson, H. K. Im, J. K. Pritchard, Annotation-free quantification of RNA splicing using LeafCutter. Nat. Genet. 50, 151–158 (2018).

82. Home | AMP-PD, (available at https://amp-pd.org/).

83. A. Buja, N. Eyuboglu, Remarks on Parallel Analysis. Multivariate Behav. Res. 27, 509–540 (1992).

84. P. Parsana, C. Ruberman, A. E. Jaffe, M. C. Schatz, A. Battle, J. T. Leek, Addressing confounding artifacts in reconstruction of gene co-expression networks. Genome Biol. 20, 94 (2019).

85. P. Langfelder, S. Horvath, WGCNA: an R package for weighted correlation network analysis. BMC Bioinformatics. 9 (2008),, doi: 10.1186/1471-2105-9-559.

86. Pedersen, T. L, ggraph. GitHub (2017), (available at https://github.com/thomasp85/ggraph).

87. B. W. Kunkle, B. Grenier-Boley, R. Sims, J. C. Bis, V. Damotte, A. C. Naj, A. Boland, M. Vronskaya, S. J. van der Lee, A. Amlie-Wolf, C. Bellenguez, A. Frizatti, V. Chouraki, E. R. Martin, K. Sleegers, N. Badarinarayan, J. Jakobsdottir, K. L. Hamilton-Nelson, S. Moreno-Grau, R. Olaso, R. Raybould, Y. Chen, A. B. Kuzma, M. Hiltunen, T. Morgan, S. Ahmad, B. N. Vardarajan, J. Epelbaum, P. Hoffmann, M. Boada, G. W. Beecham, J.-G. Garnier, D. Harold, A. L. Fitzpatrick, O. Valladares, M.-L. Moutet, A. Gerrish, A. V. Smith, L. Qu, D. Bacq, N. Denning, X. Jian, Y. Zhao, M. Del Zompo, N. C. Fox, S.-H. Choi, I. Mateo, J. T. Hughes, H. H. Adams, J. Malamon, F. Sanchez-Garcia, Y. Patel, J. A. Brody, B. A. Dombroski, M. C. D. Naranjo, M. Daniilidou, G. Eiriksdottir, S. Mukherjee, D. Wallon, J. Uphill, T. Aspelund, L. B. Cantwell, F. Garzia, D. Galimberti, E. Hofer, M. Butkiewicz, B. Fin, E. Scarpini, C. Sarnowski, W. S. Bush, S. Meslage, J. Kornhuber, C. C. White, Y. Song, R. C. Barber, S. Engelborghs, S. Sordon, D. Voijnovic, P. M. Adams, R. Vandenberghe, M. Mayhaus, L. A. Cupples, M. S. Albert, P. P. De Deyn, W. Gu, J. J. Himali, D. Beekly, A. Squassina, A. M. Hartmann, A. Orellana, D. Blacker, E. Rodriguez-Rodriguez, S. Lovestone, M. E. Garcia, R. S. Doody, C. Munoz-Fernadez, R. Sussams, H. Lin, T. J. Fairchild, Y. A. Benito, C. Holmes, H. Karamujic-Comic, M. P. Frosch, H. Thonberg, W. Maier, G. Roshchupkin, B. Ghetti, V. Giedraitis, A. Kawalia, S. Li, R. M. Huebinger, L. Kilander, S. Moebus, I. Hernández, M. I. Kamboh, R. Brundin, J. Turton, Q. Yang, M. J. Katz, L. Concari, J. Lord, A. S. Beiser, C. D. Keene, S. Helisalmi, I. Kloszewska, W. A. Kukull, A. M. Koivisto, A. Lynch, L. Tarraga, E. B. Larson, A. Haapasalo, B. Lawlor, T. H. Mosley, R. B. Lipton, V. Solfrizzi, M. Gill, W. T. Longstreth Jr, T. J. Montine, V. Frisardi, M. Diez-Fairen, F. Rivadeneira, R. C. Petersen, V. Deramecourt, I. Alvarez, F. Salani, A. Ciaramella, E. Boerwinkle, E. M. Reiman, N. Fievet, J. I. Rotter, J. S. Reisch, O. Hanon, C. Cupidi, A. G. Andre Uitterlinden, D. R. Royall, C. Dufouil, R. G. Maletta, I. de Rojas, M. Sano, A. Brice, R. Cecchetti, P. S. George-Hyslop, K. Ritchie, M. Tsolaki, D. W. Tsuang, B. Dubois, D. Craig, C. -Wu, H. Soininen, D. Avramidou, R. L. Albin, L. Fratiglioni, A. Germanou, L. G. Apostolova, Keller, M. Koutroumani, S. E. Arnold, F. Panza, O. Gkatzima, S. Asthana, D. Hannequin, P. Whitehead, C. S. Atwood, P. Caffarra, H. Hampel, I. Quintela, Á. Carracedo, L. Lannfelt, D. C. Rubinsztein, L. L. Barnes, F. Pasquier, L. Frölich, S. Barral, B. McGuinness, T. G. Beach, J. A. Johnston, J. T. Becker, P. Passmore, E. H. Bigio, J. M. Schott, T. D. Bird, J. D. Warren, B. F. Boeve, M. K. Lupton, J. D. Bowen, P. Proitsi, A. Boxer, J. F. Powell, J. R. Burke, J. S. K. Kauwe, J. M. Burns, M. Mancuso, J. D. Buxbaum, U. Bonuccelli, N. J. Cairns, A. McQuillin, C. Cao, G. Livingston, C. S. Carlson, N. J. Bass, C. M. Carlsson, J. Hardy, R. M. Carney, J. Bras, M. M. Carrasquillo, R. Guerreiro, M. Allen, H. C. Chui, E. Fisher, C. Masullo, E. A. Crocco, C. DeCarli, G. Bisceglio, M. Dick, L. Ma, R. Duara, N. R. Graff-Radford, D. A. Evans, A. Hodges, K. M. Faber, M. Scherer, K. B. Fallon, M. Riemenschneider, D. W. Fardo, R. Heun, M. R. Farlow, H. Kölsch, S. Ferris, M. Leber, T. M. Foroud, I. Heuser, D. R. Galasko, I. Giegling, M. Gearing, M. Hüll, D. H. Geschwind, J. R. Gilbert, J. Morris, R. C. Green, K. Mayo, J. H. Growdon, T. Feulner, R. L. Hamilton, L. E. Harrell, D. Drichel, L. S. Honig, T. D. Cushion, M. J. Huentelman, P. Hollingworth, C. M. Hulette, B. T. Hyman, R. Marshall, G. P. Jarvik, A. Meggy, E. Abner, G. E. Menzies, L.-W. Jin, G. Leonenko, L. M. Real, G. R. Jun, C. T. Baldwin, D. Grozeva, A. Karydas, G. Russo, J. A. Kaye, R. Kim, F. Jessen, N. W. Kowall, B. Vellas, J. H. Kramer, E. Vardy, F. M. LaFerla, K.-H. Jöckel, J. J. Lah, M. Dichgans, J. B. Leverenz, D. Mann, A. I. Levey, S. Pickering-Brown, A. P. Lieberman, N. Klopp, K. L. Lunetta, H.-E. Wichmann, C. G. Lyketsos, K. Morgan, D. C. Marson, K. Brown, F. Martiniuk, C. Medway, D. C. Mash, M. M. Nöthen, E. Masliah, N. M. Hooper, W. C. McCormick, A. Daniele, S. M. McCurry, A. Bayer, A. N. McDavid, J. Gallacher, A. C. McKee, H. van den Bussche, M. Mesulam, C. Brayne, B. L. Miller, S. Riedel-Heller, C. A. Miller, J. W. Miller, A. Al-Chalabi, J. C. Morris, C. E. Shaw, A. J. Myers, J. Wiltfang, S. O’Bryant, J. M. Olichney, V. Alvarez, J. E. Parisi, A. B. Singleton, H. L. Paulson, J. Collinge, W. R. Perry, S. Mead, E. Peskind, D. H. Cribbs, M. Rossor, A. Pierce, N. S. Ryan, W. W. Poon, B. Nacmias, H. Potter, S. Sorbi, J. F. Quinn, E. Sacchinelli, A. Raj, G. Spalletta, M. Raskind, C. Caltagirone, P. Bossù, M. D. Orfei, B. Reisberg, R. Clarke, C. Reitz, A. D. Smith, J. M. Ringman, D. Warden, E. D. Roberson, G. Wilcock, E. Rogaeva, A. C. Bruni, H. J. Rosen, M. Gallo, R. N. Rosenberg, Y. Ben-Shlomo, M. A. Sager, P. Mecocci, A. J. Saykin, P. Pastor, M. L. Cuccaro, J. M. Vance, J. A. Schneider, L. S. Schneider, S. Slifer, W. W. Seeley, A. G. Smith, J. A. Sonnen, S. Spina, R. A. Stern, R. H. Swerdlow, M. Tang, R. E. Tanzi, J. Q. Trojanowski, J. C. Troncoso, V. M. Van Deerlin, L. J. Van Eldik, H. V. Vinters, J. P. Vonsattel, S. Weintraub, K. A. Welsh-Bohmer, K. C. Wilhelmsen, J. Williamson, T. S. Wingo, R. L. Woltjer, C. B. Wright, C.-E. Yu, L. Yu, Y. Saba, A. Pilotto, M. J. Bullido, O. Peters, P. K. Crane, D. Bennett, P. Bosco, E. Coto, V. Boccardi, P. L. De Jager, A. Lleo, N. Warner, O. L. Lopez, M. Ingelsson, P. Deloukas, C. Cruchaga, C. Graff, R. Gwilliam, M. Fornage, A. M. Goate, P. Sanchez-Juan, P. G. Kehoe, N. Amin, N. Ertekin-Taner, C. Berr, S. Debette, S. Love, L. J. Launer, S. G. Younkin, J.-F. Dartigues, C. Corcoran, M. A. Ikram, D. W. Dickson, G. Nicolas, D. Campion, J. Tschanz, H. Schmidt, H. Hakonarson, J. Clarimon, R. Munger, R. Schmidt, L. A. Farrer, C. Van Broeckhoven, M. C O’Donovan, A. L. DeStefano, L. Jones, J. L. Haines, J.-F. Deleuze, M. J. Owen, V. Gudnason, R. Mayeux, V. Escott-Price, B. M. Psaty, A. Ramirez, L.-S. Wang, A. Ruiz, C. M. van Duijn, P. A. Holmans, S. Seshadri, J. Williams, P. Amouyel, G. D. Schellenberg, J.-C. Lambert, M. A. Pericak-Vance, Alzheimer Disease Genetics Consortium (ADGC), European Alzheimer’s Disease Initiative (EADI), Cohorts for Heart and Aging Research in Genomic Epidemiology Consortium (CHARGE),, Genetic and Environmental Risk in AD/Defining Genetic, Polygenic and Environmental Risk for Alzheimer’s Disease Consortium (GERAD/PERADES), Genetic meta-analysis of diagnosed Alzheimer’s disease identifies new risk loci and implicates Aβ, tau, immunity and lipid processing. Nat. Genet. 51, 414–430 (2019).

88. Schizophrenia Working Group of the Psychiatric Genomics Consortium, Biological insights from 108 schizophrenia-associated genetic loci. Nature. 511, 421–427 (2014).

89. A. R. Wood, T. Esko, J. Yang, S. Vedantam, T. H. Pers, S. Gustafsson, A. Y. Chu, K. Estrada, J. ‘an Luan, Z. Kutalik, N. Amin, M. L. Buchkovich, D. C. Croteau-Chonka, F. R. Day, Y. Duan, T. Fall, R. Fehrmann, T. Ferreira, A. U. Jackson, J. Karjalainen, K. S. Lo, A. E. Locke, R. Mägi, E. Mihailov, E. Porcu, J. C. Randall, A. Scherag, A. A. E. Vinkhuyzen, H.-J. Westra, T. W. Winkler, T. Workalemahu, J. H. Zhao, D. Absher, E. Albrecht, D. Anderson, J. Baron, M. Beekman, A. Demirkan, G. B. Ehret, B. Feenstra, M. F. Feitosa, K. Fischer, R. M. Fraser, A. Goel, J. Gong, A. E. Justice, S. Kanoni, M. E. Kleber, K. Kristiansson, U. Lim, V. Lotay, J. C. Lui, M. Mangino, I. Mateo Leach, C. Medina-Gomez, M. A. Nalls, D. R. Nyholt, C. D. Palmer, D. Pasko, S. Pechlivanis, I. Prokopenko, J. S. Ried, S. Ripke, D. Shungin, A. Stancáková, R. J. Strawbridge, Y. J. Sung, T. Tanaka, A. Teumer, S. Trompet, S. W. van der Laan, J. van Setten, J. V. Van Vliet-Ostaptchouk, Z. Wang, L. Yengo, W. Zhang, U. Afzal, J. Arnlöv, G. M. Arscott, S. Bandinelli, A. Barrett, C. Bellis, A. J. Bennett, C. Berne, M. Blüher, J. L. Bolton, Y. Böttcher, H. A. Boyd, M. Bruinenberg, B. M. Buckley, S. Buyske, I. H. Caspersen, P. S. Chines, R. Clarke, S. Claudi-Boehm, M. Cooper, E. W. Daw, P. A. De Jong, J. Deelen, G. Delgado, J. C. Denny, R. Dhonukshe-Rutten, M. Dimitriou, A. S. F. Doney, M. Dörr, N. Eklund, E. Eury, L. Folkersen, M. E. Garcia, F. Geller, V. Giedraitis, A. S. Go, H. Grallert, T. B. Grammer, J. Gräßler, H. Grönberg, L. C. P. G. M. de Groot, C. J. Groves, J. Haessler, P. Hall, T. Haller, G. Hallmans, A. Hannemann, C. A. Hartman, M. Hassinen, C. Hayward, N. L. Heard-Costa, Q. Helmer, G. Hemani, A. K. Henders, H. L. Hillege, M. A. Hlatky, W. Hoffmann, P. Hoffmann, O. Holmen, J. J. Houwing-Duistermaat, T. Illig, A. Isaacs, A. L. James, J. Jeff, B. Johansen, Å. Johansson, J. Jolley, T. Juliusdottir, J. Junttila, A. N. Kho, L. Kinnunen, N. Klopp, T. Kocher, W. Kratzer, P. Lichtner, L. Lind, J. Lindström, S. Lobbens, M. Lorentzon, Y. Lu, V. Lyssenko, P. K. E. Magnusson, A. Mahajan, M. Maillard, W. L. McArdle, C. A. McKenzie, S. McLachlan, P. J. McLaren, C. Menni, S. Merger, L. Milani, A. Moayyeri, K. L. Monda, M. A. Morken, G. Müller, M. Müller-Nurasyid, A. W. Musk, N. Narisu, M. Nauck, I. M. Nolte, M. M. Nöthen, L. Oozageer, S. Pilz, N. W. Rayner, F. Renstrom, N. R. Robertson, L. M. Rose, R. Roussel, S. Sanna, H. Scharnagl, S. Scholtens, F. R. Schumacher, H. Schunkert, R. A. Scott, J. Sehmi, T. Seufferlein, J. Shi, K. Silventoinen, J. H. Smit, A. V. Smith, J. Smolonska, A. V. Stanton, K. Stirrups, D. J. Stott, H. M. Stringham, J. Sundström, M. A. Swertz, A.-C. Syvänen, B. O. Tayo, G. Thorleifsson, J. P. Tyrer, S. van Dijk, N. M. van Schoor, N. van der Velde, D. van Heemst, F. V. A. van Oort, S. H. Vermeulen, N. Verweij, J. M. Vonk, L. L. Waite, M. Waldenberger, R. Wennauer, L. R. Wilkens, C. Willenborg, T. Wilsgaard, M. K. Wojczynski, A. Wong, A. F. Wright, Q. Zhang, D. Arveiler, S. J. L. Bakker, J. Beilby, R. N. Bergman, S. Bergmann, R. Biffar, J. Blangero, D. I. Boomsma, S. R. Bornstein, P. Bovet, P. Brambilla, M. J. Brown, H. Campbell, M. J. Caulfield, A. Chakravarti, R. Collins, F. S. Collins, D. C. Crawford, L. A. Cupples, J. Danesh, U. de Faire, H. M. den Ruijter, R. Erbel, J. Erdmann, J. G. Eriksson, M. Farrall, E. Ferrannini, J. Ferrières, I. Ford, N. G. Forouhi, T. Forrester, R. T. Gansevoort, P. V. Gejman, C. Gieger, A. Golay, O. Gottesman, V. Gudnason, U. Gyllensten, D. W. Haas, A. S. Hall, T. B. Harris, A. T. Hattersley, A. C. Heath, C. Hengstenberg, A. A. Hicks, L. A. Hindorff, A. D. Hingorani, A. Hofman, G. K. Hovingh, S. E. Humphries, S. C. Hunt, E. Hypponen, K. B. Jacobs, M.-R. Jarvelin, P. Jousilahti, A. M. Jula, J. Kaprio, J. J. P. Kastelein, M. Kayser, F. Kee, S. M. Keinanen-Kiukaanniemi, L. A. Kiemeney, J. S. Kooner, C. Kooperberg, S. Koskinen, P. Kovacs, A. T. Kraja, M. Kumari, J. Kuusisto, T. A. Lakka, C. Langenberg, L. Le Marchand, T. Lehtimäki, S. Lupoli, P. A. F. Madden, S. Männistö, P. Manunta, A. Marette, T. C. Matise, B. McKnight, T. Meitinger, F. L. Moll, G. W. Montgomery, A. D. Morris, A. P. Morris, J. C. Murray, M. Nelis, C. Ohlsson, A. J. Oldehinkel, K. K. Ong, W. H. Ouwehand, G. Pasterkamp, A. Peters, P. P. Pramstaller, J. F. Price, L. Qi, O. T. Raitakari, T. Rankinen, D. C. Rao, T. K. Rice, M. Ritchie, I. Rudan, V. Salomaa, N. J. Samani, J. Saramies, M. A. Sarzynski, P. E. H. Schwarz, S. Sebert, P. Sever, A. R. Shuldiner, J. Sinisalo, V. Steinthorsdottir, R. P. Stolk, J.-C. Tardif, A. Tönjes, A. Tremblay, E. Tremoli, J. Virtamo, M.-C. Vohl, Electronic Medical Records and Genomics (eMEMERGEGE) Consortium, MIGen Consortium, PAGEGE Consortium, LifeLines Cohort Study, P. Amouyel, F. W. Asselbergs, T. L. Assimes, M. Bochud, B. O. Boehm, E. Boerwinkle, E. P. Bottinger, C. Bouchard, S. Cauchi, J. C. Chambers, S. J. Chanock, R. S. Cooper, P. I. W. de Bakker, G. Dedoussis, L. Ferrucci, P. W. Franks, P. Froguel, L. C. Groop, C. A. Haiman, A. Hamsten, M. G. Hayes, J. Hui, D. J. Hunter, K. Hveem, J. W. Jukema, R. C. Kaplan, M. Kivimaki, D. Kuh, M. Laakso, Y. Liu, N. G. Martin, W. März, M. Melbye, S. Moebus, P. B. Munroe, I. Njølstad, B. A. Oostra, C. N. A. Palmer, N. L. Pedersen, M. Perola, L. Pérusse, U. Peters, J. E. Powell, C. Power, T. Quertermous, R. Rauramaa, E. Reinmaa, P. M. Ridker, F. Rivadeneira, J. I. Rotter, T. E. Saaristo, D. Saleheen, D. Schlessinger, P. E. Slagboom, H. Snieder, T. D. Spector, K. Strauch, M. Stumvoll, J. Tuomilehto, M. Uusitupa, P. van der Harst, H. Völzke, M. Walker, N. J. Wareham, H. Watkins, H.-E. Wichmann, J. F. Wilson, P. Zanen, P. Deloukas, I. M. Heid, C. M. Lindgren, K. L. Mohlke, E. K. Speliotes, U. Thorsteinsdottir, I. Barroso, C. S. Fox, K. E. North, D. P. Strachan, J. S. Beckmann, S. I. Berndt, M. Boehnke, I. B. Borecki, M. I. McCarthy, A. Metspalu, K. Stefansson, A. G. Uitterlinden, C. M. van Duijn, L. Franke, C. J. Willer, A. L. Price, G. Lettre, R. J. F. Loos, M. N. Weedon, E. Ingelsson, J. R. O’Connell, G. R. Abecasis, D. I. Chasman, M. E. Goddard, P. M. Visscher, J. N. Hirschhorn, T. M. Frayling, Defining the role of common variation in the genomic and biological architecture of adult human height. Nat. Genet. 46, 1173–1186 (2014).

90. *gtex-pipeline* (Github; https://github.com/broadinstitute/gtex-pipeline).

91. O. Stegle, L. Parts, M. Piipari, J. Winn, R. Durbin, Using probabilistic estimation of expression residuals (PEER) to obtain increased power and interpretability of gene expression analyses. Nat. Protoc. 7, 500–507 (2012).

92. O. Delaneau, H. Ongen, A. Brown, A. Fort, N. Panousis, E. Dermitzakis, A complete tool set for molecular QTL discovery and analysis. Nature Communications. 8, 15452. (2017).

93. B. P. Fairfax, P. Humburg, S. Makino, V. Naranbhai, D. Wong, E. Lau, L. Jostins, K. Plant, R. Andrews, C. McGee, J. C. Knight, Innate immune activity conditions the effect of regulatory variants upon monocyte gene expression. Science. 343, 1246949 (2014).

94. F. Aguet, A. N. Barbeira, R. Bonazzola, A. Brown, S. E. Castel, B. Jo, S. Kasela, S. Kim-Hellmuth, Y. Liang, M. Oliva, P. E. Parsana, E. Flynn, L. Fresard, E. R. Gaamzon, A. R. Hamel, Y. He, F. Hormozdiari, P. Mohammadi, M. Muñoz-Aguirre, Y. Park, A. Saha, A. V. Segrc, B. J. Strober, X. Wen, V. Wucher, S. Das, D. Garrido-Martín, N. R. Gay, R. E. Handsaker, P. J. Hoffman, S. Kashin, A. Kwong, X. Li, D. MacArthur, J. M. Rouhana, M. Stephens, E. Todres, A. Viñuela, G. Wang, Y. Zou, The GTEx Consortium, C. D. Brown, N. Cox, E. Dermitzakis, B. E. Engelhardt, G. Getz, R. Guigo, S. B. Montgomery, B. E. Stranger, H. K. Im, A. Battle, K. G. Ardlie, T. Lappalainen, The GTEx Consortium atlas of genetic regulatory effects across human tissues. bioRxiv (2019) (available at https://www.biorxiv.org/content/10.1101/787903v1)

95. summary-gwas-imputation (Github; https://github.com/RajLabMSSM/summary-gwas-imputation).

96. HaploReg v4.1, (available at https://pubs.broadinstitute.org/mammals/haploreg/haploreg.php).

97. O. Weissbrod, F. Hormozdiari, C. Benner, R. Cui, J. Ulirsch, S. Gazal, A. P. Schoech, B. van de Geijn, Y. Reshef, C. Márquez-Luna, L. O’Connor, M. Pirinen, H. K. Finucane, A. L. Price, Functionally-informed fine-mapping and polygenic localization of complex trait heritability. BioRxiv (2019) (available at https://www.biorxiv.org/content/10.1101/807792v3)

98. G. Wang, A. Sarkar, P. Carbonetto, M. Stephens, A simple new approach to variable selection in regression, with application to genetic fine-mapping (2018) (available at https://www.biorxiv.org/content/10.1101/501114v1)

99. S. Gazal, H. K. Finucane, N. A. Furlotte, P.-R. Loh, P. F. Palamara, X. Liu, A. Schoech, B. Bulik-Sullivan, B. M. Neale, A. Gusev, A. L. Price, Linkage disequilibrium–dependent architecture of human complex traits shows action of negative selection. Nature Genetics. 49 (2017), pp. 1421–1427.

100. B. K. Bulik-Sullivan, Schizophrenia Working Group of the Psychiatric Genomics Consortium, P.-R. Loh, H. K. Finucane, S. Ripke, J. Yang, N. Patterson, M. J. Daly, A. L. Price, B. M. Neale, LD Score regression distinguishes confounding from polygenicity in genome-wide association studies. Nature Genetics. 47 (2015), pp. 291–295.

101. S. Gazal, P.-R. Loh, H. K. Finucane, A. Ganna, A. Schoech, S. Sunyaev, A. L. Price, Functional architecture of low-frequency variants highlights strength of negative selection across coding and non-coding annotations. Nature Genetics. 50 (2018), pp. 1600–1607.

102. M. Stoeckius, S. Zheng, B. Houck-Loomis, S. Hao, B. Z. Yeung, W. M. Mauck 3rd, P. Smibert, R. Satija, Cell Hashing with barcoded antibodies enables multiplexing and doublet detection for single cell genomics. Genome Biol. 19, 224 (2018).

103. A. Butler, P. Hoffman, P. Smibert, E. Papalexi, R. Satija, Integrating single-cell transcriptomic data across different conditions, technologies, and species. Nat. Biotechnol. 36, 411–420 (2018).

104. T. Stuart, A. Butler, P. Hoffman, C. Hafemeister, E. Papalexi, W. M. Mauck 3rd, Y. Hao, M. Stoeckius, P. Smibert, R. Satija, Comprehensive Integration of Single-Cell Data. Cell. 177, 1888–1902.e21 (2019).

105. S. Durinck, Y. Moreau, A. Kasprzyk, S. Davis, B. De Moor, A. Brazma, W. Huber, BioMart and Bioconductor: a powerful link between biological databases and microarray data analysis. Bioinformatics. 21, 3439–3440 (2005).

106. E. Becht, L. McInnes, J. Healy, C.-A. Dutertre, I. W. H. Kwok, L. G. Ng, F. Ginhoux, E. W. Newell, Dimensionality reduction for visualizing single-cell data using UMAP. Nat. Biotechnol. (2018), doi: 10.1038/nbt.4314.

107. L. McInnes, J. Healy, J. Melville, UMAP: Uniform Manifold Approximation and Projection for Dimension Reduction. arXiv [stat.ML] (2018), (available at http://arxiv.org/abs/1802.03426).

108. Y. Benjamini, Y. Hochberg, Controlling the False Discovery Rate: A Practical and Powerful Approach to Multiple Testing. J. R. Stat. Soc. Series B Stat. Methodol. 57, 289–300 (1995).

109. J. Melief, M. A. M. Sneeboer, M. Litjens, P. R. Ormel, S. J. M. C. Palmen, I. Huitinga, R. S. Kahn, E. M. Hol, L. D. de Witte, Characterizing primary human microglia: A comparative study with myeloid subsets and culture models. Glia. 64, 1857–1868 (2016).

110. O. Weissbrod, F. Hormozdiari, C. Benner, R. Cui, Functionally-informed fine-mapping and polygenic localization of complex trait heritability. BioRxiv (2019) (available at https://www.biorxiv.org/content/10.1101/807792v1.abstract).

111. C. Trapnell, D. Cacchiarelli, J. Grimsby, P. Pokharel, S. Li, M. Morse, N. J. Lennon, K. J. Livak, T. S. Mikkelsen, J. L. Rinn, The dynamics and regulators of cell fate decisions are revealed by pseudotemporal ordering of single cells. Nat. Biotechnol. 32, 381–386 (2014).

