## Supplementary Figures 1-14 for "Discordant transcriptional signatures of mitochondrial genes in Parkinson’s disease human myeloid cells"

**Fig. S1.**

A

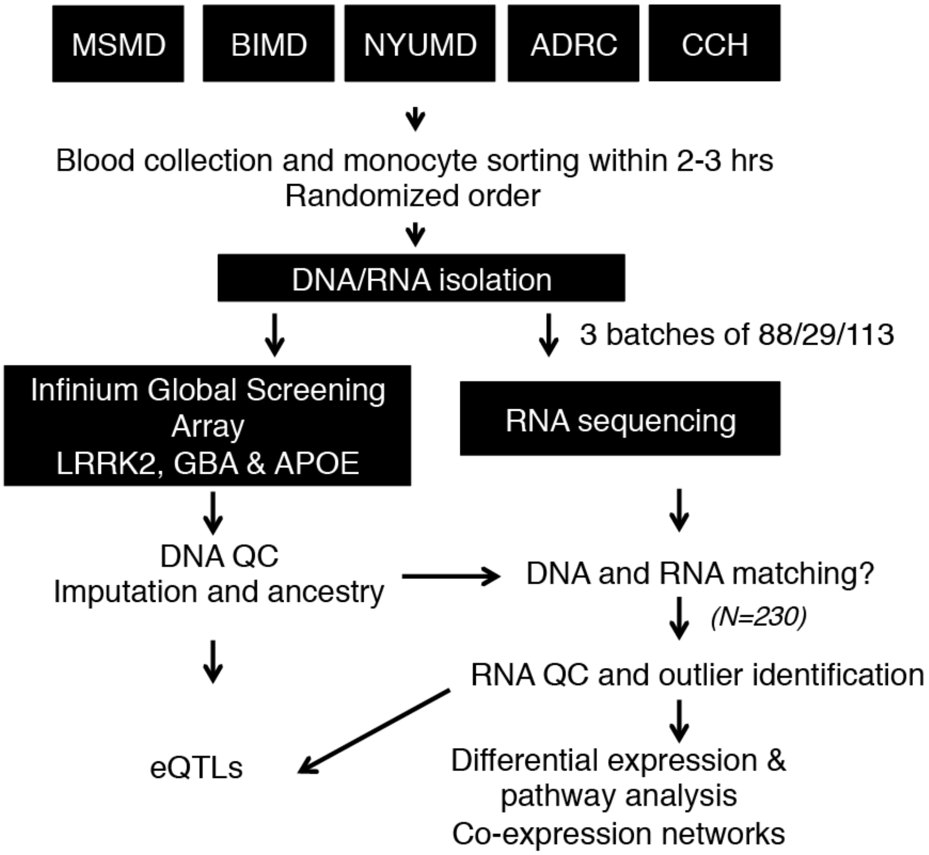


B

| Diagnosis | Samples | Age | Gender | | Genotype | | Population | | | | | AJ |
|---|---|---|---|---|---|---|---|---|---|---|---|---|
| | | | M | F | LRRK2 | GBA | EUR | AFR | AMR | SAS | NA | |
| **Control** | 95 | 68 | 30 | 65 | 3 | 8 | 75 | 4 | 9 | 0 | 5 | 24 |
| **PD** | 135 | 66 | 86 | 49 | 19 | 27 | 113 | 0 | 6 | 1 | 15 | 58 |

C

| Diagnosis | Samples | Age of onset | Disease duration | L-DOPA | Family history PD | H&R (mean) | UPDRS III (mean) | MOCA (mean) |
|---|---|---|---|---|---|---|---|---|
| PD | 135 | 57.3 | 8.3 | 79 % | 44 % | 1.8 | 15.5 | 26.6 |

**Fig. S1**

**Supplementary Figure 1. Experimental flow outline and demographic/clinical information for subjects for monocytes isolation. (A)** Blood was collected from five independent clinics across New York City (ADRC, CCH, MSMD, BIMD, and NYUMD; details described in Methods) and transferred to the Icahn School of Medicine at Mount Sinai for monocyte sorting and RNA/DNA isolation. Samples were genotyped for common SNPs using Global Screening Array (GSA) and LRRK2, GBA and APOE were independently genotyped. RNA-seq was performed at Genewiz Inc. in three independent and randomized batches. DNA and RNA data was subjected to stringent QC, DNA data was imputed and ancestry was calculated. DNA and RNA were compared to the identification of miss-matches prior outlier identification. After QC, a total of 230 samples were used for subsequent analysis. **(B)** Demographic and **(C)** clinical variables describing the 230 samples included in the study.

Fig. S2.

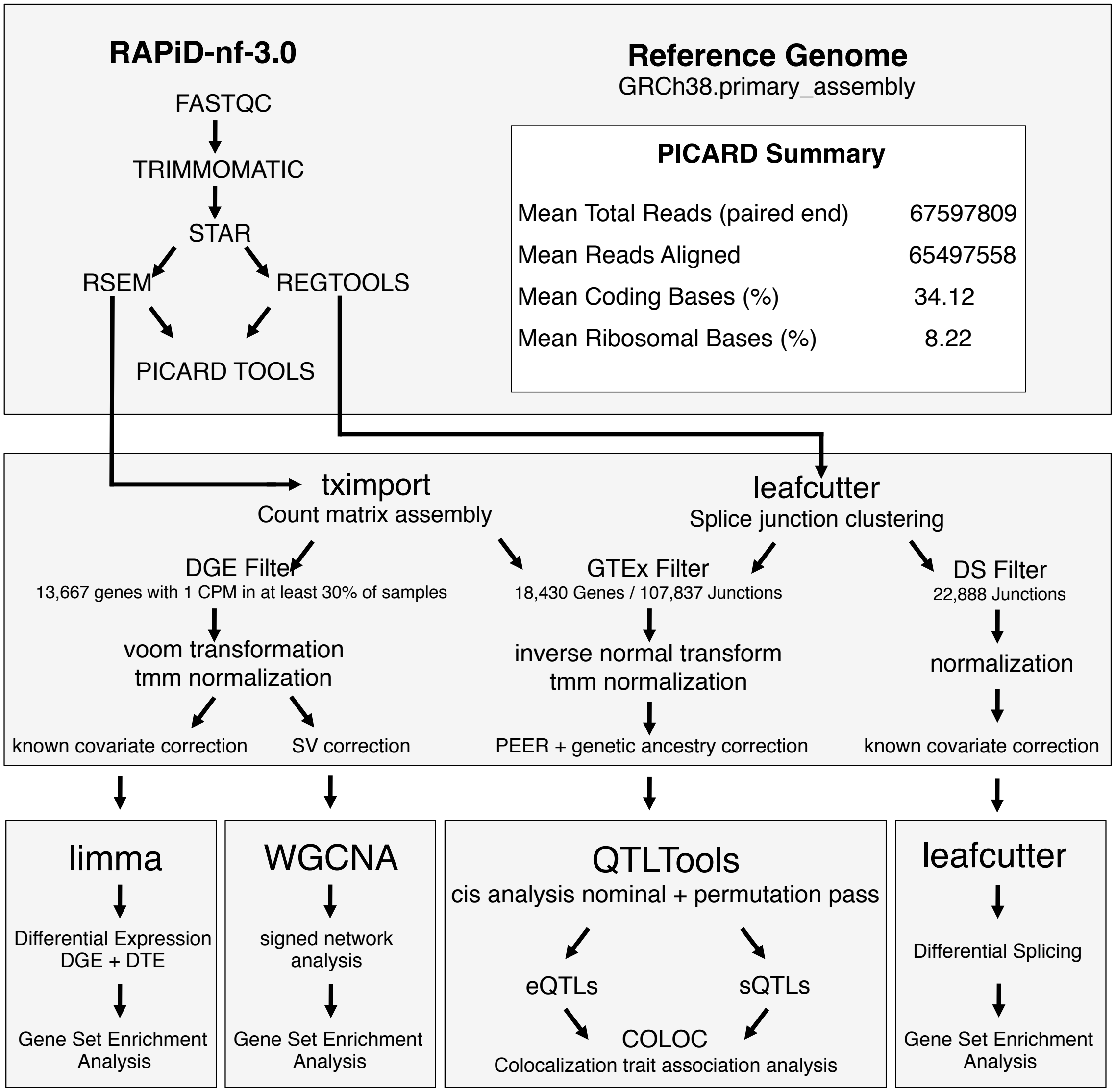


**Fig. S2**

**Supplementary Figure 2. Diagram representing the computational approach used in this study. (A)** Overview of RNA-seq analysis pipeline. RAPiD pipeline of mapping, QC, and quantification. **(B)** Normalization, covariate selection, and outlier removal. **(C)** Downstream analyses plan for differential expression, splicing, network analysis, and quantitative trait loci analyses.

**Fig. S3.**

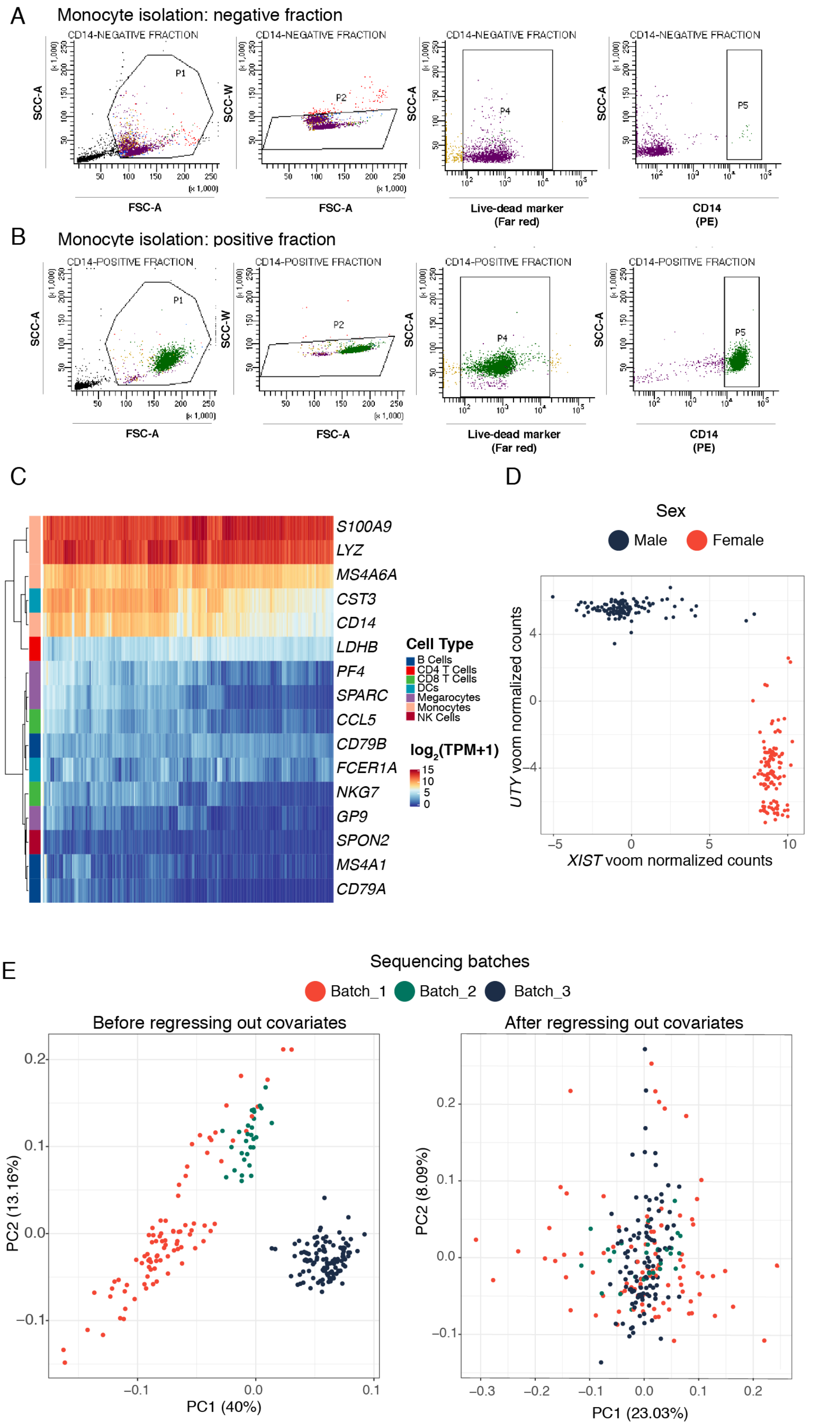
A
Monocyte isolation: negative fraction
CD14-NEGATIVE FRACTION
SCC-A
SCC-W
FSC-A
Live-dead marker
(Far red)
CD14
(PE)
P1
P2
P4
P5
B
Monocyte isolation: positive fraction
CD14-POSITIVE FRACTION
C
S100A9
LYZ
MS4A6A
CST3
CD14
LDHB
PF4
SPARC
CCL5
CD79B
FCER1A
NKG7
GP9
SPON2
MS4A1
CD79A
Cell Type
B Cells
CD4 T Cells
CD8 T Cells
DCs
Megarocytes
Monocytes
NK Cells
log2(TPM+1)
D
Sex
Male
Female
UTY voom normalized counts
XIST voom normalized counts
E
Sequencing batches
Batch_1
Batch_2
Batch_3
Before regressing out covariates
After regressing out covariates
PC2 (13.16%)
PC1 (40%)
PC2 (8.09%)
PC1 (23.03%)

**Fig. S3**

**Supplementary Figure 3. Purity of monocyte isolation and dimensionality reduction.** FACS plots of the **(A)** negative and **(B)** positive fractions after CD14-magnetic selection. Purity of the monocyte was 97.8%. **(C)** Heatmap representing in the y-axis the cell markers corresponding to different blood immune cells and the 230 samples in the x-axis. Color corresponds to the levels of expression (TPM + 1, $\log_2$ scale), blue representing low, and red meaning high. **(D)** Sex mismatch QC: scatter plot showing the voom transformed expression of *XIST* (X chromosome gene) in the x-axis and *UTY* (Y chromosome gene) in the y-axis. Each dot represents a sample and is colored by reported sex (red = female; blue = male). **(E)** PCA plots showing the expression (TPMs) of the 230 samples before (left panel) and after (right panel) regressing out known covariates. Each dot represents a sample and is colored by sequencing batches (orange = batch_1, green = batch_2, blue = batch_3). Each batch has an equal proportion of case and control subjects.

**Fig. S4.**

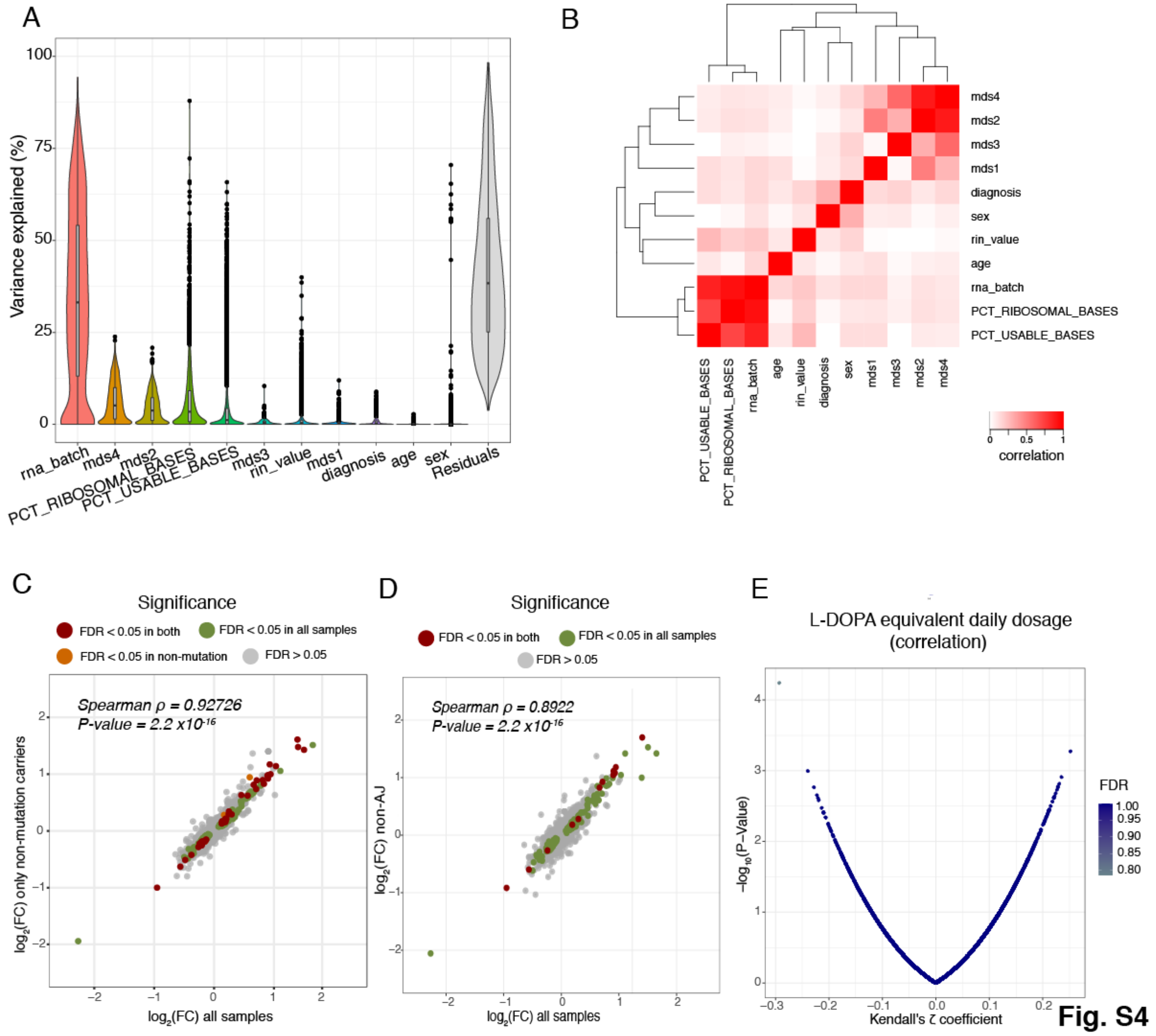


**Supplementary Figure 4. Covariate selection and correlation with possible confounders.** Main sources of variation were analyzed using the variancePartition package (76). **(A)** Violin plot showing the % of the variance (y-axis) explained by the covariates (x-axis) which contribute the most to the variability. **(B)** Heatmap showing the correlation among the different covariates (red represents high correlation vs white low correlation). **(C)** Scatter plot representing fold-change correlation transcriptome-wide when DE analysis was performed including all samples (n = 230) (x-axis) versus including only samples which are not *GBA* or *LRRK2* mutation carriers (n = 170) (y-axis). **(D)** Scatter plot representing fold-change correlation transcriptome-wide when DE analysis was performed including all samples (n = 230) (x-axis) versus only samples with non-AJ ancestry (n = 148) (y-axis). **(E)** Correlation of Levodopa equivalent daily dosage (LEDD) with gene expression levels for all genes tested in differential expression analysis.

Fig. S5.

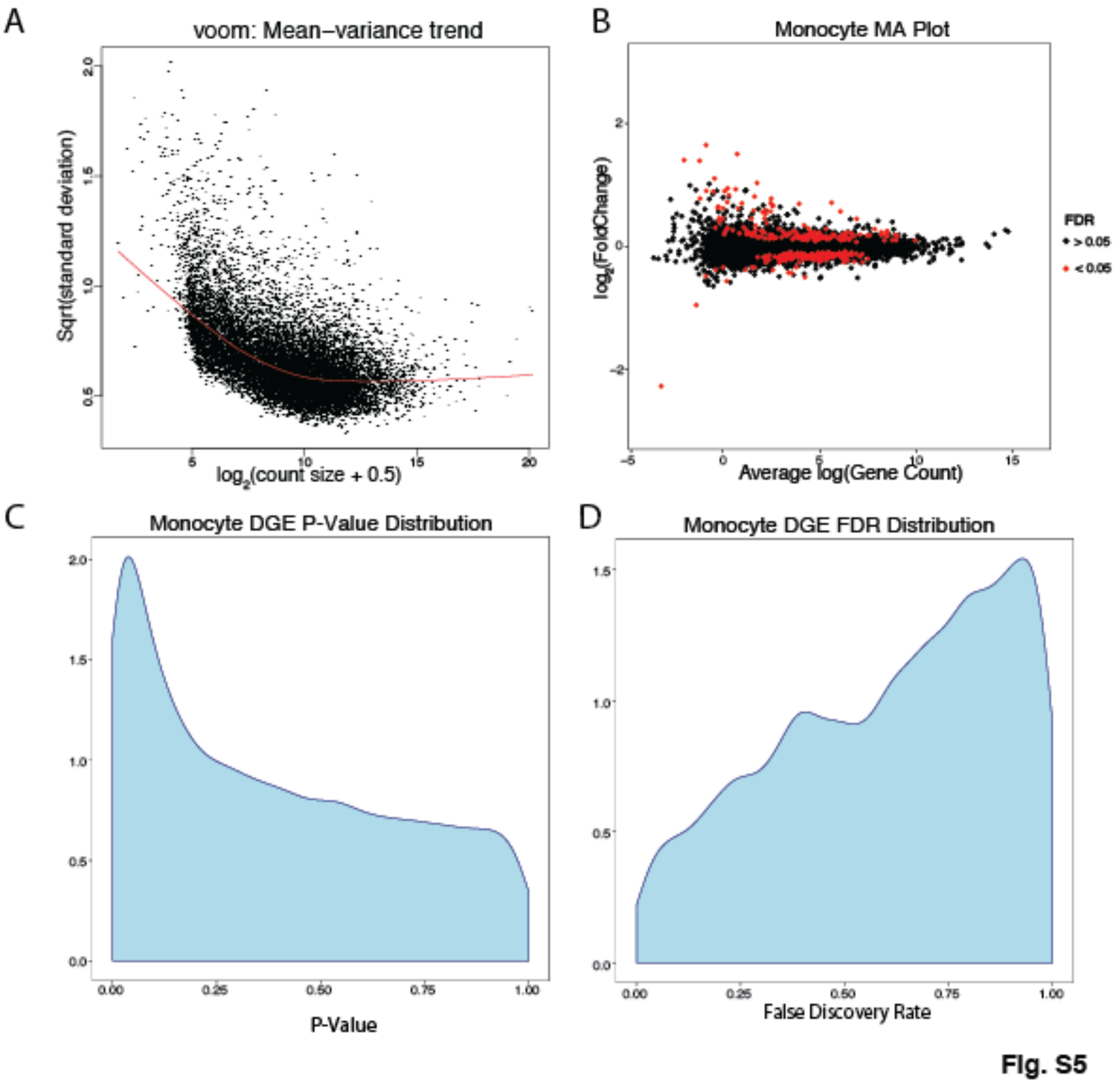


**Supplementary Figure 5. RNA-seq normalization and *P-value* distribution of differential expression.** **(A)** Plots showing voom mean-variance trend with $\log_2$ (count size + 0.5) on the x-axis and square root of the standard deviation on the y-axis **(B)** MA plot showing fold-change ($\log_2$ scale) on the y-axis and mean of $\log_2$counts (x-axis). Genes differentially expressed at FDR < 0.05 are highlighted in red. **(C)** Distributions of uncorrected *P-values* and **(D)** FDR corrected *P-values*.

**Fig. S6.**

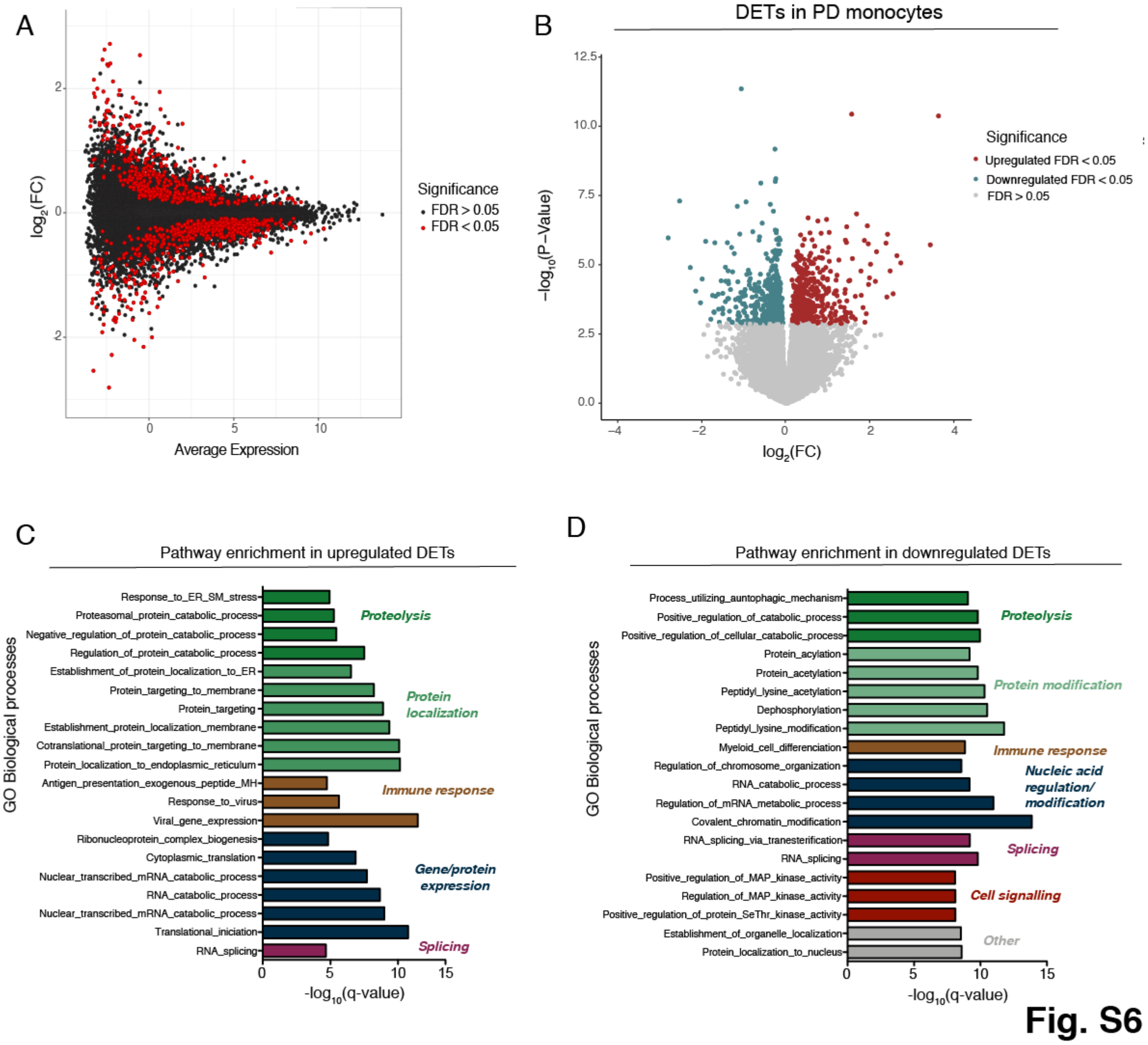


**Supplementary Figure 6. Differential expression analysis at the transcript level in PD and controls derived monocytes. (A)** MA plot showing the fold-change ($log_2$ scale) at the transcript level in the y-axis and the mean of $log_2$counts (x-axis), highlighting the DETs at FDR < 0.05 in red. **(B)** Volcano plot showing the fold-change ($log_2$ scale) of transcripts between PD-monocytes (n = 135) and controls (n = 95) (x-axis) and their significance in the y-axis -$log_{10}$ *P-value* scale). DETs at FDR < 0.05 are highlighted in red (upregulated) and blue (downregulated). Pathway enrichment analysis for the upregulated **(C)** and downregulated **(D)** DETs using Biological processes from GSEA. Significance is represented in the x-axis (-$log_{10}$ *P-value* scale of the q-value). Only the 20 most significant pathways (q-value < 0.05) with a minimum overlap of 5 genes are shown. Pathways are grouped and colored by biological related processes.

**Fig. S7.**

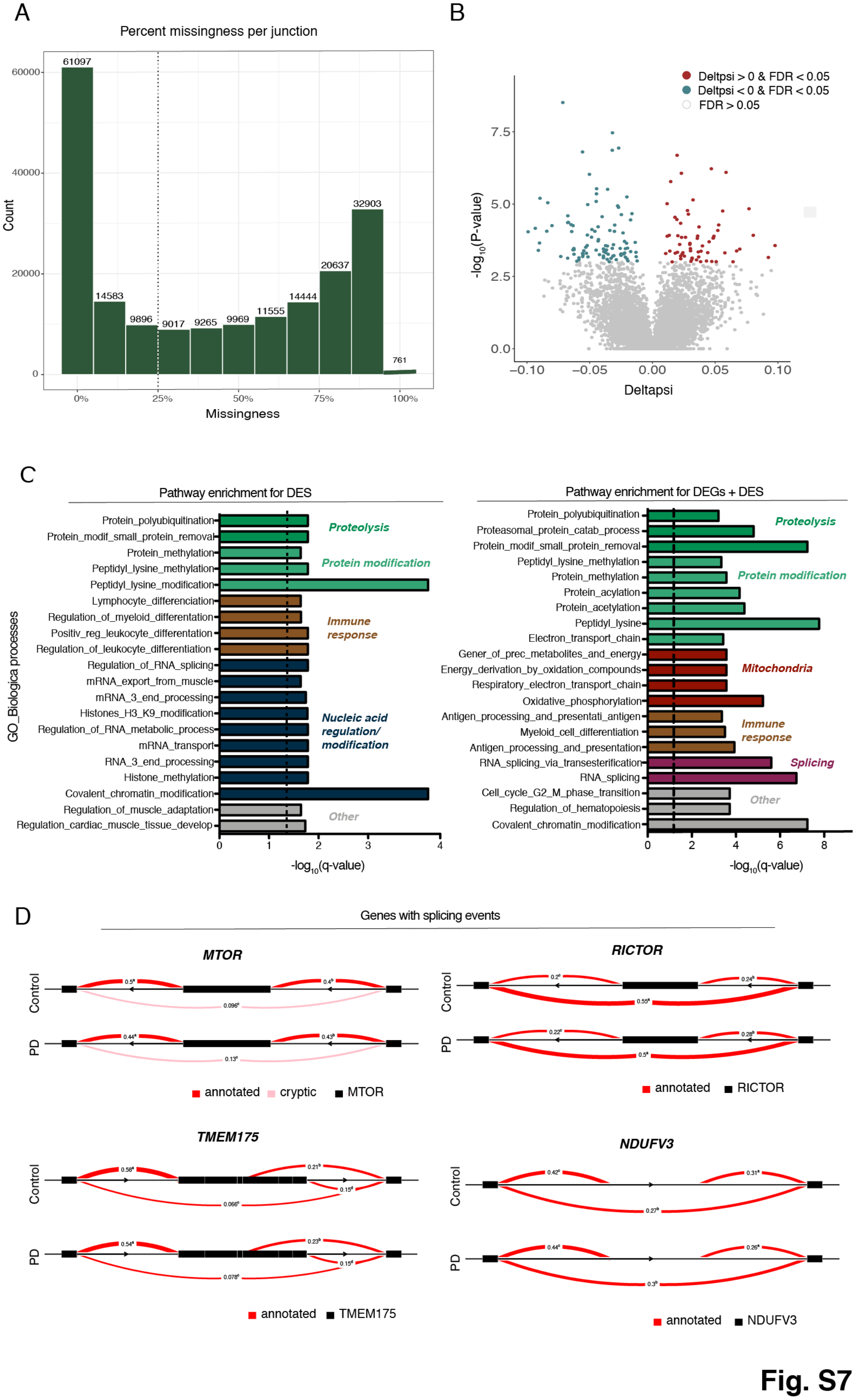
A
Percent missingness per junction
Count
Missingness
61097
14583
9896
9017
9265
9969
11555
14444
20637
32903
761
0%
25%
50%
75%
100%
B
Deltpsi > 0 & FDR < 0.05
Deltpsi < 0 & FDR < 0.05
FDR > 0.05
-log10(P-value)
Deltapsi
C
Pathway enrichment for DES
Pathway enrichment for DEGs + DES
GO_Biologica processes
-log10(q-value)
Proteolysis
Protein modification
Immune response
Nucleic acid regulation/ modification
Mitochondria
Splicing
Other
D
Genes with splicing events
MTOR
RICTOR
TMEM175
NDUFV3
Control
PD
annotated
cryptic
Fig. S7

**Supplementary Figure 7. Differential expression analysis at the splicing level in PD and controls derived monocytes. (A)** Histogram reflecting the counts (y-axis) and the % of missingness (x-axis). **(B)** Volcano plot showing the delta PSI of genes with splicing events in PD-monocytes (n = 135) and controls (n = 95) (x-axis) and their significance in the y-axis (-$\log_{10}$ *P-value* scale). DSs at FDR < 0.05 are highlighted in red (delta PSI > 0) and blue (delta PSI < 0). Positive delta-PSI indicates that the long isoform is favored whereas negative delta-PSI indicates preference for the short isoform. **(C)** Pathway enrichment analysis for the DSs at FDR < 0.0.5 (left panel) DSs + DEGs at FDR < 0.05 (right panel) using Biological processes from GSEA. Significance is represented in the x-axis (-$\log_{10}$ *P-value* scale of the q-value). Only the 20 most significant pathways (q-value < 0.05) with a minimum overlap of 5 genes are shown. Pathways are grouped and colored by biological related processes. **(D)** Examples of genes showing significant splicing events.

**Fig. S8.**

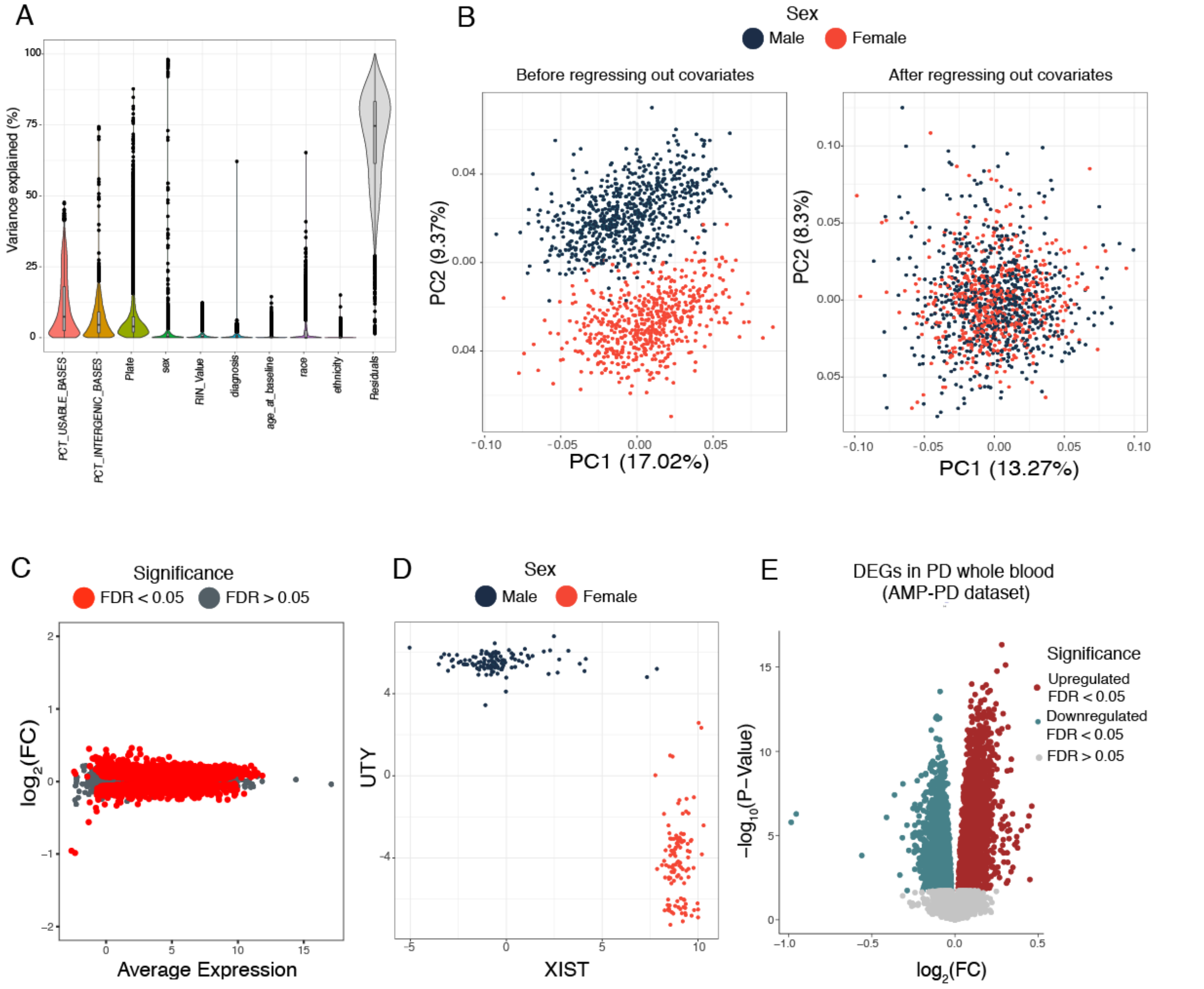


**Fig. S8**

**Supplementary Figure 8. RNA-seq QC of the whole blood transcriptomic analysis from AMP-PD dataset.** **(A)** Violin plot showing the % of variation (y-axis) explained by the covariates (x-axis). Each dot represents a gene. **(B)** Multidimensional reductionality using PCA of all the samples before (left panel) and after (right panel) regressing out the known covariates. Each dot represents a sample. Colored by sex (red = female, blue = male). **(C)** MA plot showing the fold-change ($log_2$ scale) of the DE cases vs controls in the y-axis and the mean of $log_2$counts (x-axis), highlighting the DEGs at FDR < 0.05 in red. **(D)** Sex mismatch QC: scatter plot showing the voom normalized expression of *XIST* (X-chromosome linked gene) in the x-axis and *UTY* (Y-chromosome linked gene) in the y-axis. Each dot represents a sample, colored by sex (red = female, blue = male). **(E)** Volcano plot showing the fold-change ($log_2$ scale) of PD-whole blood (n = 780) and controls (n = 504) (x-axis) and their significance in the y-axis ($-log_{10}$ scale). Upregulated DEGs at FDR < 0.05 are highlighted in red and downregulated DEGs in blue.

**Fig. S9.**

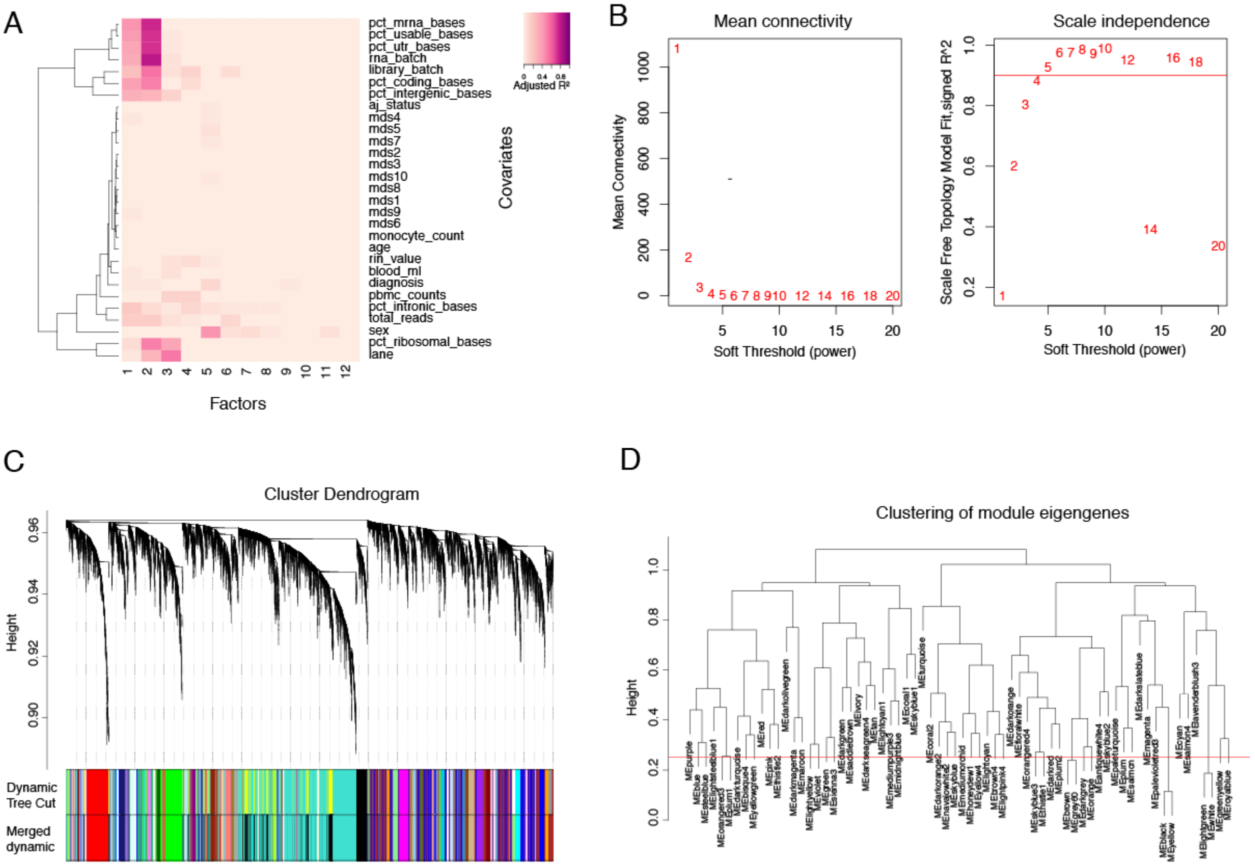


Fig. S9

**Supplementary Figure 9. Gene network construction in human monocytes using WGCNA.** Using the 230 monocyte samples, 65 co-expression modules were obtained using WGCNA **(A)** Heatmap showing the correlation of the first 12 SVs (x-axis) and the known covariates (y-axis). **(B)** Right: Evaluation of network topology with different soft-thresholding powers. The y-axis represents the scale-free fit index as a function of the soft-thresholding power (x-axis). Left: The mean connectivity (y-axis) as a function of the soft-thresholding power. **(C)** Gene dendrogram using “Dynamic Tree Cut” to assign genes to different modules and modules to colors, showing before and after collapsing modules into 65 final networks. **(D)** Module eigengenes clustering dendrogram based on topological overlap. Modules below the threshold (Module Dissimilarity = 0.25) indicated by the red line were merged. These values correspond to a correlation of 0.75.

Fig. S10.

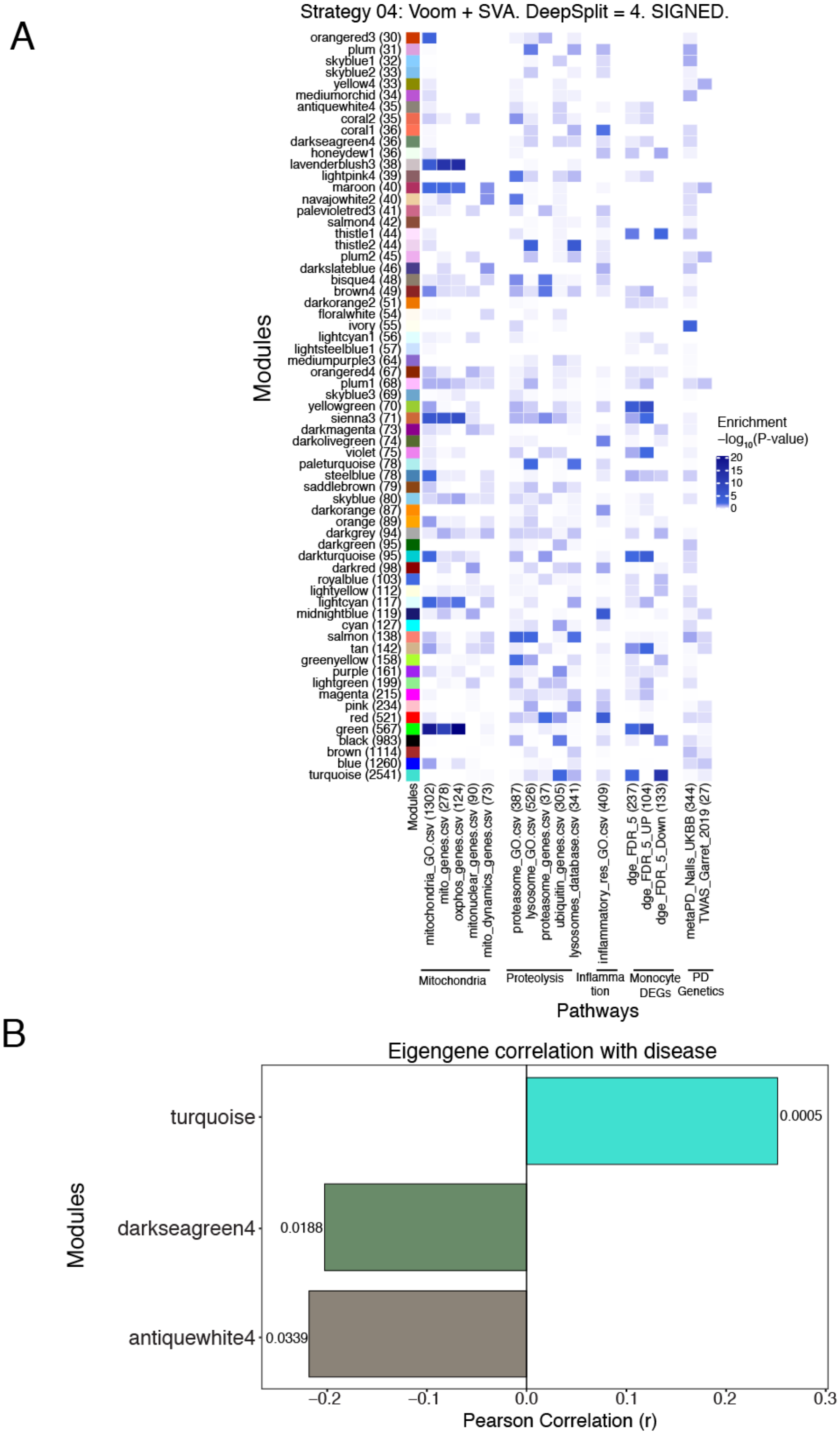


**Fig. S10**

**Supplementary Figure 10. Module enrichment for biological pathways. (A)** Heatmap showing the enrichment scores determined by Fisher exact test of the 65 modules (y-axis) for biological pathways, DEGs and PD-GWAS data (y-axis). **(B)** Barchart showing Pearson correlation coefficient (r) (x-axis) of three modules (y-axis) significantly associated with PD (FDR < 0.05) determined by the module eigengene analysis. Numbers on the plot represent adjusted *P-values,* Wilcoxon rank-signed test.

**Fig. S11.**

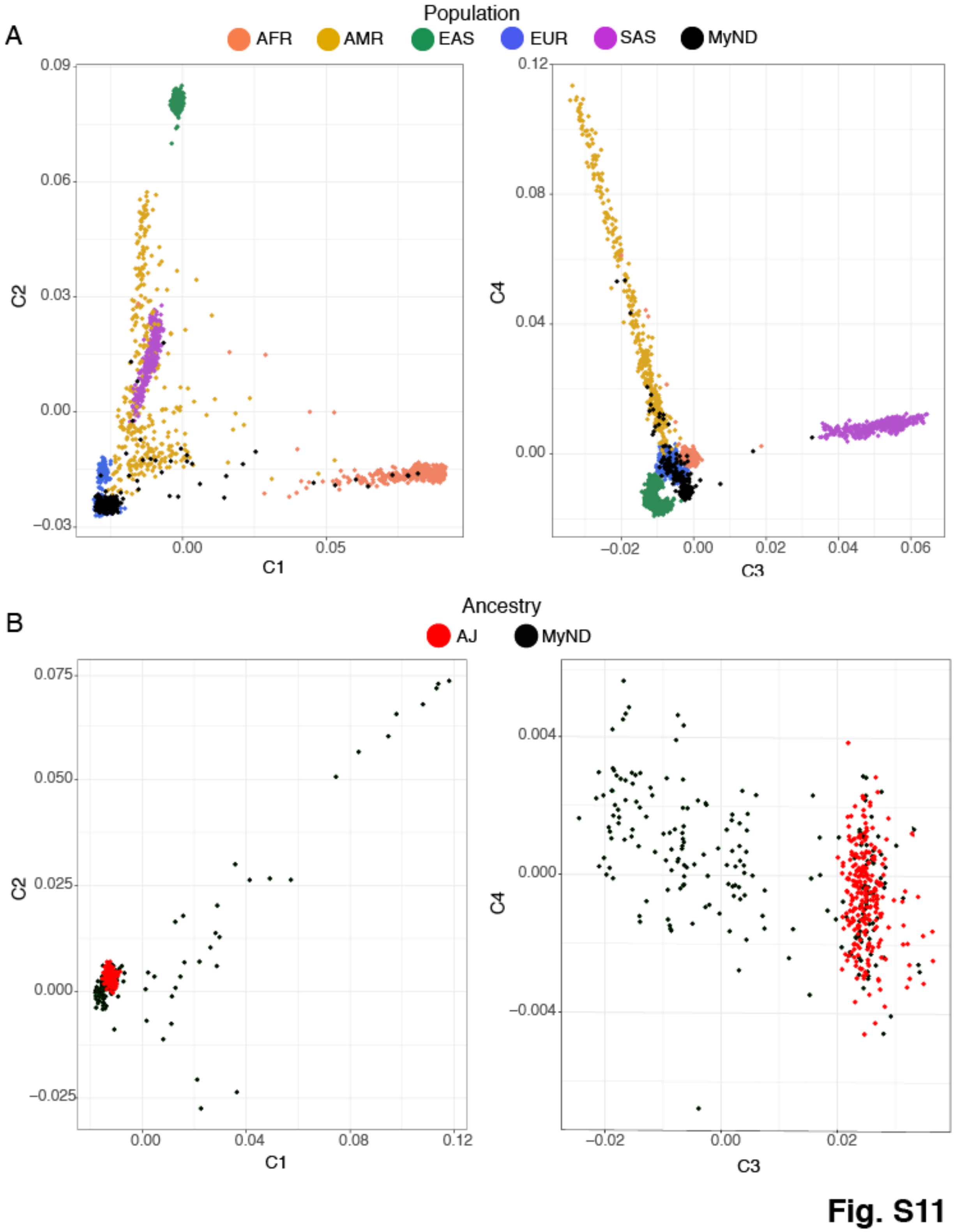
A
Population
AFR
AMR
EAS
EUR
SAS
MyND
C1
C2
C3
C4
B
Ancestry
AJ
MyND
Fig. S11

**Supplementary Figure 11. Analysis of population structure and ancestry estimates of subjects in MyND. (A)** Ancestry of each subject was confirmed by PCA using PLINK (Purcell et al., 2007 Am J Hum Genet) and multidimensional scaling (MDS) values of study subjects were compared to those of 1000 Genome Project samples (Phase 3). AFR = African, AMR = Admixed America, EAS = East Asian, EUR = European, SAS = South Asian. Samples of the MyND project are represented in black. **(B)** Multidimensional scaling values of study subjects were compared to those of Ashkenazi Jewish descent to determine ancestry.

**Fig. S12.**

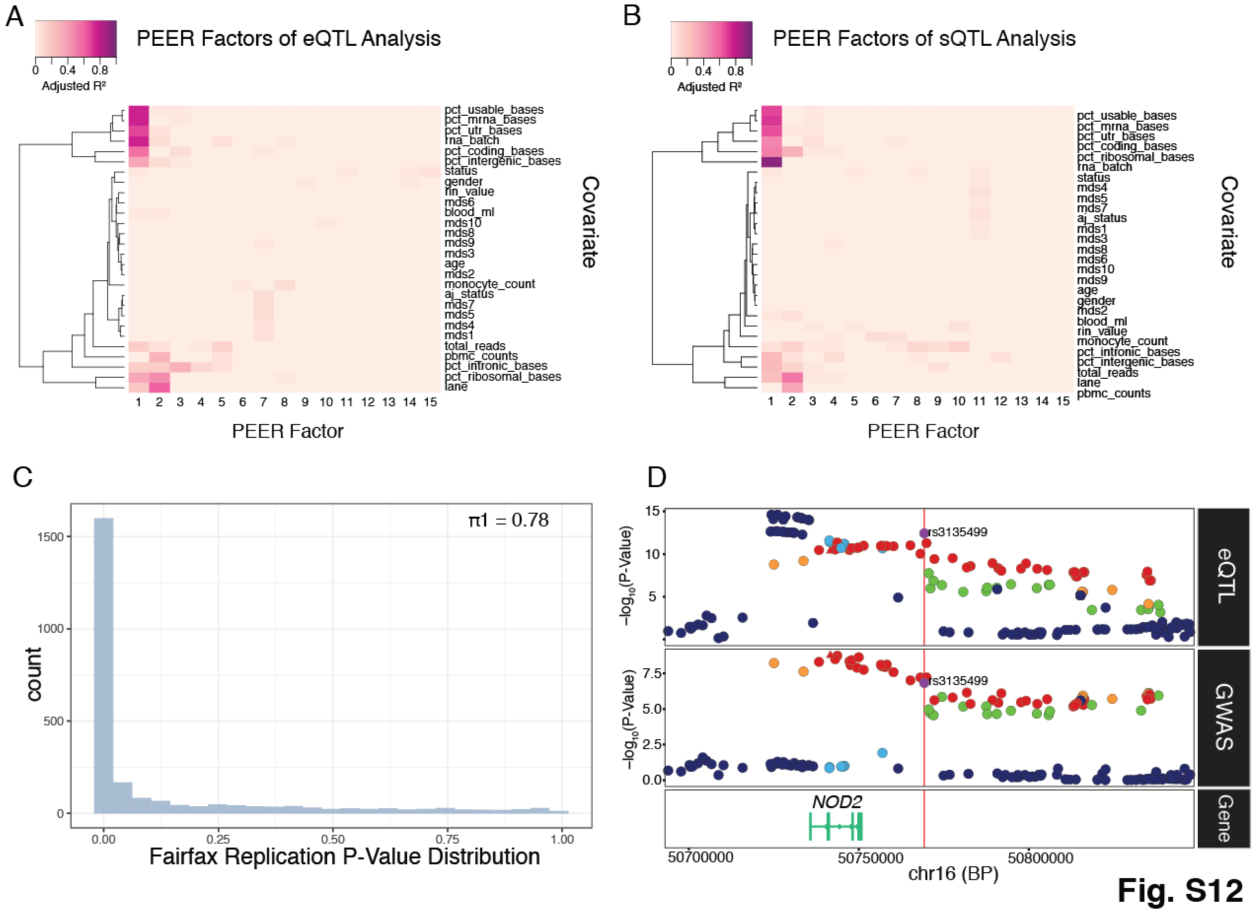


**Supplementary Figure 12. Using PEER to regress out technical confounders in eQTL and sQTL analysis.** Heatmap showing the correlation of the first 15 PEER factors (x-axis) and known covariates (y-axis) for the **(A)** eQTL analysis and **(B)** sQTL analysis**. (C)** Histogram showing replication of MyND eQTLs with Fairfax (*93*) eQTLs. **(D)** Regional association plot of monocyte eQTL (top panel) and PD GWAS (bottom panel) at the *NOD2* locus. The lead PD GWAS SNP rs6500328 is shown in red triangle. The lead eQTL SNP rs3135499 (in LD with rs6500328; $r^2 > 0.8$) is shown in purple.

**Fig. S13.**

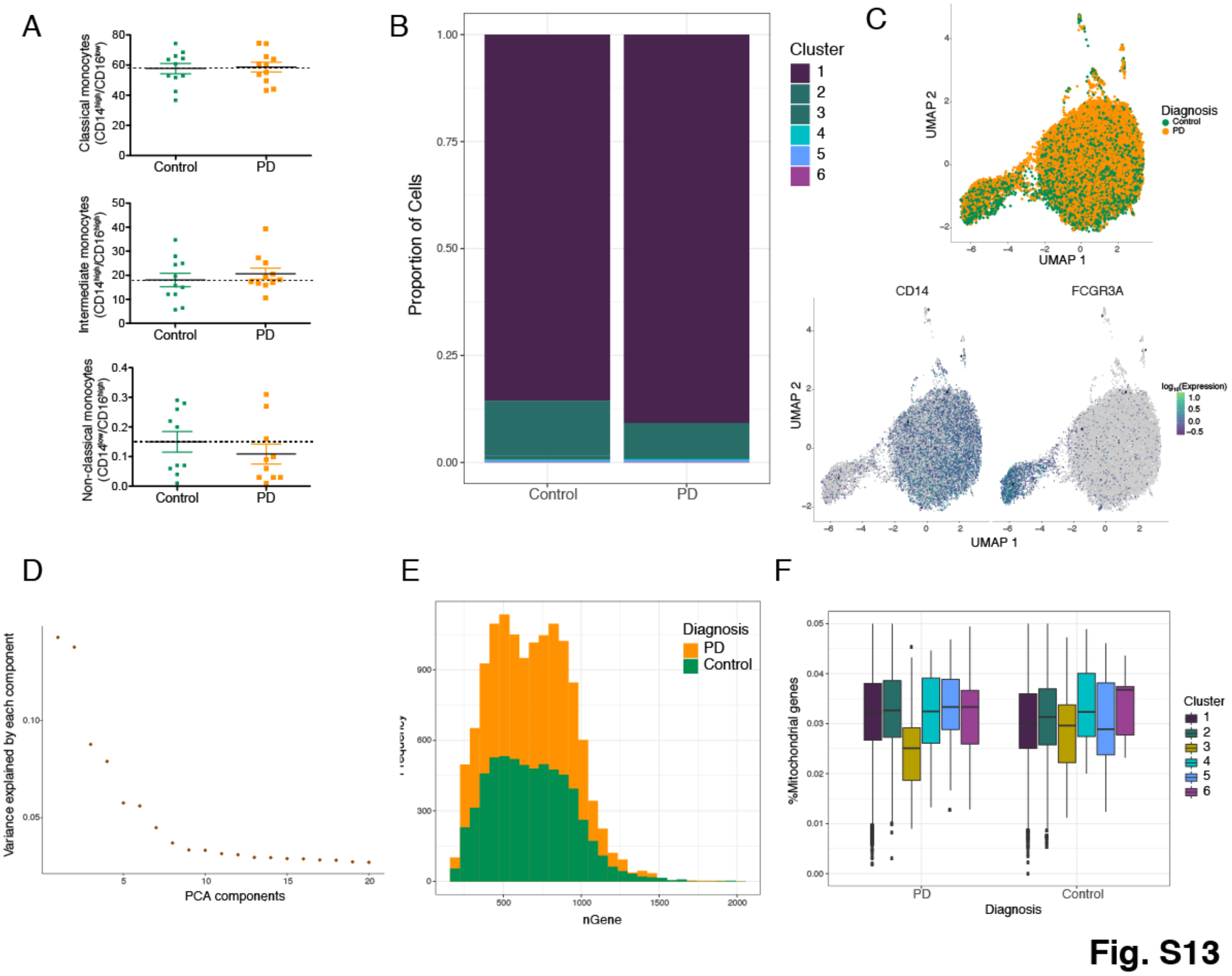


**Supplementary Figure 13. Monocyte sub-clusters characterization using single-cell RNA-seq. (A)** Proportions of the 3 main monocyte sub-clusters using FACS (n = 11 controls and 11 PD). No statistical differences were obtained between groups. **(B)** Cell proportions of the 6 sub-clusters obtained by unsupervised clustering with *monocle3* in scRNA-seq (n = 3 controls and 7 PD). No statistical differences were obtained between cases and controls in cell proportions. Cluster 1 corresponds to classical monocytes and cluster 2 to intermediate monocytes. **(C)** Top: UMAP colored by diagnosis (green = controls, yellow = PD). Bottom: UMAP colored by *CD14* and *FCGR3A* (CD16) marker genes expression. **(D)** Histogram showing the variance (y-axis) explained by the 20 first PCA components (x-axis). **(E)** Histogram showing the frequency of the genes colored by diagnosis (green = control, yellow = PD). **(F)** Expression of mitochondrial genes by each cluster and divided by diagnosis.

**Fig. S14.**

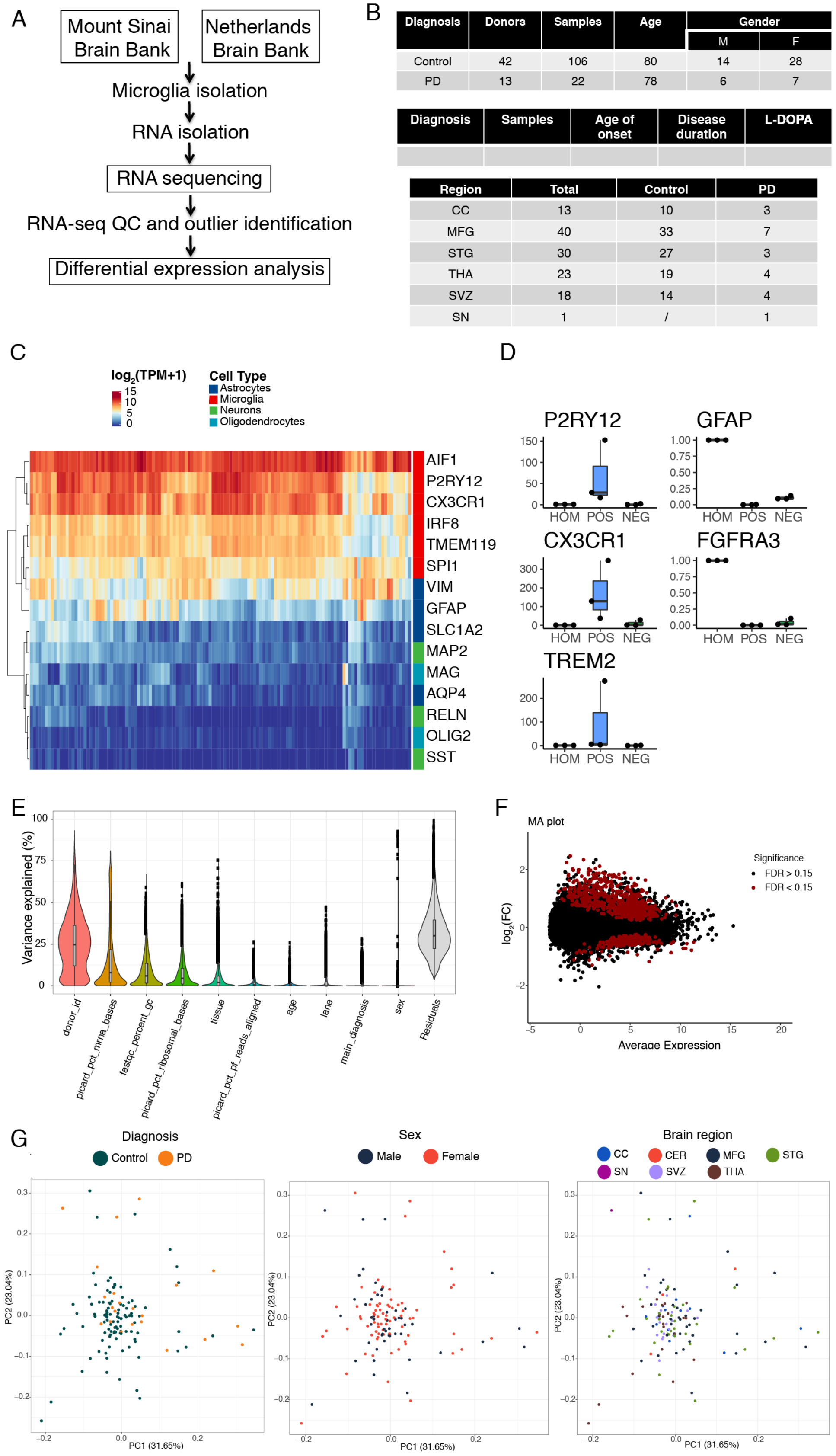




**Fig. S14**

**Supplementary Figure 14. Fresh microglia transcriptome analysis.** Microglia transcriptomic profiling was performed from 22 samples from 13 PD donors and 106 samples from 42 control donors. **(A)** Experimental workflow for the generation of human microglial transcriptomic profiles **(B)** Tables describing the samples included in the study (top: demographic information, middle: clinical information, bottom: brain regions). CC: Corpus Callosum; MFG: medial frontal gyrus; STG: Superior temporal gyrus; THA: thalamus; SVZ: subventricular zone; SN: substantia nigra **(C)** Heatmap for the expression of marker genes of different brain cell types (red: microglia, dark blue: astrocytes, green: neurons, light blue: oligodendrocytes). **(D)** Microglial isolation purity assessed by qPCR comparing the brain homogenate, and the positive and negative fractions after CD11b magnetic beads comparing microglial markers (*P2RY12*, *CXCR1*, *TREM2*) and astrocytic markers (*GFAP*, *FGFR3*). **(E)** Violin plot showing the % of variance (y-axis) explained by known covariates (x-axis) by variancePartition. Each dot represents a gene. **(F)** MA plot showing the fold-change ($log_2$ scale) of the DE cases vs controls (y-axis) and the mean of $log_2$ read counts (x-axis), highlighting the DEGs at FDR < 0.15 in red. **(G)** PCA after regressing out covariates colored by diagnosis (left panel), sex (middle panel), brain region (right panel).

**Table S1.**
Metadata for monocyte samples. This includes RNA-seq sample ID, the sample ID, ancestry (based on genetics), age, sample collection date, sex, and diagnosis

**Table S2**
Metadata for microglia samples. This includes tissue sample ID, donor ID, donor diagnosis, main diagnosis, tissue region, donor sex, donor age, post mortem delay, and tissue pH.

**Table S3**
Full summary statistics from monocyte differential gene expression between PD cases and control. This table includes $log_2$(FC), average expression, t-statistic, *P-value*, adjusted *P-value* and B (posterior probability of DE).

**Table S4**
Pathway analysis results for all monocyte differential gene expression analyses. This table includes gene set, number of genes in the set, description of the gene set, number of genes that overlap, ratio of overlapping genes with number of genes in the set, *P-value*, and adjusted *P-value*.

**Table S5**
Full summary statistics from monocyte differential transcript expression summary. This table includes $log_2$(FC), average expression, t-statistic, *P-value*, adjusted *P-value* and B (posterior probability of DE).

**Table S6**
Full summary statistics from monocyte differential splicing analysis. This table includes the cluster ID, the change in percent spliced in, number of junctions in that cluster**,** the genomics

coordinates of the cluster, the gene the cluster is in, the cluster annotation, the coordinates of the most significant intron junction, the log effect size, the estimated mean junction contribution from control subjects, the estimated mean junction contribution from PD subjects, the *P-value*, the adjusted *P-value*. Annotations are included only for significant clusters (FDR < 0.05).

**Table S7**
Pathway analysis results for all monocyte differential transcript expression analyses. Analysis was performed independently for upregulated and downregulated genes. This table includes gene set, number of genes in the set, description of the gene set, number of genes that overlap, ratio of overlapping genes with number of genes in the set, *P-value*, and adjusted *P-value*.

**Table S8**
Pathway analysis results for all monocyte differential splicing analyses. This table includes gene set, number of genes in the set, description of the gene set, number of genes that overlap, ratio of overlapping genes with number of genes in the set, *P-value*, and adjusted *P-value*.

**Table S9**
Full summary statistics from whole blood (AMP-PD) differential gene expression between PD cases and control. This table includes $\log_2$(FC), average expression, t-statistic, *P-value*, adjusted *P-value* and B (posterior probability of DE).

**Table S10**
Full summary statistics from differential gene expression from monocyte scRNA-seq across-clusters between PD cases and controls. This includes the gene name, estimate, the standard error, the Wald statistic, *P-value*, normalized effect size, and q-value.

**Table S11**
List of top 10 hub genes from WGCNA co-expression networks based on module membership values. This table includes the gene, the module to which the gene is a member, and the module membership score.

**Table S12**
Summary statistics for *cis*-eQTL permutation analysis between genes and the top variant in *cis* for all associations with FDR < 0.05. This table includes the Ensembl gene ID, the gene symbol, chromosome that the gene is located on, the gene start position, the gene position, strand orientation, number of variants tested in *cis,* distance between the lead variant and the gene, the ID of the lead variant, the chromosome of the lead variant, the variant start position, the variant end position, the number of degrees of freedom, a dummy column, the first parameter value of the fitted beta distribution, the second parameter value of the fitted beta distribution, the nominal *P-value*, the regression slope, the empirical *P-value*, the beta fitted *P-value*, and the FDR corrected beta fitted *P-value*.

**Table S13**
Summary statistics for *cis*-sQTL permutation analysis between genes and the top variant in *cis* for all associations with FDR < 0.05. This table includes the Ensembl gene ID, the gene symbol, chromosome that the gene is located on, the gene start position, the gene position, strand

orientation, the lead intron cluster, the number of clusters in the gene, the number of number of variants tested in *cis,* distance between the lead variant and the lead cluster, the ID of the lead variant, the chromosome of the lead variant, the variant start position, the variant end position, the number of degrees of freedom, a dummy column, the first parameter value of the fitted beta distribution, the second parameter value of the fitted beta distribution, the nominal *P-value*, the regression slope, the empirical *P-value*, the beta fitted *P-value*, and the FDR corrected beta fitted *P-value*, and the nominal *P-value* threshold per phenotype.

**Table S14**
Summary of PD GWAS and monocyte QTL (expression and splicing) colocalization analysis. The table includes the most recent PD GWAS reported locus, the lead SNP from GWAS in that locus, the colocalized QTL gene, the QTL lead SNP in LD with the lead GWAS SNP ($R^2 > 0.8$), the QTL lead SNP-Gene *P-value* and beta, the lead QTL reference allele, colocalization statistics for inclusion, and overlap of CD14 monocyte H3K27ac marks and microglia H3K27ac marks, ATAC-seq peaks, and PU.1 enhancer annotations.

**Table S15**
Metadata for monocyte scRNA-seq samples. This includes the sample ID, donor diagnosis, mutation status, reported ethnicity, donor sex, donor age.

**Table S16**
Full summary statistics from monocyte differential gene expression from scRNA-seq comparing intermediate vs classical cluster. This includes the gene name, estimate, the standard error, the Wald statistic, *P-value*, normalized effect size, and q-value.

**Table S17**
Full summary statistics from monocyte differential gene expression from scRNA-seq between PD cases and controls in the intermediate cluster. This includes the gene name, estimate, the standard error, the Wald statistic, *P-value*, normalized effect size, and q-value.

**Table S18**
Full summary statistics from microglia differential gene expression between PD cases and control. This table includes $log_2(FC)$, average expression, t-statistic, *P-value*, adjusted *P-value* and z.std (*P-value* transformed signed z-score).
